# Improving fine-mapping by modeling infinitesimal effects

**DOI:** 10.1101/2022.10.21.513123

**Authors:** Ran Cui, Roy A Elzur, Masahiro Kanai, Jacob C Ulirsch, Omer Weissbrod, Mark J Daly, Benjamin M Neale, Zhou Fan, Hilary K Finucane

**Affiliations:** Analytic and Translational Genetics Unit, Massachusetts General Hospital, Boston, MA, USA; Program in Medical and Population Genetics, Broad Institute of MIT and Harvard, Cambridge, MA, USA; Stanley Center for Psychiatric Research, Broad Institute of MIT and Harvard, Cambridge, MA, USA; The Novo Nordisk Foundation Center for Genomic Mechanisms of Disease, Broad Institute of MIT and Harvard, Cambridge, MA, USA; Department of Biomedical Informatics, Harvard Medical School, Boston, MA, USA; Institute for Molecular Medicine Finland (FIMM), University of Helsinki, Helsinki, Finland; Department of Statistical Genetics, Osaka University Graduate School of Medicine, Suita, Japan; Program in Biological and Biomedical Sciences, Harvard Medical School, Boston, MA, USA; Department of Epidemiology, Harvard T.H. Chan School of Public Health, Boston, MA, USA; Department of Statistics and Data Science, Yale University, New Haven, CT, USA

## Abstract

Fine-mapping aims to identify causal variants for phenotypes. Bayesian fine-mapping algorithms (e.g.: SuSiE, FINEMAP, ABF, and COJO-ABF) are widely used, but assessing posterior probability calibration remains challenging in real data, where model misspecification likely exists, and true causal variants are unknown. We introduce Replication Failure Rate (RFR), a metric to assess fine-mapping consistency by down-sampling. SuSiE, FINEMAP and COJO-ABF show high RFR, indicating potential under-conservative mis-calibration. Simulations reveal that non-sparse genetic architecture can lead to miscalibration, while imputation noise, non-uniform distribution of causal variants, and QC filters have minimal impact. We present SuSiE-inf and FINEMAP-inf, novel fine-mapping methods modeling infinitesimal effects alongside fewer larger causal effects. Our methods exhibit improved calibration, RFR and functional enrichment, competitive recall and computational efficiency. Notably, using our methods’ posterior effect sizes substantially increases PRS accuracy over SuSiE and FINEMAP. Our work improves causal variants identification for complex traits, a fundamental goal of human genetics.

## Introduction

Over the past two decades, genome-wide association studies (GWAS) have successfully identified thousands of loci that are associated with various diseases and traits^1^. However, refining these associations to identify causal variants remains challenging, due to extensive linkage disequilibrium (LD) among associated variants^2^. Many approaches can be taken to help nominate causal variants from associations, such as overlapping GWAS signals with coding or functional elements of the genome^3^, with eQTLs^4^, and across populations having different ancestries and patterns of LD^5–7^. Complementary to and in conjunction with these approaches, Bayesian sparse regression and variable selection methods, which aim to identify causal variants and quantify their uncertainty based upon a statistical model (e.g.: SuSiE^8^, FINEMAP^9,10^, ABF^11^, and COJO^12^-ABF) are widely applied in practice^13–19^.

The appeal of Bayesian approaches to fine-mapping is two-fold. First, these methods determine a posterior inclusion probability (PIP) for each variant, quantifying the probability that the variant is causal under the model, which can reflect uncertainty due to LD. For example, two variants in strong LD and harboring a strong association with the phenotype may each have PIP close to 50%, representing confidence that there is a causal signal but uncertainty about which variant(s) is/are causal. Second, these methods incorporate assumptions about genetic architecture -- namely, the relative probabilities of different numbers of and configurations of causal SNPs, as reflected by a Bayesian prior -- to improve statistical power for identifying high-confidence variants.

Bayesian fine-mapping methods are correctly calibrated when the PIPs accurately reflect the true proportions of causal variants, e.g., 9 out of 10 variants having PIP 90% are truly causal for the trait. Calibration (i.e. whether or not the posterior probability of causality reflects the true proportion of causal variants) is ensured when the linear model for genetic effects and Bayesian prior for genetic architecture across loci are both correctly specified, and accurate calibration has also been demonstrated empirically in simulations to be robust under mild model misspecifications^20^. However, the actual calibration and false discovery rates of these methods in real data applications are not easily determined, as true causal variants and the sources of model misspecification may be unknown.

Here, we propose the Replication Failure Rate (RFR) to assess the stability of fine-mapping methods by evaluating the consistency of PIPs in random subsamples of individuals from a larger well-powered cohort. We found the RFR to be higher than expected across traits for several Bayesian fine-mapping methods. Moreover, variants that failed to replicate at the higher sample size were less likely to be coding. Together these analyses suggest that SuSiE, FINEMAP, and COJO-ABF may be mis-calibrated in real data applications. In other words, they may return a disproportionately large number of false discoveries among high-PIP variants.

We performed large-scale simulations to assess the effects of several plausible sources of model misspecification on calibration. These simulations — which include, among other factors, varying levels of non-sparsity and stratification — suggest that a denser and more polygenic architecture of genetic effects may be a major contributor to PIP mis-calibration. We thus propose incorporating a model of infinitesimal effects when performing Bayesian sparse fine-mapping, recasting the goal of fine-mapping as the identification of a sparse set of large-effect causal variants among many variants having smaller effects. We develop and implement fine-mapping tools SuSiE-inf and FINEMAP-inf that extend the computational ideas of SuSiE and FINEMAP to model additional infinitesimal genetic effects within each fine-mapped locus.

Applying SuSiE-inf and FINEMAP-inf to 10 quantitative traits in the UK Biobank shows improved RFR. SuSiE-inf high-PIP variants are more functionally enriched than SuSiE high-PIP variants. Cross-ancestry phenotype prediction using SuSiE-inf/FINEMAP-inf shows significant improvement over SuSiE/FINEMAP across 7 traits and 6 diverse ancestries. These results suggest that explicit modeling of a polygenic genetic architecture, even within individual genome-wide significant loci, may substantially improve fine-mapping accuracy.

## Results

### Real data benchmarking shows that current fine-mapping methods are likely mis-calibrated

Real-data benchmarking of fine-mapping methods is challenging due to the lack of ground truth. However, down-sampling large cohorts allows assessment of the methods’ stability. We chose 10 well-powered quantitative phenotypes (**Online Methods**) in the UK Biobank and computed the Replication Failure Rate (RFR) for SuSiE, FINEMAP as follows (see **Supplementary Note** for results related to ABF and COJO-ABF). Our group previously performed fine-mapping^20^ on a cohort of 366,194 unrelated “white British” individuals defined in the Neale Lab UKBB GWAS^21^. We down-sampled this cohort to a random subsample of 100,000 and performed fine-mapping with the same pipeline (**Online Methods**). RFR is defined as the proportion of high-confidence (PIP > 0.9) variants fine-mapped in the 100K subsample that failed to replicate (PIP < 0.1) in the full 366K cohort. This RFR is an estimate of the conditional probability *Pr*(*PIP*^366K^ < 0.1 | *PIP*^100K^ > 0.9) for a randomly chosen variant. In a truly sparse causal model, assuming that the method is well-powered at sample size N=366K to detect true causal variants which are identified with high confidence at 100K, the RFR is an approximate lower bound for the false discovery rate *Pr*(*not causal* | *PIP*^100K^ > 0.9) (see **Supplementary Note**).

Across all 10 traits, we observed different levels of RFR for different phenotypes, and an aggregated RFR of 15% for SuSiE and 12% for FINEMAP (**Fig. 1a-b**; see **Extended Data Fig. 1** for other PIP thresholds). These values far exceed the false discovery rate expected in a correctly specified sparse Bayesian model (SuSiE 1.8%, FINEMAP 2.0%), which we denote by EPN (Expected Proportion of Non-causal variants) and estimate from the mean reported PIPs exceeding 0.9. In contrast, ideal simulations under correctly specified models show close agreement between RFR and EPN (**Fig. 1a, Online Methods**, and **Supplementary Note**).

**Fig. 1.**
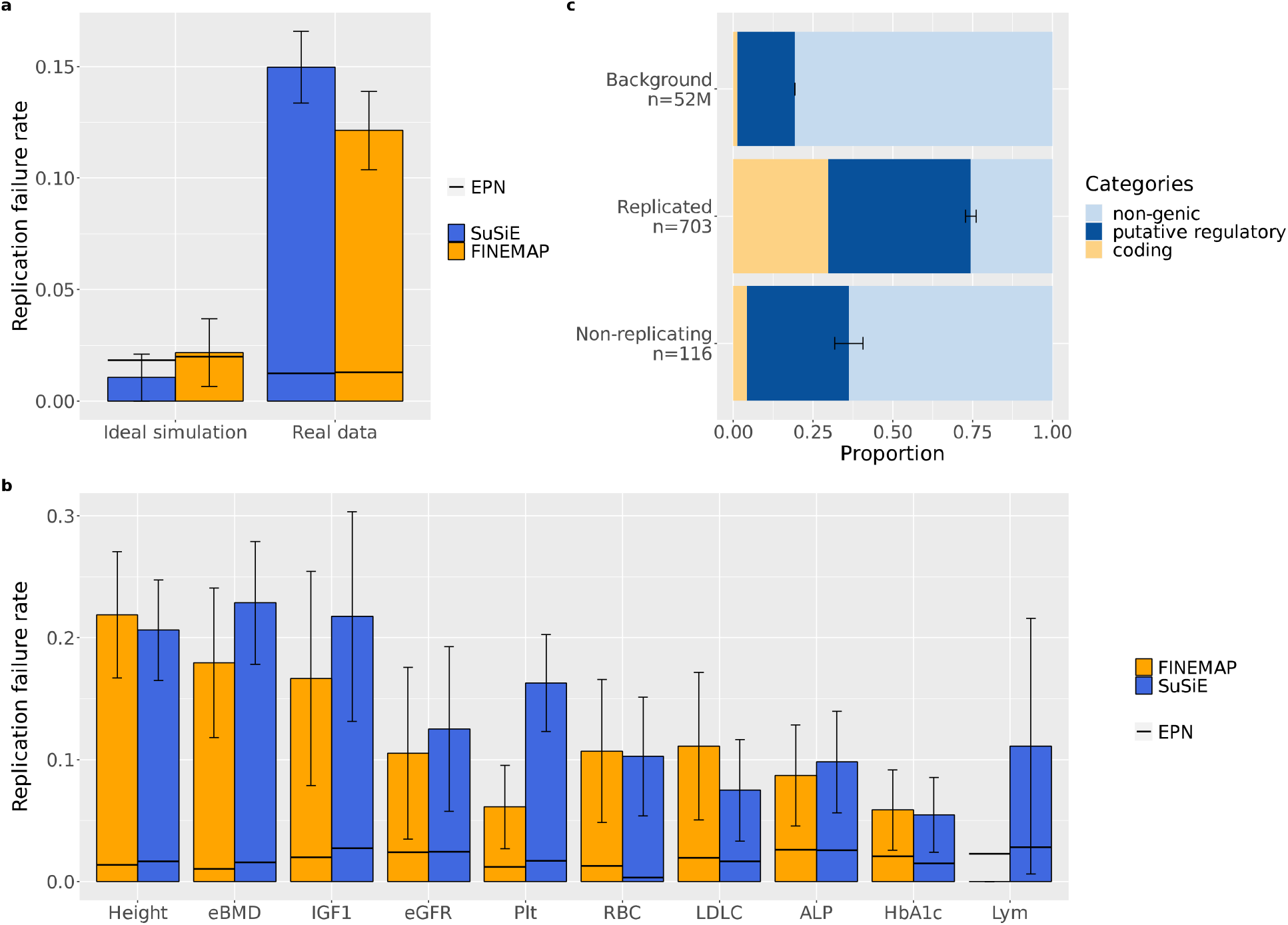
Replication failure rates and functional enrichments. **a**. RFRs for SuSiE and FINEMAP aggregated across 10 UKBB quantitative phenotypes, contrasted with ideal simulations and with Expected Proportion of Non-causal variants (EPN). **b**. Trait-separated RFRs for SuSiE and FINEMAP. **c**. Functional annotations in 3 disjoint categories: coding, putative regulatory and non-genic (see **Online Methods** for detailed definitions). Variants are aggregated between SuSiE and FINEMAP. Non-replicating: the set of non-replicating variants (PIP>0.9 at N=100K and PIP<0.1 at N=366K); Replicated: the set of replicated variants (PIP>0.9 at both N=100K and N=366K); Background: the set of all variants included in the fine-mapping analysis, aggregated across 10 traits. n denotes the total number of variants in each set. See **Supplementary Fig. 1** for method-separated plots and more sets of variants. Error bars represent one standard deviation of the corresponding binomial distribution. Numerical results are available in **Supplementary Table 4**,**5**.

Across all 10 traits, we observed varying levels of RFR, with an aggregated RFR of 15% for SuSiE and 12% for FINEMAP (**Fig. 1a-b**; see **Extended Data Fig. 1** for other PIP thresholds). These values far exceed the false discovery rate expected in a correctly specified sparse Bayesian model (SuSiE 1.8%, FINEMAP 2.0%), which we denote by EPN (Expected Proportion of Non-causal variants), estimated from the mean reported PIPs exceeding 0.9. In contrast, ideal simulations under correctly specified models show close agreement between RFR and EPN (**Fig. 1a, Online Methods**, and **Supplementary Note**).

To gain insight into whether non-replicating variants (PIP>0.9 at 100K and PIP<0.1 at 366K) are causal, we examined the functional annotations, focusing on two distinct categories: coding and putative regulatory (**Online Methods**). We found a significant depletion of functionally important variants in the non-replicating set compared to the replicated set (P=1.7e-7) (**Fig. 1b, Online Methods**). This suggests that many non-replicating variants may be non-causal, and that SuSiE and FINEMAP may be miscalibrated when applied in real data (see **Online Methods** for our investigation into other potential causes for high RFR). We found higher functional enrichment in the set of non-replicating variants than the background, suggesting some PIPs at N=366K may be too conservative. However, here we focus on investigating the more concerning under-conservative PIPs which can lead to elevated false discovery rate.

### Un-modeled non-sparse causal effects can lead to miscalibration

Bayesian sparse variable selection approaches to fine-mapping, including SuSiE and FINEMAP, commonly rely on some of the following assumptions: (1) Within each genome-wide significant locus, one or a small number of variants have a true causal contribution to the phenotype. (2) All true causal variants within the locus are included in, or tagged by a sparse subset of, the analyzed genotypes. (3) The distribution of causal variant effect sizes is well-approximated by a simple, oftentimes Gaussian, prior. (4) There is no uncorrected confounding, and the residual error is uncorrelated with the genotype. (5) There is no imputation noise or error in the genotypes. Violations of any of these assumptions can, in principle, cause miscalibration, although the severity of such miscalibration under the degrees of violation that are present in fine-mapping applications is unclear a priori.

We designed large-scale simulations to investigate how SuSiE and FINEMAP may be affected by these five sources of misspecification. Our simulations use UK Biobank genotypes (N=149,630 individuals of white British ancestry) and BOLT-LMM^22^ for GWAS, incorporating (1) varying amounts of unmodeled non-sparse causal effects (varying both the coverage of non-sparsity, i.e., the proportion of variants with non-zero effects, and the amount of heritability the non-sparse component explains), (2) missing causal variants that are removed by quality-control filtering prior to fine-mapping, (3) effect size distributions for the large and sparse causal variants that reflect estimates from fine-mapping of real traits, (4) varying amounts of uncorrected population stratification, and (5) imputation noise in the input genotypes (see **Online Methods** for detailed description of our simulations and other misspecifications we considered). In previous work^20^ by our group, we found that quality control filters and imputation noise did not contribute to miscalibration in simulations; here we continued to include them while adding non-sparsity, effect size estimates from real data, and uncorrected population stratification as additional sources of miscalibration. Note that we simulated a single cohort, without the heterogeneity that often comes with meta-analysis where quality control and imputation are important contributors to miscalibration^23^. Moreover, we did not consider error in the probabilities outputted by standard imputation software or different types of genotyping error, which could contribute to miscalibration even in the absence of heterogeneity.

Our simulations show that missing causal variants due to QC, use of a realistic non-Gaussian effect size distribution estimated from real data, and imputation error did not induce miscalibration, consistent with and extending previous results^20^.

SuSiE and FINEMAP were both miscalibrated in simulations with non-sparse effects. For example, when non-sparse causal effects explain 75% of the total SNP-heritability, only about 80% of variants with PIP ≥ 0.9 are causal, far below the rate of approximately 97% that we would expect given the variants’ mean PIP. Miscalibration increased and recall decreased as we increased the proportion of total SNP-heritability (set at 0.5; for comparison, see **Supplementary Table 1** for common-SNP heritability in real traits) explained by non-sparse effects from 58% to 100% (**Fig 2a-b, Table 1**) while fixing the coverage to 1%. This trend was consistently observed at different levels of coverage, see **Online Methods, Extended Data Fig. 2** for results at 0.5% and 5% coverage. We emphasize that calibration was measured against the set of all causal variants, including the non-sparse causal effects.

**Fig. 2.**
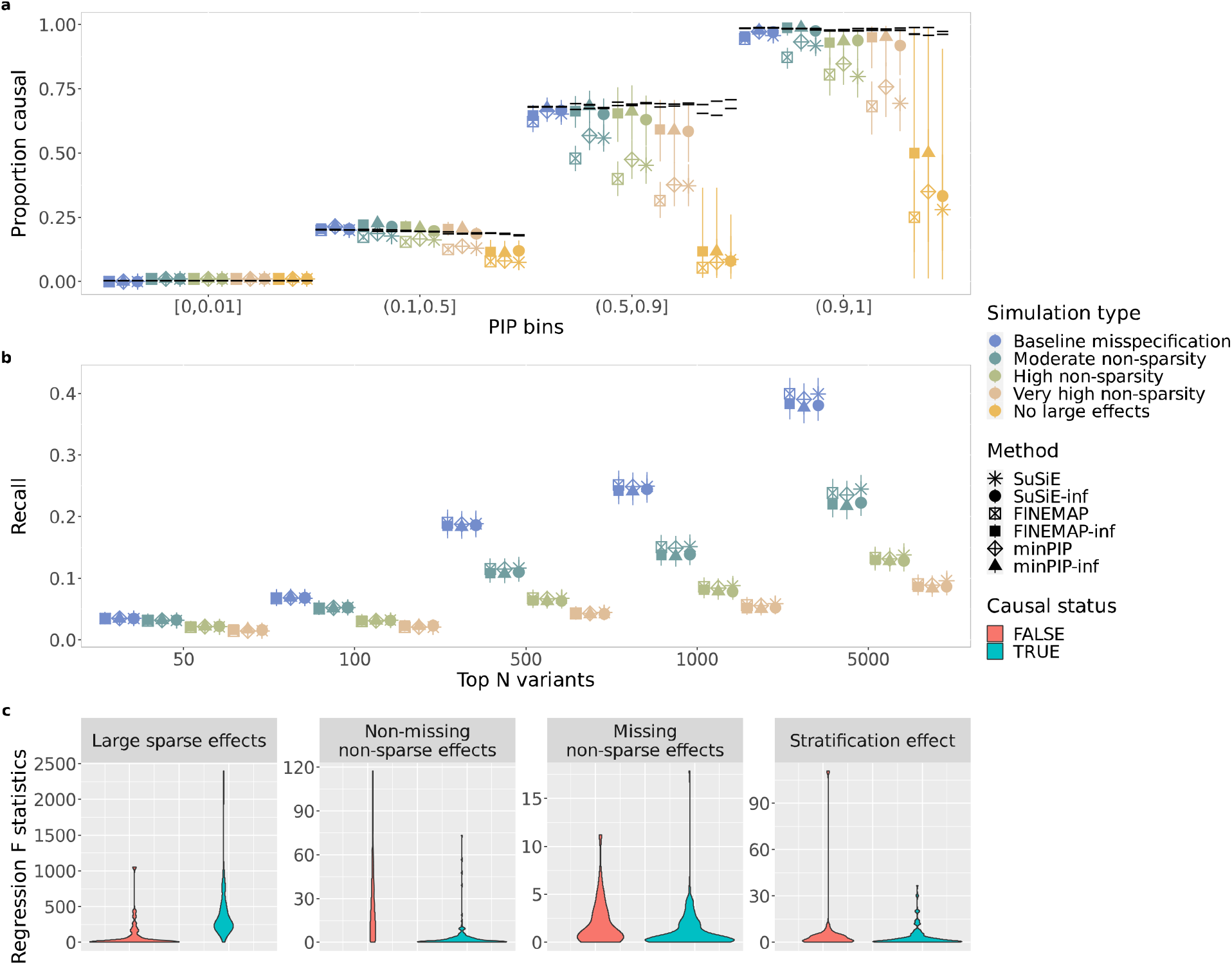
non-sparsity simulation. **a**. Calibration for SuSiE, FINEMAP, minPIP, and corresponding “inf” methods under non-sparsity simulation settings detailed in **Table 1, Online Methods**. minPIP and minPIP-inf are aggregating methods: minPIP-inf = min(PIP) between SuSiE-inf and FINEMAP-inf; minPIP = min(PIP) between SuSiE and FINEMAP. **b**. Recall for these same methods, defined as the percentage of simulated large effects among the top N variants when ranked by PIP. Error bars on calibration and recall plots correspond to 95% Wilson confidence interval. Note that “No large effects” simulations are not shown on the recall plot because there are zero simulated large effects. **c**. Regressing sub-components of “high non-sparsity” phenotype on true vs. false positives (variants with PIP > 0.9 that are either causal or non-causal). Numerical results are available in **Supplementary Table 6-8**.

**Table 1.** Parameters for selected large-scale simulations. Different parameter settings for ten sets of simulations mentioned in the main text. Note that PCs corrected in GWAS used in-sample (N=150K) PCs as covariates for phenotypes generated with full sample (N=366K) PCs. See **Online Methods** for details on how each misspecification is incorporated.

|  | Imputation noise | Sparse causal prior | 20 PC effects multiplier | PCs corrected in GWAS | Non-sparse causal effects | Missing causal effects |
| --- | --- | --- | --- | --- | --- | --- |
| Ideal | No | Uniform | 0 | 0 | None | None |
| Baseline misspecification | Yes | SuSiE Height posterior | 1 | 19 out of 20 | None | None |
| Moderate stratification w/ BOLT | Yes | SuSiE Height posterior | 5 | 0 out of 20 | None | None |
| Severe stratification w/ BOLT | Yes | SuSiE Height posterior | 8 | 0 out of 20 | None | None |
| Moderate stratification w/ OLS | Yes | SuSiE Height posterior | 1 | 0 out of 20 | None | None |
| Severe stratification w/ OLS | Yes | SuSiE Height posterior | 2 | 0 out of 20 | None | None |
| Moderate non-sparsity | Yes | SuSiE Height posterior | 1 | 19 out of 20 | 58% of h2, 1% coverage | Yes |
| High non-sparsity | Yes | SuSiE Height posterior | 1 | 19 out of 20 | 75% of h2, 1% coverage | Yes |
| Very high non-sparsity | Yes | SuSiE Height posterior | 1 | 19 out of 20 | 83% of h2, 1% coverage | Yes |
| No large effects | Yes | SuSiE Height posterior | 1 | 19 out of 20 | 100% of h2, 1% coverage | Yes |
| 0.5% coverage, ratio 3:1 | Yes | SuSiE Height posterior | 1 | 19 out of 20 | 75% of h2, 0.5% coverage | Yes |
| 5% coverage, ratio 3:1 | Yes | SuSiE Height posterior | 1 | 19 out of 20 | 75% of h2, 5% coverage | Yes |
| 5% coverage, ratio 15:1 | Yes | SuSiE Height posterior | 1 | 19 out of 20 | 94% of h2, 5% coverage | Yes |

To further support that unmodeled non-sparse causal effects, among all the misspecification we incorporated, formed the primary driver of the observed miscalibration, we decomposed the simulated genetic component *Xβ* of the phenotype into the sum of four sub-components representing sparse causal effects, missing causal variants, uncorrected stratification, and unmodeled non-sparse causal effects. Regressing each of these four sub-components on the true and false positive variants (respectively defined as causal and non-causal variants with PIP ≥ 0.9), false positive variants were significantly more correlated with the non-sparse causal effects than true positive variants (**Fig. 2c, Online Methods**).

Our simulated population stratification under standard pipeline (**Online Methods**) where BOLT-LMM was used for association mapping failed to induce miscalibration. Replacing BOLT-LMM with ordinary least squares (OLS) for association mapping allowed us to induce higher levels of uncorrected confounding (**Supplementary Table 2**) that did lead to miscalibration (**Fig. 3**) but are less true to the pipeline used in our real data applications. See **Online Methods** for interpretation and more discussion on these results.

**Fig. 3.**
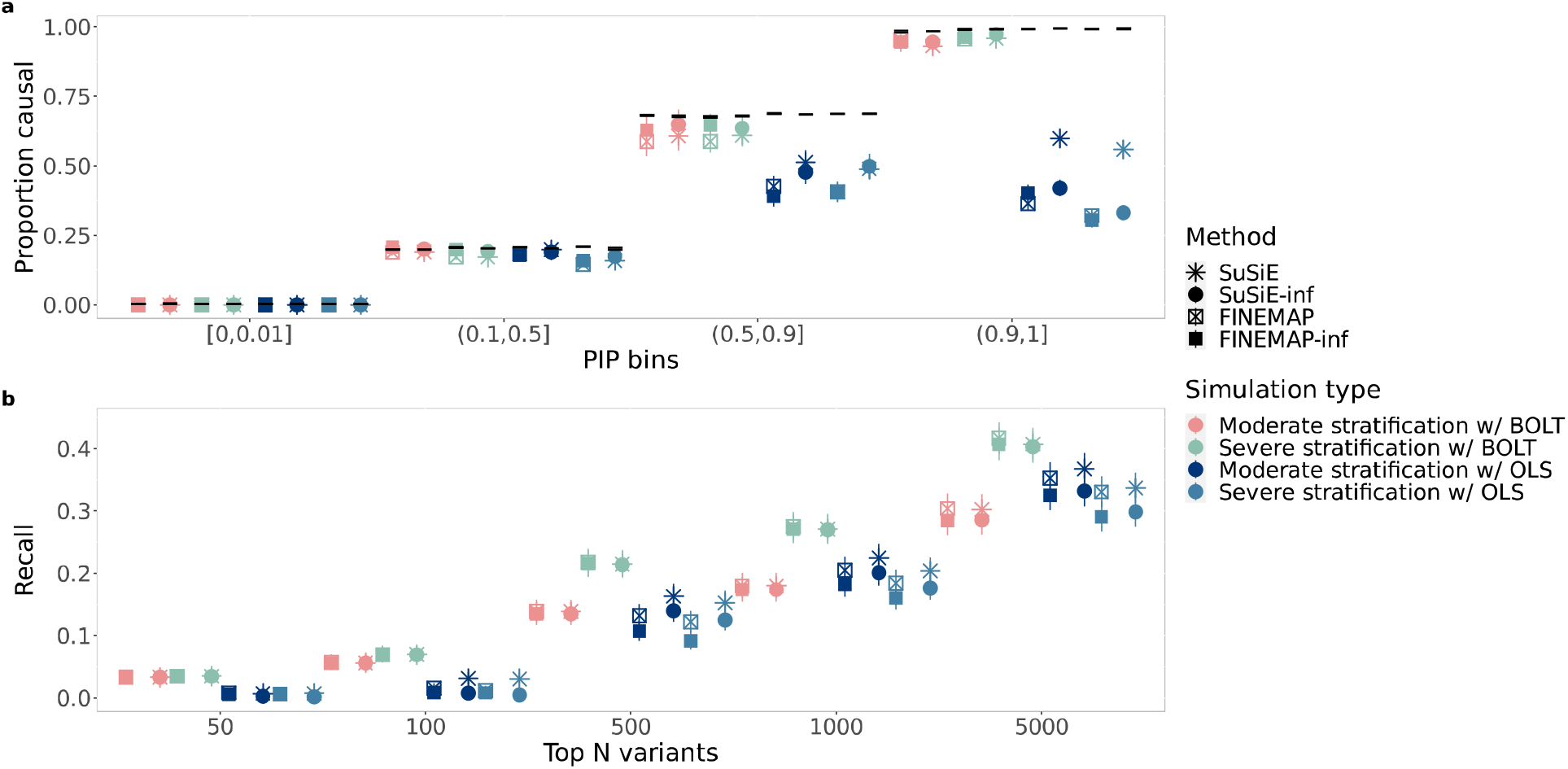
Calibration and recall for stratification simulation. **a**. Calibration plot for six methods in four stratification simulation settings (**Table 1**). **b**. Recall for the same methods and simulations. Numerical results are available in **Supplementary Table 6-7**.

In conclusion, the presence of non-sparse effects is a driver of miscalibration for SuSiE and FINEMAP. The stratification we simulated only induced miscalibration when using OLS for association mapping but not when using BOLT-LMM. None of the other sources of misspecification incorporated in our simulations caused miscalibration within our fine-mapping pipeline.

### New methods for Bayesian fine-mapping

To address PIP miscalibration that may arise from non-sparse causal effects, we propose to explicitly incorporate a model of broad infinitesimal genetic effects when fine-mapping causal variants. Here, we describe two specific implementations of this idea that extend FINEMAP and SuSiE. We call the resulting methods FINEMAP-inf and SuSiE-inf.

FINEMAP-inf and SuSiE-inf are based on a random-effects linear model *y* = *X*(*β* + *α*) + *ϵ* for observed phenotypes *y* across *n* samples, where X is a *n* by *p* genotype matrix for *p* variants, *β* is a vector of sparse genetic effects of interest, *α* is an additional vector of dense infinitesimal effects, and *ϵ* is residual error. In the context of such a model, we define the primary goal of fine-mapping as inferring the non-zero coordinates of the sparse component *β*. We will refer to these coordinates as the “causal model” and the “causal variants”, although in this model, every variant may have an additional small causal effect on y through the infinitesimal component *α*.

We model coordinates of *α* and of the residual error *ϵ* as independent and identically distributed (i.i.d.) with normal distributions 𝒩(0,*τ*^2^) and 𝒩(0,*σ*^2^) respectively, where *τ*^2^ is the effect size variance for the infinitesimal effect. For FINEMAP-inf, coordinates of the sparse effects *β* are also modeled as i.i.d., with point-normal distribution *π*_0_𝒩(0, *s*^2^) + (1−*π*_0_)*δ*_0_. We use a shotgun-stochastic-search (SSS) procedure as in FINEMAP for performing approximate posterior inference of the sparse component *β*, marginalizing its posterior distribution over both the infinitesimal effects *α* and the residual errors *ϵ*. The SSS is divided into several epochs, and we propose a method-of-moments approach to update estimates of the variance components (*σ*^2^,*τ*^2^) between epochs.

For SuSiE-inf, we follow the approach of SuSiE and instead parametrize the sparse causal effects as a sum of single effects 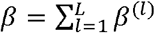 for a pre-specified number of causal variants *L*. As in SuSiE, we perform posterior inference for *β* using a variational approximation for the joint posterior distribution of *β*^(1)^, …, *β*^(*L*)^, again marginalizing over both *α* and *ϵ*. The approximation is computed by iterative stepwise optimization of an evidence lower bound (ELBO), where updated estimates of the variance components (*σ*^2^,*τ*^2^) are computed within each iteration using a method-of-moments approach.

The resulting models are similar to linear mixed models commonly used in contexts of association testing and phenotype prediction^22,24–26^. See **Discussion** for an explanation of why we do not apply existing methods for fitting linear mixed models.

Both methods take as input either the GWAS data (*y,X*) or sufficient summary statistics given by the un-standardized per-SNP z-scores 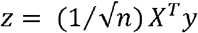, the in-sample LD correlation matrix LD = (1⁄*n*) *X*^*T*^ *X*, and the mean-squared phenotype ⟨*y*^2^⟩ = (1⁄*n*)*y*^*T*^ *y* . Both methods output estimates of (*σ*^2^,*τ*^2^) for each locus fine-mapped, together with a posterior-inclusion-probability (PIP) and posterior-mean effect size estimate for each SNP. Computational cost is reduced by expressing all operations in terms of the eigenvalues and eigenvectors of LD, which may be pre-computed separately for each fine-mapped locus (**Fig. 4**). Details of the methods and computations are provided in **Supplementary Note**. We have released open-source software implementing these methods (see **Code availability**).

**Fig. 4.**
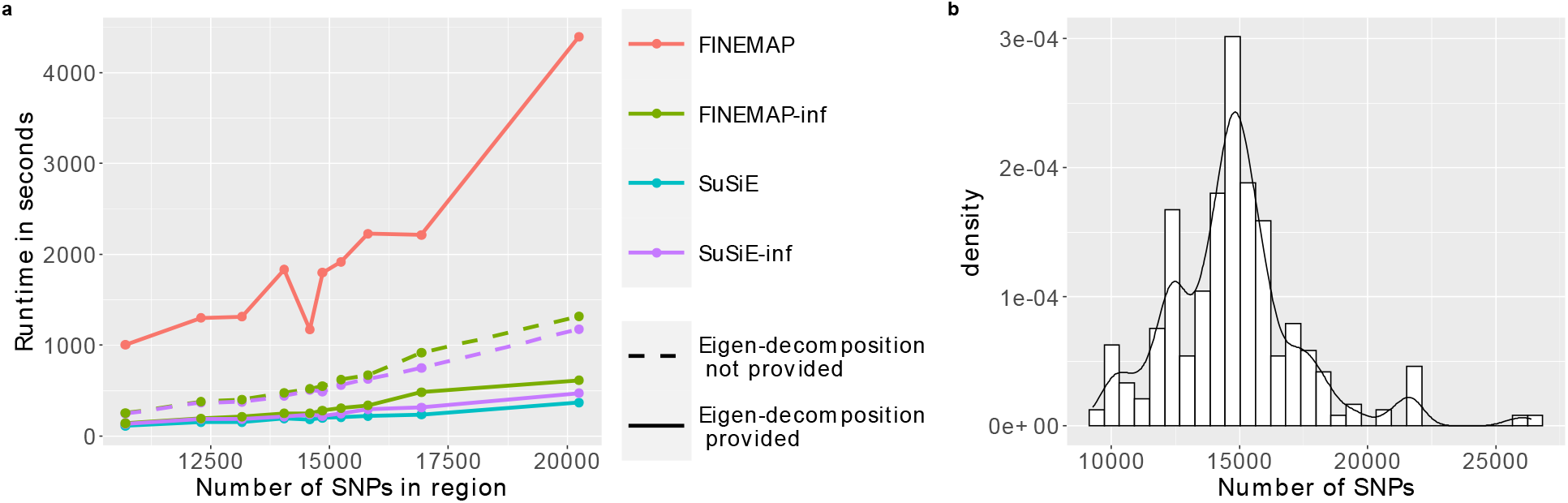
Runtime comparison. **a**. Average runtime in ten quantiles based on number of SNPs in fine-mapped region for SuSiE, SuSiE-inf, FINEMAP and FINEMAP-inf, as well as SuSiE-inf and FINEMAP-inf without provided eigen-decomposition of the LD matrix. **b**. Distribution of locus sizes in terms of number of SNPs, aggregated across 10 UKBB phenotypes and across two sample sizes: N=100K and N=366K. Numerical results available in **Supplementary Table 9**.

### SuSiE-inf and FINEMAP-inf show improved performance

In our simulations, we find that SuSiE-Inf and FINEMAP-Inf have improved calibration over SuSiE and FINEMAP, respectively, except for simulations using ordinary least squares (OLS) that introduced uncorrected population stratification, which are less relevant to our findings in real data using BOLT-LMM (**Fig. 2a, 3a, Online Methods**). Recall of SuSiE-Inf and FINEMAP-Inf was very similar to, but slightly lower than, that of SuSiE and FINEMAP, respectively (**Fig. 2b, 3b**). With improved performance in simulations having non-sparse genetic effects, and similar performance in simulations with stratification using BOLT-LMM (**Fig. 3a**), we turned to real data benchmarking to assess whether the new methods improve performance in practice.

Real data benchmarking shows improvements by several metrics. RFR was substantially decreased for SuSiE-inf (**Fig. 5a**). SuSiE-inf-specific high-PIP variants (variants that are assigned a high PIP by SuSiE-inf but not by SuSiE) are 58% more enriched in functionally important categories than SuSiE-specific high-PIP variants (P=6e-4); the analogous difference in functional enrichment for FINEMAP vs. FINEMAP-inf was non-significant (38% more for FINEMAP-inf specific variants, P=0.07, **Fig. 5b-c**). In additional to high-PIP variants identified with PIP>0.9, we also observed better functional enrichment for top N (N=500, 1000, 1500, and 3000) variants (**Extended Data Fig. 3a**), demonstrating better prioritization of variants by our methods. Similar improvements were observed when using OLS for GWAS instead of BOLT-LMM (**Extended Data Fig. 3b-c**), upon correcting for stratification using PCA. Compared to SuSiE and FINEMAP, we obtained fewer high-PIP variants (16% reduction aggregated between SuSiE and FINEMAP); however, the reduction is smaller for high-confidence variants, characterized either by replicated variants (11% reduction), or variants achieving PIP>0.9 for both SuSiE-Inf and FINEMAP-Inf/both SuSiE and FINEMAP (11% reduction) (**Extended Data Fig. 3d**). We observed a more substantial reduction of 42% in the number of credible sets when using SuSiE-inf; however, the reduction for smaller credible sets (number of variants < 10) was somewhat smaller (36% reduction). Credible sets generated by SuSiE-inf are smaller on average than those generated by SuSiE (**Extended Data Fig. 3e**). Together, these results demonstrate both that SuSiE-inf and FINEMAP-inf allow for more confident identification of likely causal variants than the current state of the art, and that there is room for further methodological improvement.

**Fig. 5.**
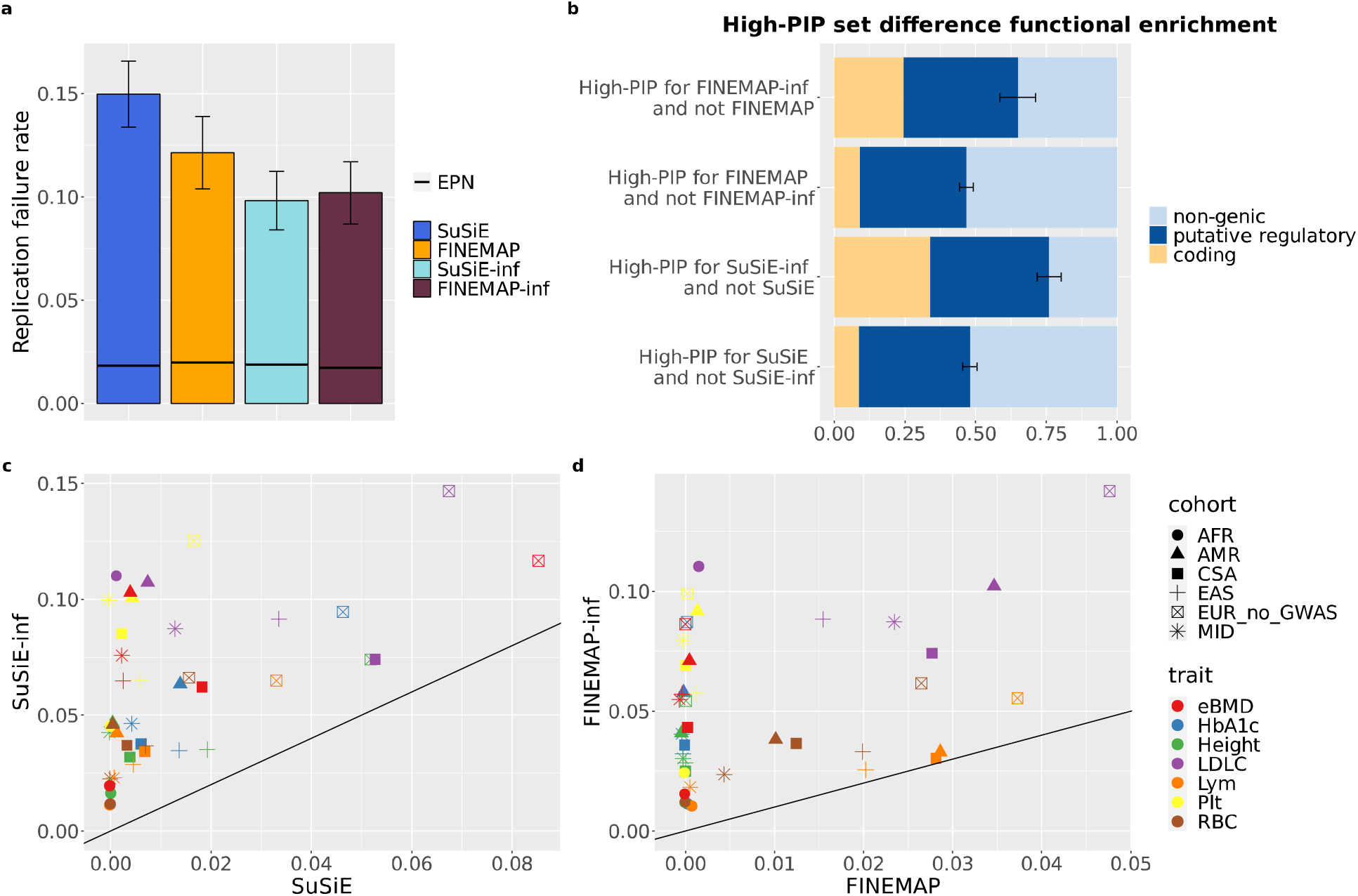
Real data performance improvements. **a**. Replication Failure Rates for SuSiE, FINEMAP, SuSiE-inf and FINEMAP-inf aggregated across 10 UKBB traits (**Supplementary Table 4**). **b**. Functional enrichment of the set difference between SuSiE and SuSiE-inf, and FINEMAP and FINEMAP-inf. Numerical results are available in **Supplementary Table 10. c-d**. Delta R^2^ comparison of PRS predictions using SuSiE vs. SuSiE-inf and FINEMAP vs. FINEMAP-inf. For standard errors, see **Extended Data Fig. 10**. Numerical results are available in **Supplementary Table 11-12**.

In simulation, estimates of the infinitesimal variance *τ*^2^ were higher on average for simulation settings with higher true infinitesimal variance (**Extended Data Fig. 4a-b**). Estimates of *τ*^2^ were also higher in the presence of more residual stratification in the simulations, fixing all other simulation parameters (**Extended Data Fig. 4c**). In UKBB data, estimates of *τ*^2^ varied across traits, with height showing the highest estimates and LDL showing the lowest estimates (**Extended Data Fig. 5a-b**). We also found that estimates of *τ*^2^ increased, on average, as the number of credible sets in a locus increased (**Extended Data Fig. 5c**). Estimates of *τ*^2^ varied across loci for a given trait, due either to differences in genetic architecture, residual stratification, or estimation noise. We caution against the interpretation of *τ*^2^ as a direct reflection of trait heritability or genetic architecture, without further investigations into these factors that may contribute to the *τ*^2^ estimates.

To further validate our methods in real data, we performed cross-ancestry Polygenic Risk Score (PRS) prediction^27,28^, using posterior effect sizes estimated on 366K samples from the “white British” cohort in UK Biobank to predict phenotypes in six held-out cohorts of different ancestries^29^: AFR (N=6637), AMR (N=982), CSA (N=8876), EAS (N=2709), EUR no-GWAS (N=54,337) and MID (N=1599). Prediction accuracy is measured by “delta R^2^” which is the difference in R^2^ from a model that includes both the covariates and genotype effects relative to a model that includes the covariates alone. Using posterior mean effect size estimates for the sparse component *β* in SuSiE-inf/FINEMAP-inf yields, on average, a near 10-fold increase in delta R^2^ over SuSiE/FINEMAP across all held-out cohorts and traits (**Online Methods, Fig 5c-d, Supplementary Table 3**). Here we compute PRS using only the sparse component, to provide a validation metric for the fine-mapped SNPs. We leave an exploration of improving PRS accuracy by integrating estimated infinitesimal effect sizes to future work.

Our group has shown previously that combining SuSiE and FINEMAP can yield more reliable PIPs^30^. Here we recommend using the minimum PIP between SuSiE-inf and FINEMAP-inf (minPIP-inf) for each fine-mapped variant. Compared to minPIP (minimum PIP between SuSiE and FINEMAP), minPIP-inf retains more high confidence variants, showing better agreement between SuSiE-inf and FINEMAP-inf (**Extended Data Fig. 6**). We observed substantially improved RFR for minPIP-inf over minPIP (**Extended Data Fig. 7a**). Functional enrichment for the top N variants, simulation and PRS performance for minPIP-inf is comparable to either SuSiE-inf or FINEMAP-inf individually (**Extended Data Fig. 7b-d, Fig. 2a-b**). As examples of the improved effectiveness of the minPIP-inf method over minPIP, we examined two loci. At the PCSK9 locus for LDLC, in addition to the well-known causal variant rs11591147, SuSiE-inf and FINEMAP-inf consistently identified two intronic variants: rs499883 and rs7552841 with high confidence, replicating a previous finding using functionally informed priors^31^, whereas SuSiE did not identify variant rs499883. At the AK3 locus for Plt, a known causal missense variant, rs7412, is in high LD with variant rs1065853. Only FINEMAP-inf captured rs7412 while SuSiE, SuSiE-inf and FINEMAP-inf captured another known causal variant, rs429358. MinPIP between SuSiE and FINEMAP missed both, whereas minPIP-inf captured one (**Extended Data Fig. 8-9**).

## Discussion

We propose fine-mapping methods that control for infinitesimal causal effects while fine-mapping sparse causal effects. Using our new methods, we observed significant improvements in simulations with non-sparse genetic architecture. Our results when simulating uncorrected stratification were ambiguous: when using BOLT-LMM, stratification did not lead to miscalibration and the new methods performed similarly to the previous methods; however, when using OLS, stratification led to substantial miscalibration that was similar between FINEMAP and FINEMAP-inf and worse for SuSiE-inf than SuSiE. In contrast, real data benchmarking demonstrated an unambiguous improvement in performance, e.g., decreased RFR, improved functional enrichment of top variants, and large gains in polygenic risk prediction.

Put together, the accuracy of identifying sparse causal variants is greatly improved when incorporating the infinitesimal model, although our results show that there is also room for further methodological improvement.

The models we propose here are similar to models that have been used previously to model genome-wide genetic architecture for risk prediction, heritability estimation, and association mapping^22,24,26,32^. Fine-mapping differs from these other applications in that (a) fine-mapping requires inclusion of a denser set of variants with higher LD in each locus, so that the causal variants are likely to be included; (b) fine-mapping requires accurate inference of posterior inclusion probabilities; (c) fine-mapping is often performed at very large sample sizes; and (d) fine-mapping does not require joint modeling of genome-wide data, which would be computational challenging given the density of variants and typical sample sizes. To emphasize the distinction between fine-mapping and risk prediction, if two variants are in perfect LD with large marginal effect sizes, a risk prediction method would perform equally well upon attributing this effect to either variant, whereas the desired outcome in fine-mapping is a more precise quantification of uncertainty for which variant(s) harbors the true effect. Because of these factors, we do not apply existing methods for fitting linear mixed models in other contexts. We instead extend algorithmic ideas in the fine-mapping literature to better estimate and quantify uncertainty for a sparse genetic component in the presence of strong LD, while estimating an infinitesimal variance component separately for each genome-wide significant locus. Our model incorporates infinitesimal effects for variants in LD with those of the sparse component, which we believe is important for obtaining improved calibration and fine-mapping accuracy. With careful translation, we anticipate that methodological innovations in risk prediction may continue to lead to advances in fine-mapping and vice versa.

We view our methods as complementary to a body of recent statistical developments that seek to accurately quantify and control false discoveries under weaker modeling assumptions, using constructions of knock-off variables and related conditional re-randomization ideas^33,34^. Such methods have been applied to GWAS and genetic fine-mapping applications in^35–37^. Our perspective differs in the following ways: We choose not to assume a sparse causal model or test null hypotheses of exact conditional independence, and instead aim to accurately identify large effects that drive observed GWAS associations in a model where every variant may be causal. To yield adequate statistical power for detecting causal variants at fine-mapping resolutions, we rely on a strong assumption about genetic architecture, as reflected by a Bayesian prior probability for each candidate model, rather than testing a null hypothesis for each variant that allows for an arbitrary genetic architecture excluding that variant. Thus, our methods remain largely model-dependent, and we view the potential integration of these ideas into a more model-robust framework as an important direction for future research.

While our work improves fine-mapping accuracy, further advances are needed. First, exploring the effects of stratification and different association mapping methods on fine-mapping should be a priority. Second, our methods improve on RFR over previous methods, but RFR is still elevated compared to ideal simulations, suggesting room for improved modeling. In addition to better modeling, independent replication in another biobank^30^ and incorporation of functional evidence such as annotations and eQTLs^20^ can help boost accuracy of discovery. Further methodological advancements, which may come from more flexible models of genetic architecture or further study of uncorrected confounding, may also contribute to further improvements in cross-population polygenic risk prediction.

## Online Methods

### Selection of UKBB phenotypes and down-sampling analysis

To select the 10 phenotypes for which to perform down-sampling analyses, we used results from ^30^ and computed the combined number of high-PIP (PIP > 0.9) variants fine-mapped at N = 366K samples using both SuSiE and FINEMAP. From the top 15 phenotypes (out of 94) with the highest number of high-PIP variants (**Supplementary Table 13**) we selected: height, estimated heel bone mineral density (eBMD), platelet count (Plt), hemoglobin A1c (HbA1c), red blood cell count (RBC), alkaline phosphatase (ALP), insulin-like growth factor 1 (IGF1), low density lipoprotein cholesterol (LDLC), lymphocyte count (Lym), and estimated glomerular filtration rate based on serum creatinine (eGFR) to perform down-sampling analyses.

We down-sampled from N=366K to a random subset of N=100K twice (to increase the number of discoveries and therefore statistical power for RFR analyses) and performed GWAS and fine-mapping on both set of the N=100K individuals using the same pipeline used at N=366K (see below for pipeline description).

### Fine-mapping pipeline

GWAS and fine-mapping in this paper were performed following the pipeline described in our group’s previous work^30^. Briefly, GWAS summary statistics were computed using BOLT-LMM with covariates including sex, age, age2, age and sex interaction term, age2 and sex interaction term, and the top 20 genotype Principal Components (PCs). Fine-mapping regions were defined using a 3Mb window around each lead GWAS variant, with merging of overlapping regions. Fine-mapping was performed with in-sample LD computed using LDstore v2.0^38^.

Excessively large regions (consequence of merging) that could not be fine-mapped due to computational limitations were tiled with overlapping 3Mb loci, with 1Mb spacing between the start points of consecutive loci. For these tiled regions, we computed a PIP for each SNP based on the 3Mb locus whose center was closest to the SNP. This tiling approach was previously described and applied in^31^.

Although BOLT-LMM is the GWAS method-of-choice in our group’s previous work^30^, we also used ordinary least squares (OLS) regression for some of our simulations and real data applications.

### Ideal simulations

To establish reference Replication Failure Rate (RFR) and calibration for all tested methods, we performed ideal simulations without model misspecification using UK Biobank genotypes. For simulating RFR, we performed two sets of simulations each at sample size N = 366K and subsample size N = 100K. We used UK Biobank imputed dosages as true genotypes, and only selected “white British” individuals defined previously in the Neale lab GWAS^21^ https://github.com/Nealelab/UK_Biobank_GWAS). We drew 1000 causal variants per simulation uniformly randomly from a total of 6.6 million common (MAF ≥ 1%) imputed variants genome-wide. We standardized genotypes to mean 0 and variance 1 and drew per-standardized-genotype causal effect sizes from the same normal distribution N (0, 0.5/1000) for all selected causal variants. We then added errors randomly drawn from a normal distribution N (0, 0.5) to simulate phenotypes. For comparison of calibration with our simulations under model misspecifications, three additional sets of ideal simulations at a matching sample size N = 150K were performed. Phenotypes were generated similarly, with 700 uniformly sampled true causal variants having effect sizes drawn from N (0, 0.5/700).

### Functional enrichment

We analyzed functional annotations to gain insights into the potential causal status of non-replicating variants (defined in the main text and in the next paragraph). We define three main disjoint functional categories: coding, putative regulatory, and non-genic. These categories are derived from the seven main functional categories defined in^30^. The “coding” category is the union of pLoF and missense categories; the “putative regulatory” category is the union of synonymous, 5’ UTR, 3’ UTR, promoter, and cis-regulatory element (CRE) categories. We compare the proportion of non-genic variants in the following groups of variants:

1. Non-replicating, the set of variants with PIP ≥ 0.9 at N = 100K and PIP ≤ 0.1 at N = 366K.
2. Replicated, the set of variants with PIP ≥ 0.9 at N = 100K and PIP ≥ 0.9 at N = 366K.
3. Matched on PIP at 100K, a set of replicated variants chosen to match the non-replicating variants on PIP at N = 100K. For each non-replicating variant with PIP<1, we find a replicated variant whose PIP is the closest as its match, and the matched variant is removed for future matches. If the non-replicating variant has PIP = 1, we match a random (if there are multiple) replicated variant with PIP = 1. If there are more non-replicating variants with PIP = 1 than there are replicated variants with PIP = 1, we do not remove the matched replicated variant from future matches, resulting in repeated matches.
4. Matched on PIP at 366K, a set of low-PIP variants (PIP ≤ 0.1 at N=366K) chosen to match the non-replicating variants on PIP at N = 366K. Matching is performed the same way as described above, except that there are no repeated matches.
5. Background, defined as the union of all variants included in fine-mapping from all 10 phenotypes.

P-values are reported when assessing the significance of the difference between proportions of non-genic variants in different groups of variants. Fisher’s exact test was performed using the R function fisher.test, and one-sided p-values were reported from the output of this function.

## Large-scale simulations with misspecification

We selected 149,630 UK Biobank individuals from a set of 366,194 unrelated “white British” individuals defined previously in the Neale lab GWAS^21^ for our large-scale simulations. We performed simulations under models that are misspecified in the following ways: (1) genotype imputation noise, (2) non-uniform probabilities for the identities of causal variants, (3) non-sparsity of true causal effects, (4) uncorrected population stratification, and (5) missing causal variants. We performed 9 sets of simulations. All simulations included the same amount of (1) imputation noise, (2) non-uniform prior causal probabilities, and (5) missing causal variants. The first simulation, “baseline misspecification” in **Table 1**, included a small amount of (4) uncorrected stratification. Another four simulations varied, in addition, (3) the level of non-sparsity of causal effects. Finally, four additional simulations varied (4) the amount of simulated stratification and the methods for correcting this stratification (see **Population stratification** below).

### Genotypes

To simulate genotypes for 149,630 individuals, we randomly drew true genotypes for all autosomes based on the imputed genotype probabilities in the “bgen” files provided by UKBB. Briefly, probabilistic true genotypes (*pGT*s) for a given variant *i*, denoted *pGT*_*i*_, were computed via *pGT*_*i*_ = ⌈*u*_*i*_ − *GP*(*X*_*i*_ = 0)⌉ + ⌈*u*_*i*_ − *GP*(*X*_*i*_ = 0) − *GP*(*X*_*i*_ = 1)⌉, where *GP*(*X*_*i*_ = *k*), *k* ∈ {0,1,2} represents the genotype probability of having *k* copies of alternative alleles and *u*_*i*_ ∼ *Uniform*(0,1) represents a uniform random variable. Phenotypes were generated using the *pGT*s. In downstream GWAS and fine-mapping, we use imputed genotype dosages provided by UKBB, thus simulating imputation noise. We only included variants with minor allele count > 10, INFO score > 0.2, and Hardy-Weinberg equilibrium p-value > 1e-10 in our simulations.

### Causal variants

To incorporate a more realistic non-uniform distribution over causal variants, we simulated sparse causal effects from the SuSiE posterior distribution for UKBB Height, as computed in the larger 366K sample in^30^. Specifically, in each locus, for each credible set *cs*_*i*_,*i* ∈ *I*, where I indexes all credible sets outputted by fine-mapping Height with SuSiE, we chose a causal variant according to normalized posterior inclusion probabilities within the corresponding SuSiE single effect (denoted *α*_*ik*_ for *k* ∈ *cs*_*i*_). We then drew the chosen variant’s raw effect size (to be scaled later) from a normal distribution with mean and standard deviation given by the SuSiE posterior mean and standard deviation conditional on inclusion in the model. In total, 1434 sparse causal variants were chosen.

For the simulations that investigated non-sparsity of causal effects, we drew additional causal variants uniformly at random such that x% (*x* ∈ {0.5,1,5}) of all simulated variants have a non-zero effect. For each selected variant, we sampled its raw effect size (to be scaled later) from 𝒩(0, *v*) where *v* = [2*p*(1−*p*)]^*α*^, *p* represents the MAF, and *α* = −0.38. The value alpha is estimated in ^39^. For all simulation settings, simulated non-sparse effects had an overall effect size standard deviation approximately on the order of 1e-4 units per normalized genotype.

We simulated 3 settings of non-sparsity coverage: 0.5%, 1% and 5%, where coverage is the percentage of variants with non-zero effects on the phenotype. For the simulations with 1% coverage, we varied the heritability explained by the non-sparse causal variants, it was set to be 58%, 75%, 83% and 100%, corresponding to heritability ratios between sparse and non-sparse causal effects of 1-to-1.4, 1-to-3, 1-to-5, and 0-to-1. To achieve these heritability proportions, we scaled all the simulated sparse and non-sparse causal effect sizes by corresponding constants. We observed that, for all simulation settings, simulated large effects had an overall effect size standard deviation approximately on the order of 1e-2 units per normalized genotype. For the simulations with 0.5% coverage and one set of simulations with 5% coverage we fixed the heritability ratio to 1-to-3. We performed an additional set of simulations with non-sparsity coverage of 5% and heritability ratio between sparse and non-sparse causal effects of 1-to-15. The purpose of this setting is to match the simulated per-SNP heritability with the 1% coverage 1-to-3 ratio simulations. See **Table 1** for a summary of the different settings. We set the total SNP heritability to be 0.5. Note that the 0.5 heritability accounts for all simulated causal SNPs and not just the common SNPs. We have computed s-LDSC estimates of common SNP heritability for all the simulations and all 10 UK Biobank phenotypes, and the results are available in **Supplementary Table 1-2**.

Interestingly, changing the coverage of non-sparsity from 0.5% to 1% then to 5% while fixing the proportion of heritability explained by non-sparse effects showed a non-monotonic behavior in the level of mis-calibration. This is likely due to multiple factors influencing calibration: per-SNP heritability of non-sparse effects and LD between non-sparse and sparse causal variants. We observed increased mis-calibration when per-SNP heritability is fixed and coverage changes from 1% to 5%. Similarly, when coverage is fixed at 1% and per-SNP heritability increases by 50% calibration also worsens (**Fig. 2, Extended data Fig. 2**).

### Population stratification

To simulate population stratification, we first regressed UKBB Height on the top 20 principal components (PCs) of the genotyped variants for N = 360,415 individuals. We then added the sum of the principal component scores multiplied by their respective regression coefficients to the simulated phenotype, scaling this sum by a factor to vary the amount of simulated stratification. We assessed the amount of stratification by running s-LDSC^40^ on the resulting GWAS summary statistics (without using any in-sample PCs as covariates) and examining the fitted intercept (**Supplementary Table 2**). As expected, we see higher s-LDSC intercept as we increase the PC scaling factor.

For the stratification simulations referenced in the main text and **Table 1**, we scaled PC effects by a factor of 5 (resp. 8) for moderate (resp. severe) stratification with BOLT, yielding a phenotype with 16.4% (resp. 42.9%) of its variance explained by stratification. For stratification with OLS, we scaled PC effects by 1 and 2 for moderate and severe stratification, yielding phenotypes with 0.6%, 2.6% of their variance due to stratification, respectively. s-LDSC intercepts of the stratification simulations are available in **Supplementary Table 2**.

### Phenotype

Phenotypes were generated as *y* = *Xβ* + *Cζ* + *ϵ*, where *X* is the above true genotype (*pGT*) matrix, *β* is a vector of the (sparse and non-sparse) causal effects, *C* is a matrix with top 20 principal components with corresponding effects *ζ*, and *ϵ* ∼ 𝒩(0, *σ*^2^*I*_*n*_) where *σ*^2^ was chosen to yield total phenotypic variance equal to 1.

### Missing causal variants

After generating phenotypes and before performing GWAS and fine-mapping, we applied variant-level quality-control criteria as previously defined in the Neale lab GWAS^21^, which retained 13,364,303 variants after filtering for: INFO > 0.8, MAF > 0.001, and Hardy-Weinberg equilibrium P value > 1e-10, with exception for the VEP-annotated coding variants where we allowed MAF > 1e-6. Notably, this QC step resulted in the exclusion of approximately 71% of the simulated “non-sparse” causal variants.

### GWAS and fine-mapping

We performed GWAS on N = 149,630 individuals using BOLT-LMM^22^ v2.3.2, with corresponding imputed variant dosages from UKBB. We used the top 19 principal components computed in-sample as covariates in the GWAS, except in the population stratification simulations, which included no covariates. For some of the population stratification simulations, we performed GWAS with ordinary least squares regression, rather than BOLT-LMM. We performed OLS using the linear regression rows method in Hail^41^ v0.2.93. For fine-mapping we used the pipeline previously described in section **Fine-mapping pipeline**.

### Interpreting population stratification simulation results

When scaling PC effects by a factor of 5 and computing GWAS summary statistics using BOLT-LMM, we observed an s-LDSC intercept of 1.07, which is comparable to s-LDSC intercepts estimated in real complex traits (**Supplementary Table 1**), and we did not observe significant miscalibration in the downstream fine-mapping results. When we simulated a higher level of uncorrected stratification, scaling PC effects by a factor of 8 (s-LDSC intercept of 1.16, see “Severe stratification with BOLT” in **Table 1**), PIPs obtained in downstream fine-mapping remained well-calibrated (**Fig. 3**).

We hypothesize that the use of BOLT-LMM in our standard fine-mapping pipeline helped to correct for the simulated stratification effects, even though the in-sample PCs were not explicitly provided as covariates. This also likely explains the prima facie surprising recall results in **Fig. 3** where the severe stratification simulations with BOLT have higher recall than the moderate stratification simulations with BOLT. In the severe simulations, stratification accounts for 42.9% of the phenotypic variance, whereas in the moderate simulations, stratification accounts for only 16.4% of phenotypic variance. Because BOLT-LMM likely corrects for much of this simulated stratification, it effectively reduces the residual noise in the associations by much more for the severe simulations than for the moderate ones, allowing fine-mapping to nominate more causal variants.

To investigate stratification effects without using an LMM procedure, we performed 2 additional sets of simulations where GWAS summary statistics were instead computed using ordinary least squares (OLS). In these simulations, scaling PC effects by factors of 1 and 2 yielded average s-LDSC intercepts of 1.055 and 1.295, respectively (**Supplementary Table 2**), and induced significant miscalibration across all methods. This miscalibration was more severe for SuSiE-inf and FINEMAP-inf than for SuSiE and FINEMAP (**Fig. 3**).

It is unclear to us which of these simulation settings may be closer to reflecting the possible effects of uncorrected stratification in real fine-mapping applications, given that common methods of computing GWAS summary statistics do use LMM procedures and, in addition, explicitly control for in-sample PCs as covariates. Our real-data results, including functional enrichment and PRS analyses, in UKBB show evidence that SuSiE-inf and FINEMAP-inf outperform existing methods in real data. We leave to future work a fuller investigation of the possible effects of uncorrected stratification on downstream fine-mapping, and a potential extension of these methods to address uncorrected stratification.

### Regression of phenotype components on high-PIP variants

To identify which of several simulated model misspecifications were responsible for observed miscalibration, we decomposed the simulated genetic component *Xβ* of the phenotype into the sum of four sub-components representing sparse causal effects, non-sparse causal effects, non-sparse causal effects due to QC, and the effects of stratification. That is,

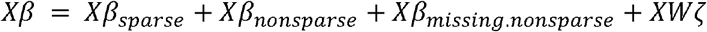

where *W* is an *n* × 20 matrix of UKBB PC loadings computed at a sample size of 360,415 and ζ is a 20 × 1 vector of regression coefficients for the top 20 PCs on UKBB Height at 360,415. For each simulation, we regressed each of the four genetic effect sub-components on each of the PIP > 0.9 variants independently, with 19 in-sample (n=149,630) PCs as covariates in the regression (i.e., the same covariates we use in GWAS in our simulations). For example, for the sparse genetic effect component, we compute the regression coefficient b and its associated F-statistic for the following equation:

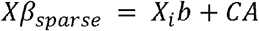

where variant *i* is the index of a PIP > 0.9 variant and *C* is a matrix of 19 in-sample PCs. We then compare the F-statistics of truly causal and non-causal variants.

## Polygenic Risk Score (PRS)

### Cohort assignment

We used six ancestry groups derived by the Pan UKBB project^29^: EUR = European ancestry (N=420,531, training=366,194, testing=54,337), CSA = Central/South Asian ancestry (N=8,876), AFR = African ancestry (N=6,636), EAS = East Asian ancestry (N=2,709), MID = Middle Eastern ancestry (N=1,599), and AMR = Admixed American ancestry (N=980). The 1000 Genomes Project and Human Genome Diversity Panel (HGDP) were used as reference panels to assign continental ancestry.

### Weights

We chose seven phenotypes: HbA1c, Height, LDLC, Lym, Plt, RBC and eBMD for PRS predictions. We fine-mapped these seven phenotypes on the training cohort: EUR (QC’ed from N=420531 to N=366,194 unrelated “white British” individuals). SuSiE, FINEMAP, SuSiE-inf and FINEMAP-inf posterior effect sizes were obtained. Due to differences in computational efficiency, not all variants that are eligible for fine-mapping were able to be fine-mapped by all methods. To ensure fair comparison between SuSiE and SuSiE-inf (resp. FINEMAP and FINEMAP-inf), we include only variants that were fine-mapped by both SuSiE and SuSiE-inf (resp. FINEMAP and FINEMAP-inf) in the PRS analyses (**Supplementary Table 3**). PLINK2.0^42^ was then used to compute polygenic risk scores for the six held-out cohorts using these posterior effect sizes. For SuSiE-inf and FINEMAP-inf we assigned weights to variants using the estimated posterior effect sizes from the sparse effects *β* and did not add the estimated posterior effect sizes of the infinitesimal effects *α*.

### Accuracy metric

We use delta R^2^ as our accuracy metric for PRS predictions, as in^43^. To obtain delta R^2^, we fit two models:

- Model 0: a linear model using only covariates as predictor, denoted model0.
- Model 1: a linear model using true phenotype as target and both the PRS generated from multiplying the fine-mapped posterior effect size estimates with the genotypes and the covariates (sex, age, age^2^, age and sex interaction term, age^2^ and sex interaction term) as predictors.

We applied the function *lm* in R to obtain *adj*.*r*.*squared*. The difference: *summary(model1)$adj*.*r*.*squared − summary(model0)$adj*.*r*.*squared* is delta R^2^.

## Data availability

The main fine-mapping results at N=100K sample size produced by this study are publicly available at https://doi.org/10.5281/zenodo.7055906. The fine-mapping results at N=366K previously produced by our group is available at https://www.finucanelab.org/data. The UKBB individual-level data is accessible on request through the UK Biobank Access Management System (https://www.ukbiobank.ac.uk/). The UKBB analysis in this study was conducted via application number 31063.

## Code availability

Software implementing SuSiE-inf and FINEMAP-inf are publicly available: https://github.com/FinucaneLab/fine-mapping-inf. The code to generate all figures in this manuscript is available at https://github.com/cuiran/improve-fine-mapping.

## Acknowledgements

This research has been conducted using the UK Biobank Resource under Application Number 31063. This work is supported by the Novo Nordisk Foundation (NNF21SA0072102). M.K. was supported by a Nakajima Foundation Fellowship and the Masason Foundation. We acknowledge all the participants of UK Biobank. We thank all the members of Finucane lab and Analytic and Translational Genetics Unit (ATGU) at Massachusetts General Hospital for their helpful feedback.

## Extended data figures

**Extended Data Fig. 1.**
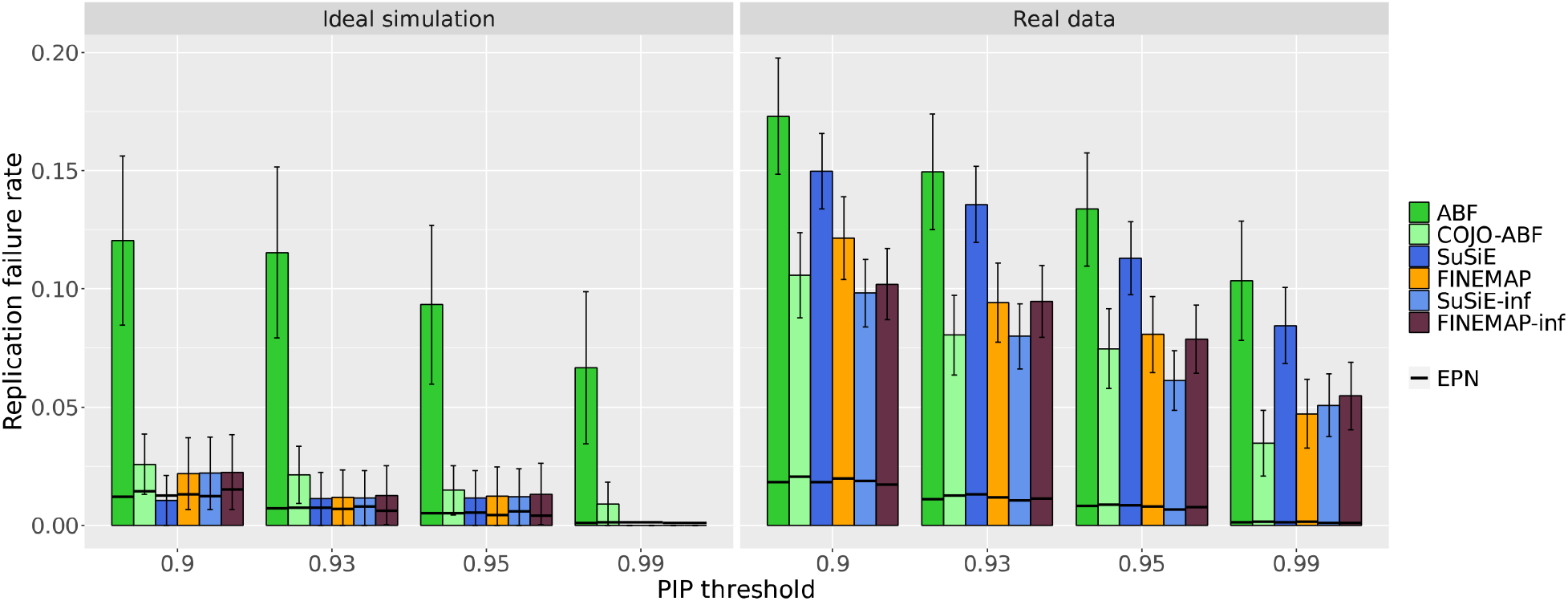
Replication failure rates at different PIP thresholds. RFR and EPN for six methods in ideal simulations and in real data aggregated across 10 UKBB phenotypes. High-PIP variants defined at four different PIP thresholds: 0.9, 0.93, 0.95, and 0.99. Numerical results available in **Supplementary Table 14**.

**Extended Data Fig. 2.**
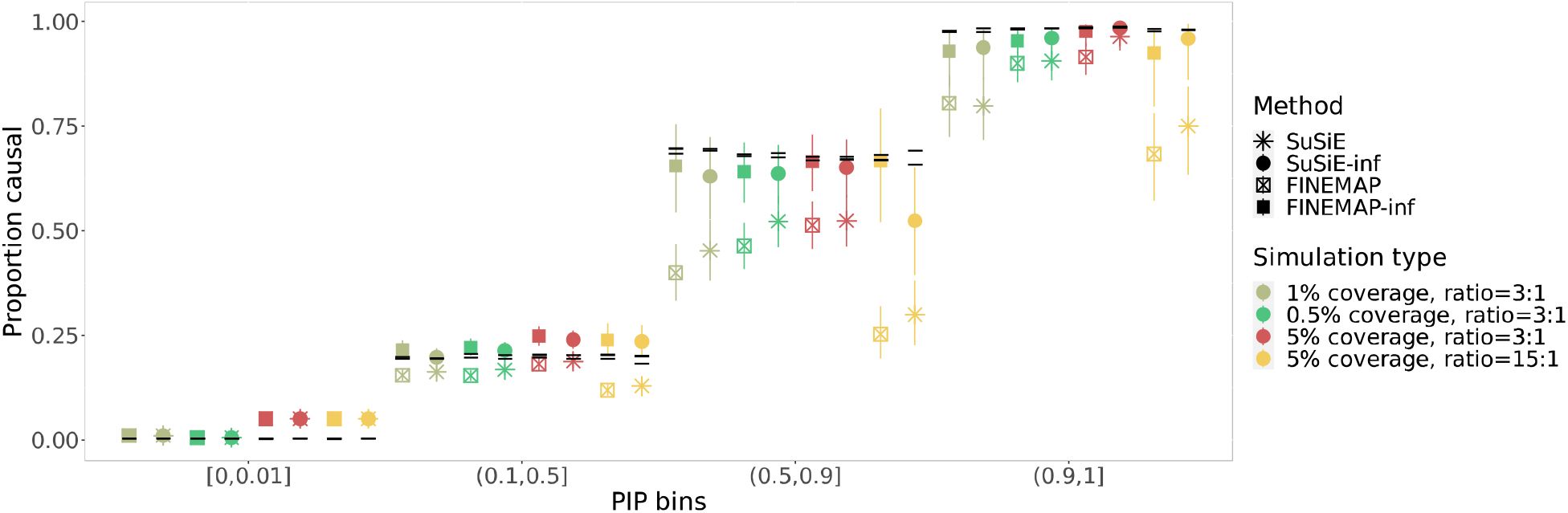
Calibration plot for varying levels of non-sparsity coverage. Calibration comparison for SuSiE, SuSiE-inf, FINEMAP and FINEMAP-inf in simulations with different non-sparsity coverage: 0.5%, 1% and 5%. Heritability ratio between small and large effects is fixed at 3:1 for three simulation scenarios with varying coverage setting, while in one simulation with coverage 5% we set the heritability ratio to be 15:1 to match the per-SNP heritability in the simulation where 1% SNPs are causal and heritability ratio is 3:1. Numerical results available in **Supplementary Table 15**.

**Extended Data Fig. 3.**
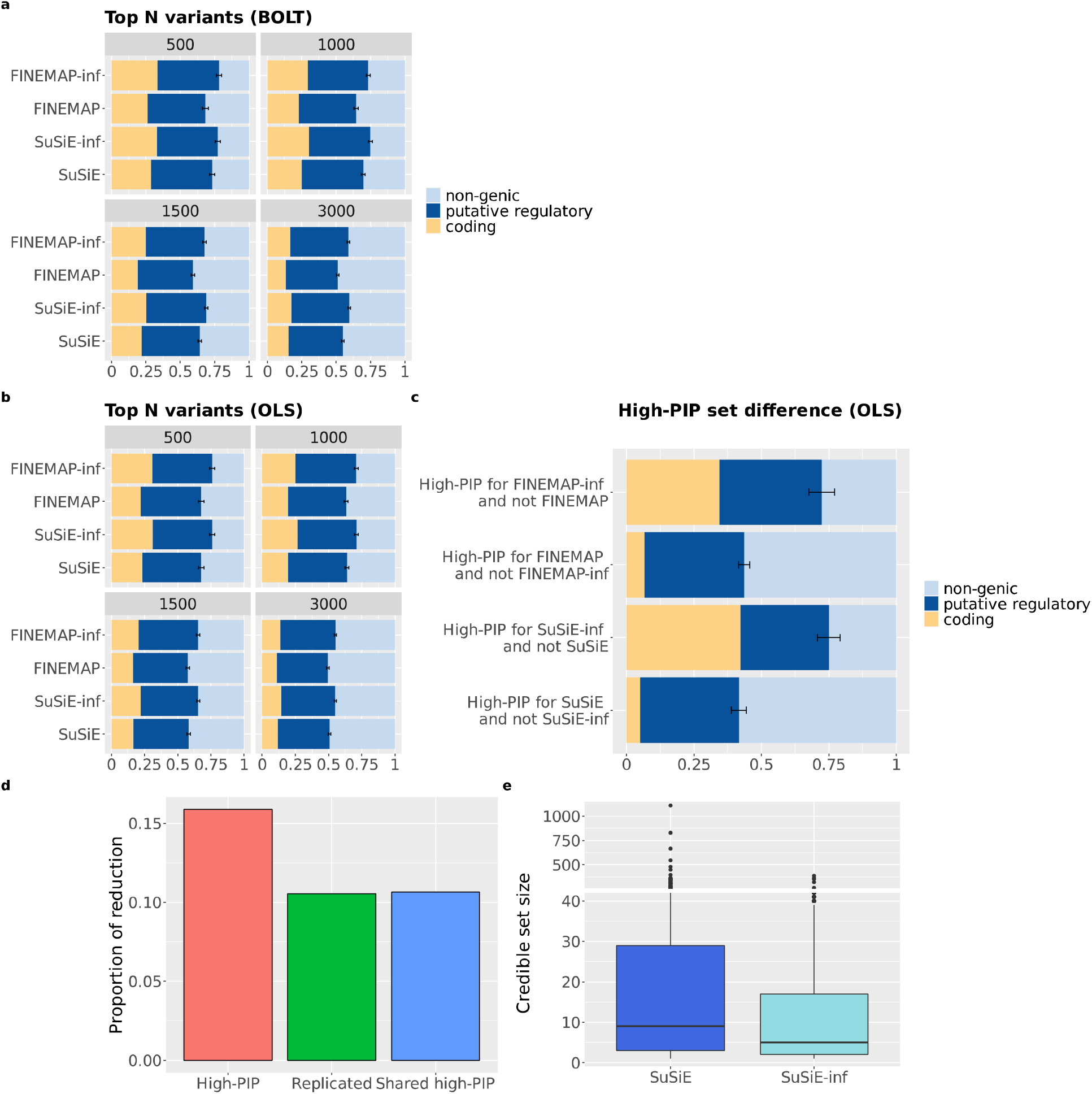
Additional evidence of performance improvements on real data. **a**. Functional enrichment of top 500, 1000, 1500, and 3000 highest PIP variants from different methods. **b**. Functional enrichment of top N (N=500, 1000, 1500, and 3000 resp.) highest PIP variants compared amongst four methods, with GWAS summary statistics computed using OLS instead of BOLT-LMM. **c**. Functional enrichment of the set difference between SuSiE and SuSiE-inf, FINEMAP and FINEMAP-inf with GWAS summary statistics computed using OLS instead of BOLT-LMM. **d**. The proportion of reduction for the number of high-PIP variants aggregated across SuSiE and FINEMAP at N=100K when using either SuSiE-inf or FINEMAP-inf than using either SuSiE or FINEMAP. **e**. Numerical results available in **Supplementary Table 16-20**.

**Extended Data Fig. 4.**
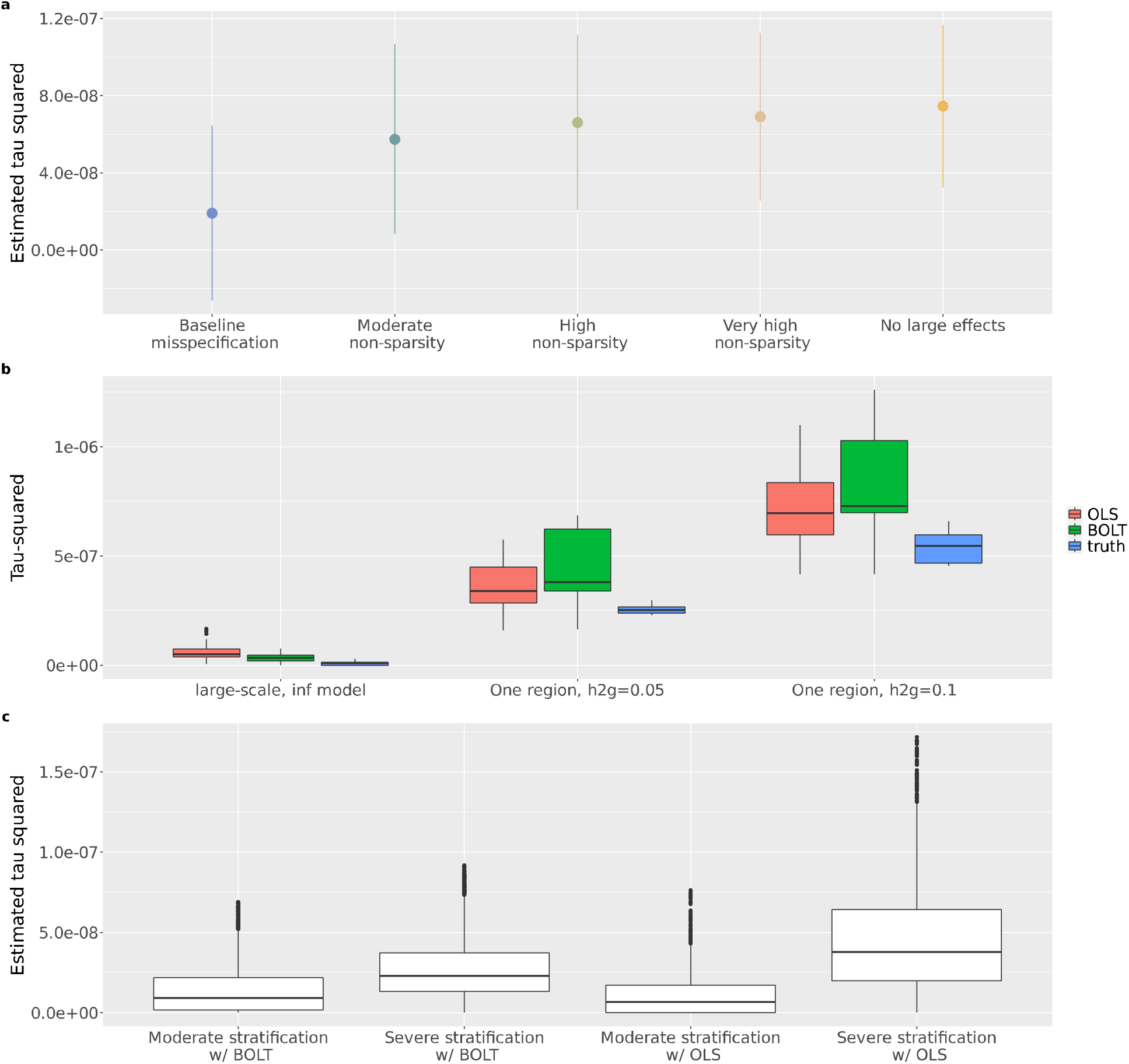
Estimated tau-squared in simulations. **a**. The estimated tau-squared in all regions are plotted for each non-sparse simulation setting. (**Table 1, Online Methods**). **b**. Estimated tau-squared and true tau-squared in three simulations are plotted. “Large-scale, inf model” represents the set of large-scale simulations (described in **Online Methods**) with 100% causal coverage setting and no missing causal variants are introduced. “One region, h2g=0.05” represents the set of simulations using imputed genotypes in one region on Chromosome 1, with 100% causal coverage, no missing causal variants and no exclusion of variants in the fine-mapping pipeline. The total SNP heritability is set to be 0.05. “One region, h2g=0.1” is similar except with total SNP heritability set to be 0.1. **c**. Boxplots of estimated tau-squared in stratification simulations. See **Table 1** for description of these simulations. Numerical results available in **Supplementary Table 21-23**.

**Extended Data Fig. 5.**
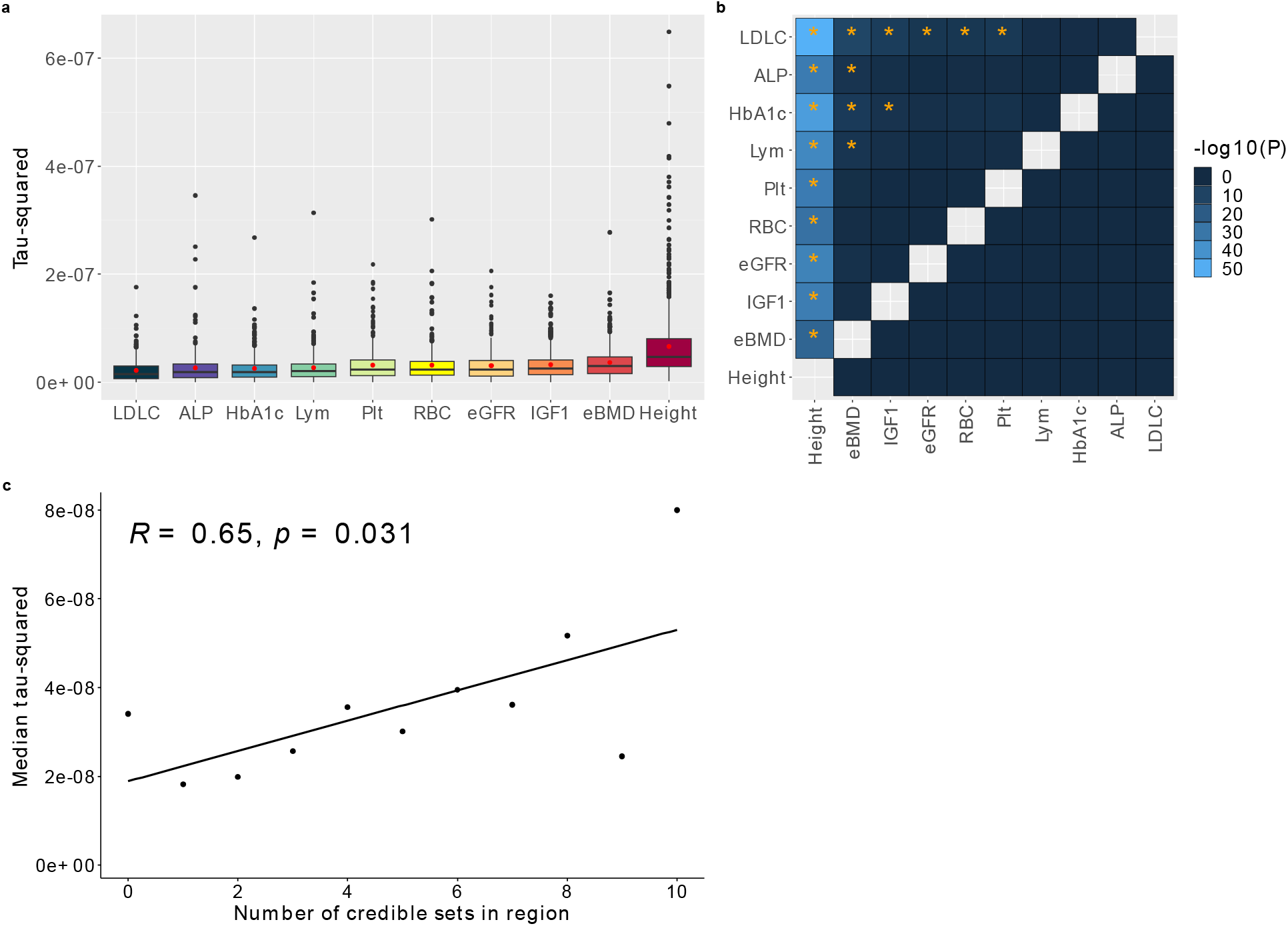
Estimated tau-squared (infinitesimal variance) in UKBB. **a**. Boxplot for estimated tau-squared in all fine-mapped regions for 10 tested phenotypes. The line across the boxplots denotes the median, the red dot denotes the mean. **b**. Two-sample T-test with alternative hypothesis: mean of estimated tau-squared for all fine-mapped regions for trait1 (x-axis) is greater than that of trait2 (y-axis). P-value cutoff is set to be 0.05/90 = 5.5e-4, correcting for the total number of trait pairs tested. Stars indicate P-values pass the significant threshold. **c**. Correlation between number of credible sets and the infinitesimal variance. Medians of tau-squared are computed for regions with the same number of credible sets. R is the Pearson correlation; p is the correlation p-value. The 95% confidence interval is shown on the plot as the gray shaded area. Numerical results available in **Supplementary Table 24-26**.

**Extended Data Fig. 6.**
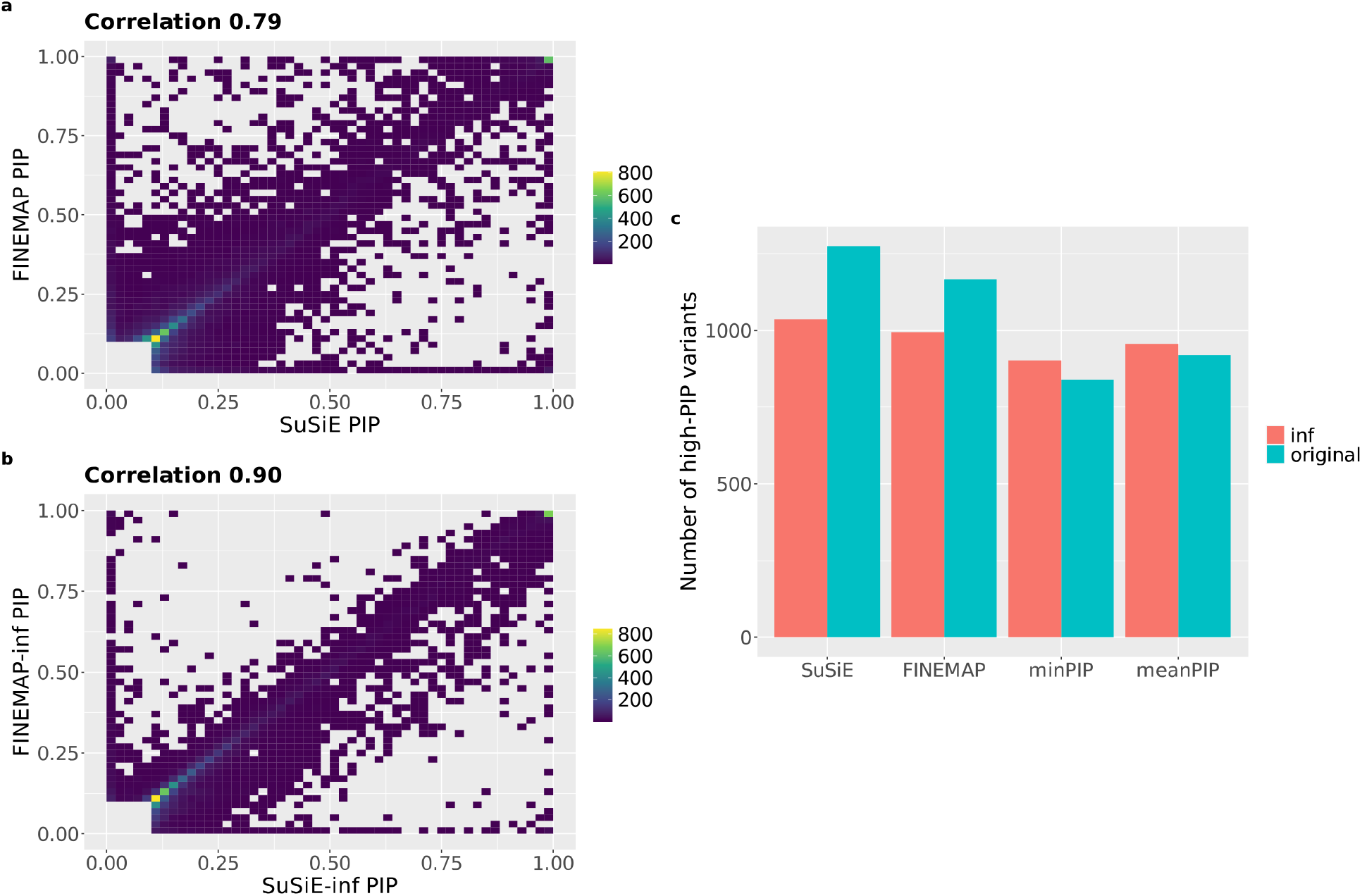
Agreement between SuSiE and FINEMAP vs. SuSiE-inf and FINEMAP-inf. **a-b**. Density plot of PIPs from 10 UKBB traits fine-mapped at N=366K. All variants with PIP>=0.1 for either method are shown on the density plots. **c**. Number of high-PIP (PIP>0.9) variants identified by SuSiE, SuSiE-inf, FINEMAP, FINEMAP-inf, minPIP, minPIP-inf, meanPIP and meanPIP-inf, where meanPIP(-inf) is defined as taking the average PIP between SuSiE and FINEMAP (resp. SuSiE-inf and FINEMAP-inf). Data aggregated across 10 UKBB traits fine-mapped at N=366K. Numerical data available in **Supplementary Table 27**.

**Extended Data Fig. 7.**
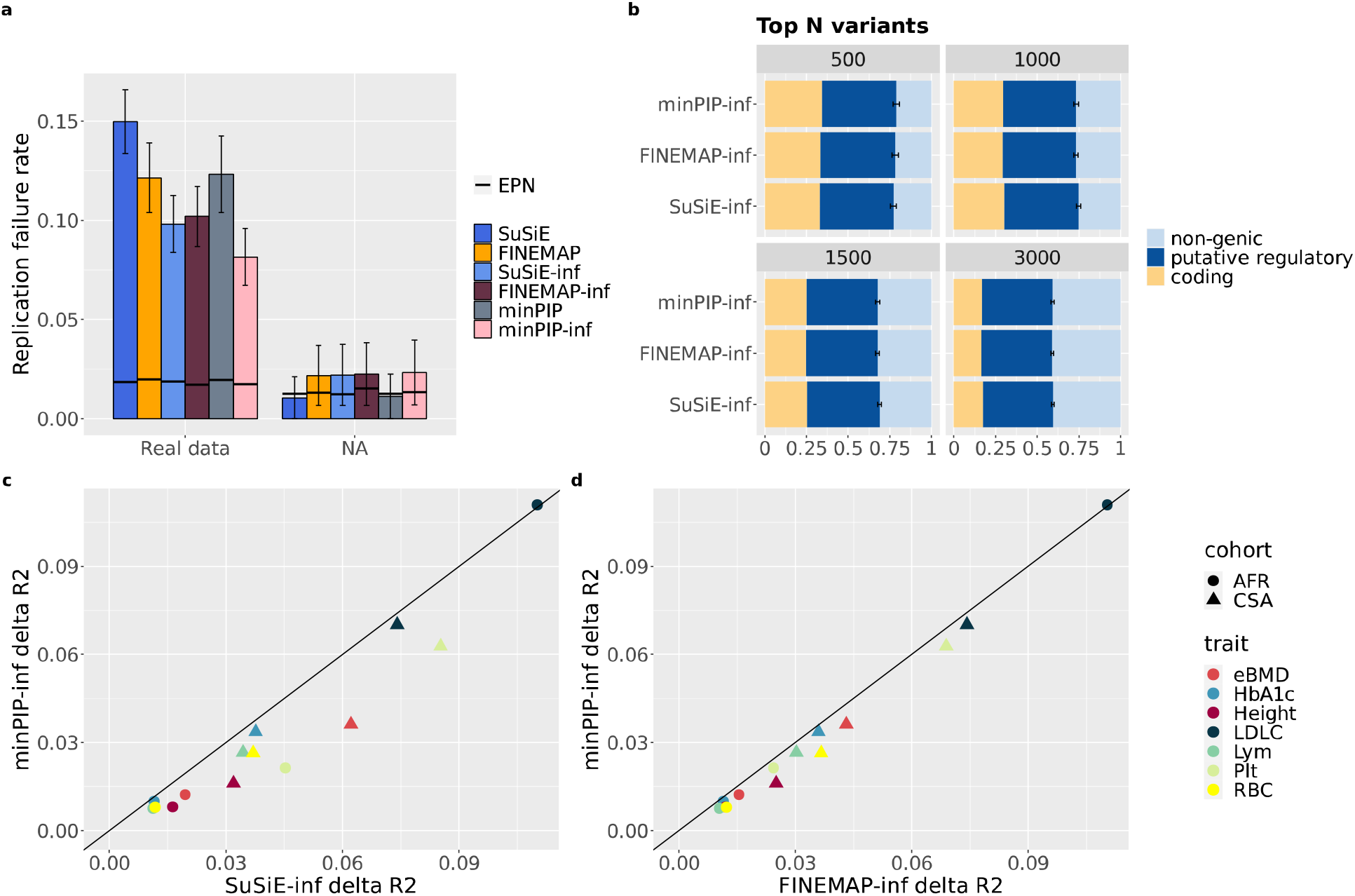
minPIP-inf performance. **a**. Replication failure rate of minPIP and minPIP-inf compared to other methods (**Supplementary Table 4**). minPIP = taking min(PIP) between SuSiE & FINEMAP; minPIP-inf = taking min(PIP) between SuSiE-inf and FINEMAP-inf. **b**. Functional enrichment of top 500, 1000, 1500, and 3000 highest PIP variants from different methods (**Supplementary Table 16**). **c-d**. Comparison of PRS accuracy (measured by Delta R^2^) of selected cohorts between minPIP-inf and SuSiE-inf (resp. FINEMAP-inf) (**Supplementary Table 11**).

**Extended Data Fig. 8.**
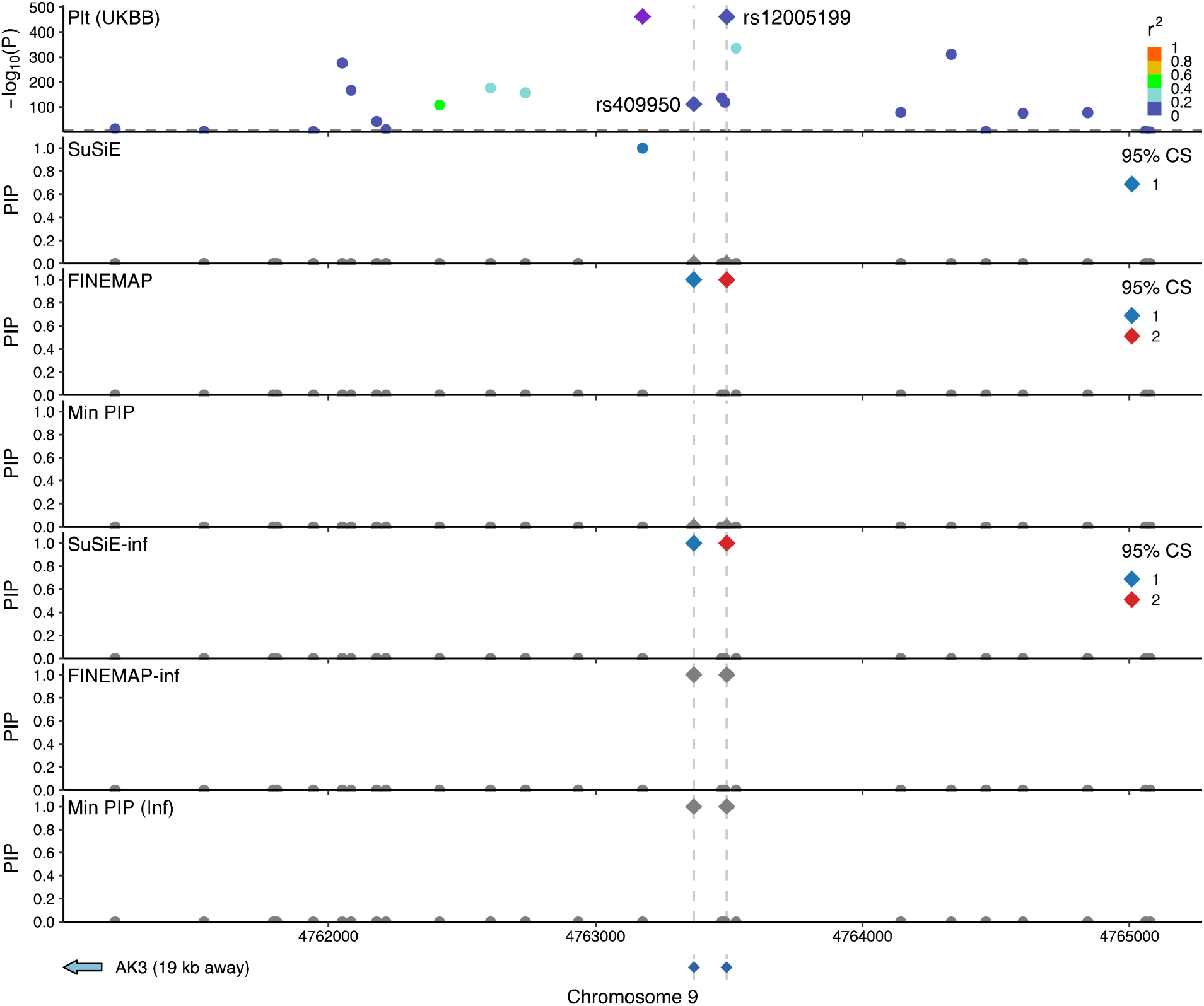
AK3 locus for Plt. 4kbp window near the AK3 gene is shown on the plot, GWAS -log10 P-values for trait Plt are plotted on the top panel, PIPs from 4 fine-mapping methods and 2 aggregating methods are plotted on the subsequent panels. Variants rs12005199 and rs409950 are highlighted using dashed lines.

**Extended Data Fig. 9.**
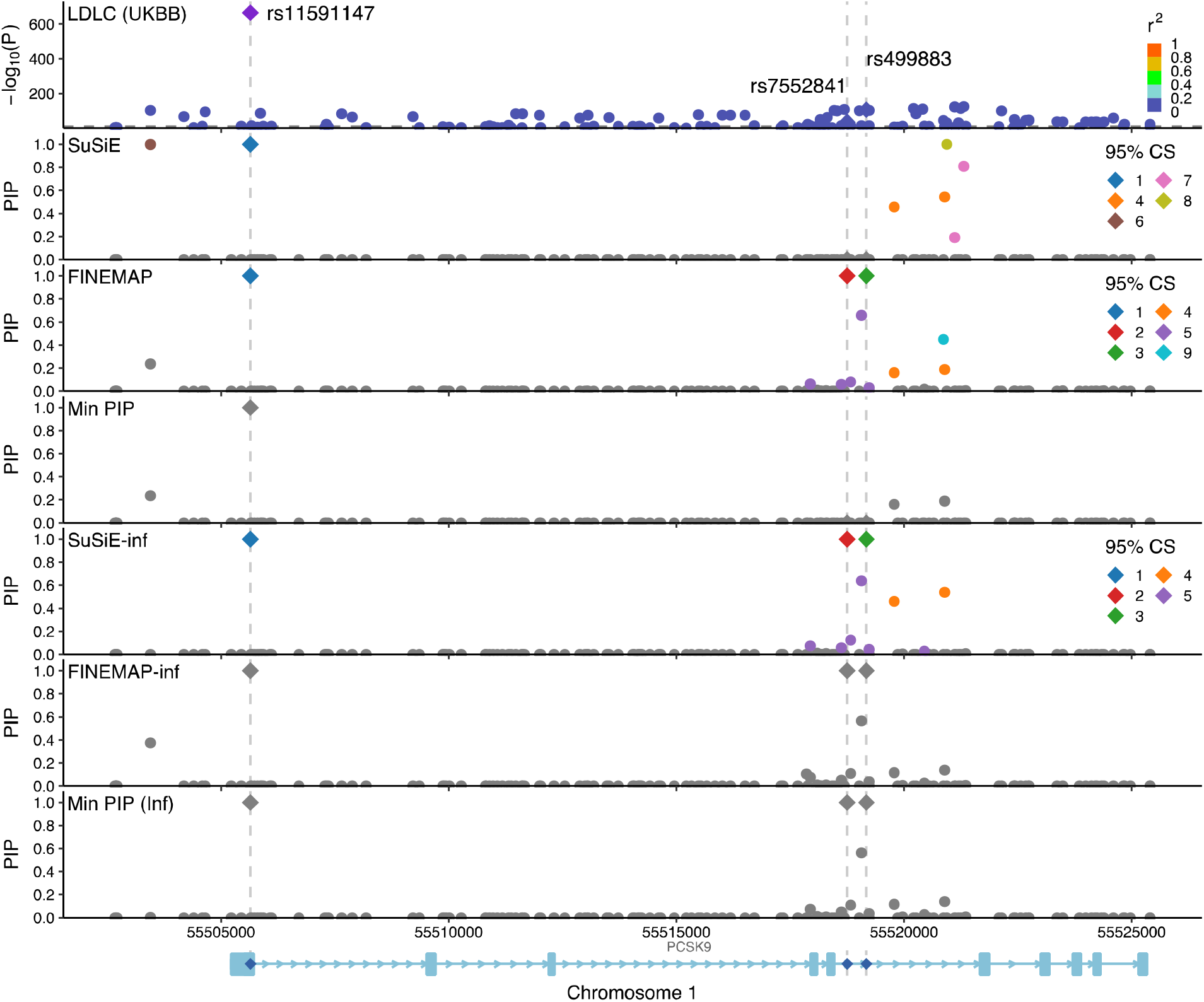
PCSK9 locus for LDLC. 23kbp window at the PCSK9 gene location is shown on the plot. GWAS -log10 P-values for trait LDLC are plotted on the top panel, PIPs from 4 fine-mapping methods and 2 aggregating methods are plotted on the subsequent panels. The well-known putative causal variant rs11591147 is highlighted with dash line, as well as two intronic variants: rs499883 and rs7552841.

**Extended Data Fig. 10.**
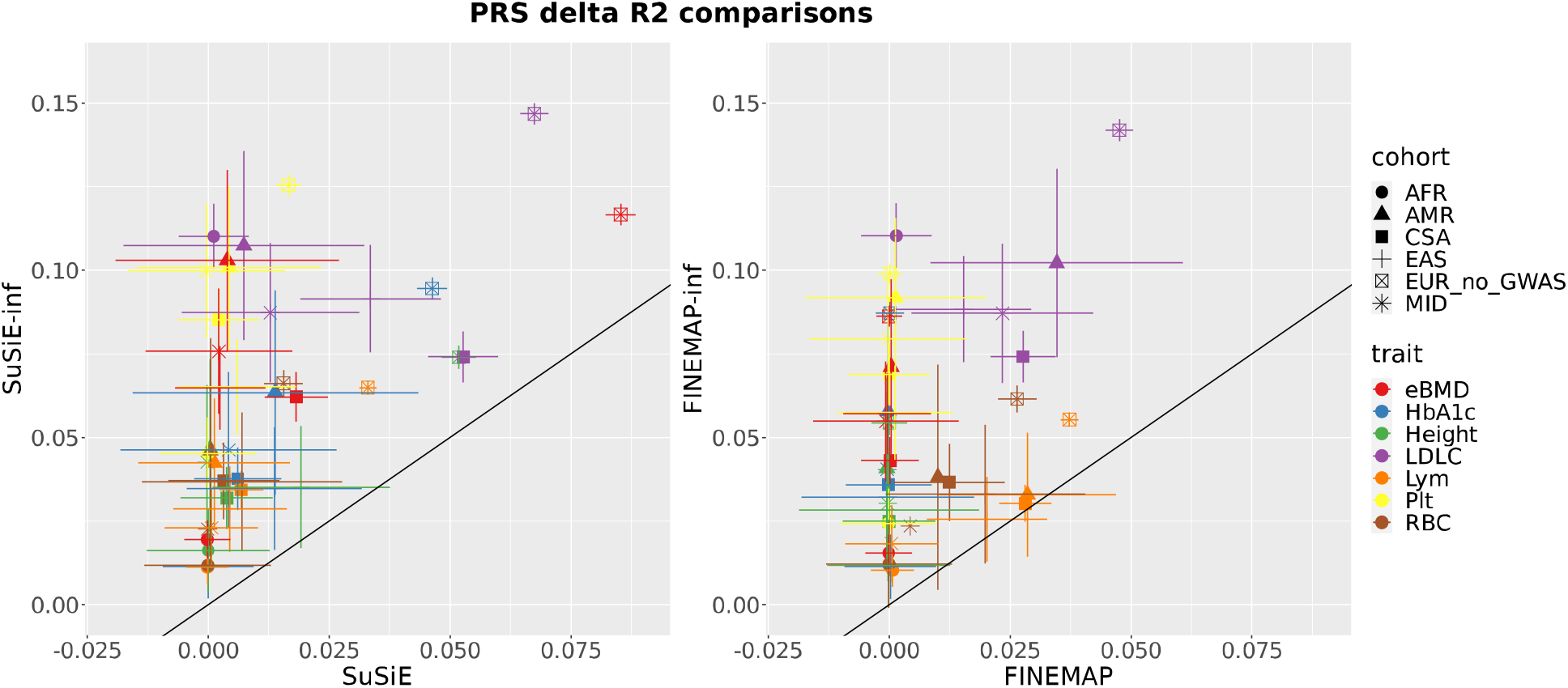
PRS comparison with standard error. Point values are the same as 790 **Fig. 5**, standard errors are computed using R function CI.rsq. Numerical results available in 791 **Supplementary Table 12**.

## References

1. Visscher, P. M. et al. 10 Years of GWAS Discovery: Biology, Function, and Translation. Am. J. Hum. Genet. 101, 5–22 (2017).

2. Shendure, J., Findlay, G. M. & Snyder, M. W. Genomic Medicine-Progress, Pitfalls, and Promise. Cell 177, 45–57 (2019).

3. Hukku, A. et al. Probabilistic colocalization of genetic variants from complex and molecular traits: promise and limitations. Am. J. Hum. Genet. 108, 25–35 (2021).

4. Hormozdiari, F. et al. Colocalization of GWAS and eQTL Signals Detects Target Genes. Am. J. Hum. Genet. 99, 1245–1260 (2016).

5. LaPierre, N. et al. Identifying causal variants by fine mapping across multiple studies. PLoS Genet. 17, e1009733 (2021).

6. Kichaev, G. & Pasaniuc, B. Leveraging Functional-Annotation Data in Trans-ethnic Fine-Mapping Studies. Am. J. Hum. Genet. 97, 260–271 (2015).

7. Kichaev, G. et al. Integrating functional data to prioritize causal variants in statistical fine-mapping studies. PLoS Genet. 10, e1004722 (2014).

8. Wang, G., Sarkar, A., Carbonetto, P. & Stephens, M. A simple new approach to variable selection in regression, with application to genetic fine mapping. J. R. Stat. Soc. Ser. B Stat. Methodol. 82, 1273–1300 (2020).

9. Benner, C. et al. FINEMAP: efficient variable selection using summary data from genomewide association studies. Bioinforma. Oxf. Engl. 32, 1493–1501 (2016).

10. Benner, C., Havulinna, A. S., Salomaa, V., Ripatti, S. & Pirinen, M. Refining finemapping: effect sizes and regional heritability. (2018) doi:10.1101/318618.

11. Wakefield, J. Bayes factors for genome-wide association studies: comparison with Pvalues. Genet. Epidemiol. 33, 79–86 (2009).

12. Yang, J. et al. Conditional and joint multiple-SNP analysis of GWAS summary statistics identifies additional variants influencing complex traits. Nat. Genet. 44, 369–375, S1-3 (2012).

13. Trubetskoy, V. et al. Mapping genomic loci implicates genes and synaptic biology in schizophrenia. Nature 604, 502–508 (2022).

14. Mahajan, A. et al. Fine-mapping type 2 diabetes loci to single-variant resolution using high-density imputation and islet-specific epigenome maps. Nat. Genet. 50, 1505–1513 (2018).

15. Locke, A. E. et al. Genetic studies of body mass index yield new insights for obesity biology. Nature 518, 197–206 (2015).

16. Ulirsch, J. C. et al. Interrogation of human hematopoiesis at single-cell and single-variant resolution. Nat. Genet. 51, 683–693 (2019).

17. Westra, H.-J. et al. Fine-mapping and functional studies highlight potential causal variants for rheumatoid arthritis and type 1 diabetes. Nat. Genet. 50, 1366–1374 (2018).

18. Zhang, Z. et al. Genetic analyses support the contribution of mRNA N6-methyladenosine (m6A) modification to human disease heritability. Nat. Genet. 52, 939–949 (2020).

19. Schaid, D. J., Chen, W. & Larson, N. B. From genome-wide associations to candidate causal variants by statistical fine-mapping. Nat. Rev. Genet. 19, 491–504 (2018).

20. Ulirsch & Kanai et al. An annotated atlas of causal variants for complex human traits. Rev.

21. Howrigan, D. P. et al. Nealelab/UK_Biobank_GWAS: v2. (2023) doi:10.5281/zenodo.8011558.

22. Loh, P.-R. et al. Efficient Bayesian mixed-model analysis increases association power in large cohorts. Nat. Genet. 47, 284–290 (2015).

23. Kanai, M. et al. Meta-analysis fine-mapping is often miscalibrated at single-variant resolution. Cell Genomics 2, 100210 (2022).

24. Zhou, X., Carbonetto, P. & Stephens, M. Polygenic modeling with bayesian sparse linear mixed models. PLoS Genet. 9, e1003264 (2013).

25. Vilhjálmsson, B. J. et al. Modeling Linkage Disequilibrium Increases Accuracy of Polygenic Risk Scores. Am. J. Hum. Genet. 97, 576–592 (2015).

26. Erbe, M. et al. Improving accuracy of genomic predictions within and between dairy cattle breeds with imputed high-density single nucleotide polymorphism panels. J. Dairy Sci. 95, 4114–4129 (2012).

27. Chatterjee, N., Shi, J. & García-Closas, M. Developing and evaluating polygenic risk prediction models for stratified disease prevention. Nat. Rev. Genet. 17, 392–406 (2016).

28. Márquez-Luna, C., Loh, P.-R., South Asian Type 2 Diabetes (SAT2D) Consortium, SIGMA Type 2 Diabetes Consortium & Price, A. L. Multiethnic polygenic risk scores improve risk prediction in diverse populations. Genet. Epidemiol. 41, 811–823 (2017).

29. Pan UKBB | Pan UKBB. https://pan.ukbb.broadinstitute.org/.

30. Kanai, M. et al. Insights from complex trait fine-mapping across diverse populations. 2021.09.03.21262975 Preprint at 10.1101/2021.09.03.21262975 (2021).

31. Weissbrod, O. et al. Functionally informed fine-mapping and polygenic localization of complex trait heritability. Nat. Genet. 52, 1355–1363 (2020).

32. Lloyd-Jones, L. R. et al. Improved polygenic prediction by Bayesian multiple regression on summary statistics. Nat. Commun. 10, 5086 (2019).

33. Bates, S., Candès, E., Janson, L. & Wang, W. Metropolized Knockoff Sampling. J. Am. Stat. Assoc. 116, 1413–1427 (2021).

34. Candès, E., Fan, Y., Janson, L. & Lv, J. Panning for Gold: ‘Model-X’ Knockoffs for High Dimensional Controlled Variable Selection. J. R. Stat. Soc. Ser. B Stat. Methodol. 80, 551–577 (2018).

35. Sesia, M., Katsevich, E., Bates, S., Candès, E. & Sabatti, C. Multi-resolution localization of causal variants across the genome. Nat. Commun. 11, 1093 (2020).

36. Sesia, M., Bates, S., Candès, E., Marchini, J. & Sabatti, C. False discovery rate control in genome-wide association studies with population structure. Proc. Natl. Acad. Sci. U. S. A. 118, e2105841118 (2021).

37. He, Z. et al. GhostKnockoff inference empowers identification of putative causal variants in genome-wide association studies. Nat. Commun. 13, 7209 (2022).

38. Benner, C. et al. Prospects of Fine-Mapping Trait-Associated Genomic Regions by Using Summary Statistics from Genome-wide Association Studies. Am. J. Hum. Genet. 101, 539–551 (2017).

39. Schoech, A. P. et al. Quantification of frequency-dependent genetic architectures in 25 UK Biobank traits reveals action of negative selection. Nat. Commun. 10, 790 (2019).

40. Finucane, H. K. et al. Partitioning heritability by functional annotation using genome-wide association summary statistics. Nat. Genet. 47, 1228–1235 (2015).

41. Hail | Index. https://hail.is/.

42. Chang, C. C. et al. Second-generation PLINK: rising to the challenge of larger and richer datasets. GigaScience 4, 7 (2015).

43. Weissbrod, O. et al. Leveraging fine-mapping and multipopulation training data to improve cross-population polygenic risk scores. Nat. Genet. 54, 450–458 (2022).

