## Supplementary Note for "Improving fine-mapping by modeling infinitesimal effects"

**C RFR and false discovery rate** **4**

**D EPN under a correctly specified model** **4**

**E FINEMAP-inf and SuSiE-inf method descriptions** **5**
E.1 FINEMAP-inf . . . 5
E.2 SuSiE-inf . . . 11

**F Variants in Low Complexity Regions (LCR)** **18**

**G Figures** **19**

### Supplementary Note

#### A Comparison with ABF and COJO-ABF

Approximate Bayes' factors (ABF[1]) and conditional and joint analysis followed by ABF (COJO[2]-ABF) are two commonly used Bayesian fine-mapping methods. ABF is a single-causal-variant fine-mapping method where only one causal variant is modeled in a given fine-mapped region. It is most commonly applied when only summary statistics are available since it does not require LD. COJO-ABF first uses conditional analysis to infer independent associations, then performs ABF on each identified association, conditioning on the others. This approach is not model-based, unlike SuSiE and FINEMAP.

COJO-ABF has RFR 11% in real data, 2.6% RFR in ideal simulations, and a similar functional enrichment profile to SuSiE and FINEMAP (**Supplementary Fig. 1**). However, we observed severe miscalibration of COJO-ABF in our ideal simulations and lower recall than the other model-based multiple-causal-variant fine-mapping methods we tested (**Supplementary Fig. 2a-b**). This is consistent with existing literature showing that conditional analysis has suboptimal precision and power/sensitivity[3, 5]. We observed that the regions with multiple causal variants are more likely to harbor false positive variants (**Supplementary Fig. 2c**) a possible reason that previous simulations with fewer causal variants did not show such severe miscalibration[3].

ABF has a relatively higher RFR of 17%, however functional annotations of the non-replicating variants show comparable enrichment to the replicated variants (**Supplementary Fig. 1b**). ABF also has higher RFR in ideal simulations (12%), where 70% of the non-replicating variants are true causal variants. Based on the evidence from both real data and simulations, we hypothesize that the high RFR for ABF in real data is mostly due to different causal variants being prioritized at different sample sizes and it mostly reflects the non-discovery rate of single-causal-variant fine-mapping in a locus with multiple causal variants, instead of a high false discovery rate (**Supplementary Note section C**). As expected, ABF exhibits much lower recall than the multiple-causal-variant methods (**Supplementary Fig. 2b**).

We recommend using model-based multiple-causal-variant methods for fine-mapping when in-sample LD or genotype/phenotype data are available for better calibration and recall.

#### B Investigating non-replication

##### B.1 BOLT-LMM v.s. OLS

We investigated whether the use of BOLT-LMM summary statistics can induce non-replication in downstream fine-mapping, by comparing results with those obtained using summary statistics computed by ordinary least squares (OLS). We performed GWAS using OLS on Height at N = 366K and N = 100K. We obtained near linear (correlation coefficient 0.95) relations between the OLS marginal association chi-squared statistics and the BOLT-LMM chi-squared statistics, with BOLT-LMM chi-squared statistics being larger (**Supplementary Fig. 3a-b**). We observed decreased RFR (SuSiE: 20% $\rightarrow$ 8% $\pm$ 5%, FINEMAP: 28% $\rightarrow$ 6% $\pm$ 5%), but also substantially reduced power, when using OLS summary statistics. To approximately match the OLS analyses on power, we re-performed analyses using BOLT-LMM and SuSiE (omitting other methods for due to computational cost concerns) with reduced sample and subsample sizes of N = 280K and N = 88K (**Supplementary Fig. 3c-d**). We chose these sample sizes based on the comparison of the chi-squared statistics: the OLS chi-squared statistics (squared z scores) are roughly proportional to the sample size and the slope in **Supplementary Fig. 3a** is 1.3, so we chose 366K/1.3$\approx$280K as the sample size where BOLT would have similar power as OLS. We chose 88K by a similar analysis of **Supplementary Fig. 3b**. In these analyses we observed an RFR reduction (SuSiE: 20% $\rightarrow$ 9%$\pm$5%) similar to the reduction observed using OLS. We conclude that the power difference between BOLT-LMM and OLS, rather than the difference between statistical models, was likely the main contributor to the RFR differences in this investigation.

#### B.2 Region definition differences between sample sizes

In the fine-mapping pipeline of[4], individual fine-mapped regions are defined by windows around genome-wide-significant variants then merging overlapping regions, and regions defined at N = 366K are often larger (because of increased power and the merging of overlapping regions) than those defined at N = 100K. We investigated whether potentially missing causal variants due to differences in region definition can contribute to non-replication. We re-applied SuSiE for Height at N = 100K using the regions defined at N = 366K, for 35 regions that harbored either non-replicating or replicated variants. We observed only one fewer (12 $\rightarrow$ 11) non-replicating variant when using the same region definitions at both sample sizes, suggesting that region definition differences are unlikely to be a main contributor to non-replication. We note that using an alternative pipeline described here[7] where the same sets of 3Mb sliding windows are used to perform fine-mapping at different sample sizes, we also observed high levels of RFR (**Supplementary Fig. 4**).

#### B.3 Maximum number of causal variants per region

In our analyses, for both SuSiE and FINEMAP, we set the maximum number of causal variants per region to be 10. We investigated whether increasing this number would reduce RFR that is potentially caused by different prioritizations of true causal variants at different sample sizes. We re-applied SuSiE on the 35 regions for Height defined above, setting the maximum number of causal variants to 20 and 50. Both settings yielded the exact same high-PIP and non-replicating variants, and the RFRs were 23% $\pm$ 6.5%, no improvement from 20%.

#### B.4 PC differences between sample sizes

We considered the possibility that PC differences at different sample sizes can potentially introduce uncorrected (or differently corrected) confounding, and therefore lead to inconsistency in fine-mapping results. We re-applied SuSiE on Height using the PCs computed at N = 366K as covariates when performing GWAS at N = 100K. The resulting RFR is 27% $\pm$ 6.5%, similar to the original RFR=20%$\pm$ 5%, which is what we observed when using PCs computed at N = 100K. We therefore rule out this possibility.

#### B.5 SNP properties

To further investigate the non-replicating variants, we attempted to characterize non-replicating and replicated variants using the following properties: (a) Minor Allele Frequency (MAF), (b) imputation INFO score, (c) chi-square statistics, (d) LD score, (e) value of the PIP, (f) SuSiE and FINEMAP PIP difference, (g) posterior expected number of other causal variants within a 100kb window. We measured the ability of each property for distinguishing non-replicating and replicated variants by how well a simple threshold rule using its value can separate these classes, as is commonly done to measure feature importance in binary classification. We found that none of these properties can lead to an effective threshold-based QC process that reduces RFR without significantly compromising power (**Supplementary Fig. 5a**).

#### B.6 Distribution of non-replicating variants under repeated subsampling

We investigated whether or not most of the non-replication may be attributed to a small number of non-representative regions. To do this, we repeated our analyses for Height using SuSiE and FINEMAP in 10 additional randomly downsampled subsets of N = 100K individuals. We call a region non-replicating if it harbors any non-replicating variant(s). Out of 88 non-replicating regions that harbored a total of 193 non-replicating variants across both methods and all 10 downsampling analyses, only 19 (22%) regions were non-replicating in more than 2 out of 10 downsampling analyses, and only 5 (6%) regions were non-replicating in more than 5 downsampling analyses. In comparison, regions containing replicated variants tended to repeatedly appear in multiple downsampling analyses (**Supplementary Fig. 5b**). We conclude that non-replication is not mainly due to complexities in a few non-representative loci.

### C RFR and false discovery rate

In this section, we discuss the relation between our Replication Failure Rate (RFR) and the false discovery rate in the $N = 100$K subsample.

Let $\text{PIP}^{366K}$ and $\text{PIP}^{100K}$ be the PIPs for a randomly chosen variant in the $N = 366$K sample and the $N = 100$K subsample. Let $\gamma = 1$ if this chosen variant is causal, and $\gamma = 0$ otherwise. The RFR is an estimate of $\Pr(\text{PIP}^{366K} \leq 0.1 \mid \text{PIP}^{100K} \geq 0.9)$.

Using the law of total probability,

$$
\begin{aligned}
&\Pr(\text{PIP}^{366K} \leq 0.1 \mid \text{PIP}^{100K} \geq 0.9) \\
&\quad = \Pr(\text{PIP}^{366K} \leq 0.1 \mid \gamma = 0, \text{PIP}^{100K} \geq 0.9) \cdot \Pr(\gamma = 0 \mid \text{PIP}^{100K} \geq 0.9) \\
&\qquad + \Pr(\text{PIP}^{366K} \leq 0.1 \mid \gamma = 1, \text{PIP}^{100K} \geq 0.9) \cdot \Pr(\gamma = 1 \mid \text{PIP}^{100K} \geq 0.9) \\
&\quad \leq \Pr(\gamma = 0 \mid \text{PIP}^{100K} \geq 0.9) + \Pr(\text{PIP}^{366K} \leq 0.1 \mid \gamma = 1, \text{PIP}^{100K} \geq 0.9)
\end{aligned}
$$

The first term $\Pr(\gamma = 0 \mid \text{PIP}^{100K} \geq 0.9)$ is the false discovery rate in the $N = 100$K subsample. The second term represents a "non-discovery rate": the probability of non-discovery at 366K, given that the variant is causal and is detected with high-confidence at 100K. Assuming that this non-discovery rate is negligible, the quantity $\Pr(\text{PIP}^{366K} \leq 0.1 \mid \text{PIP}^{100K} \geq 0.9)$ estimated by RFR provides a lower bound to the false discovery rate $\Pr(\gamma = 0 \mid \text{PIP}^{100K} \geq 0.9)$.

In our ideal simulations without model misspecification, we observed the non-discovery rate to be nearly 0 for the multiple-causal-variant methods SuSiE, FINEMAP and COJO-ABF, as may be expected since the methods are more well-powered at the larger 366K sample size to detect true causal variants. The single-causal-variant method ABF showed a higher non-discovery rate of 9%, due to the method sometimes prioritizing different causal variants at different sample sizes in loci with more than one true causal variant.

### D EPN under a correctly specified model

We refer to the false discovery rate $\Pr(\gamma = 0 \mid \text{PIP}^{100K} \geq 0.9)$ for a correctly specified Bayesian model and an exact posterior inference procedure as the Expected Proportion of Non-causal variants (EPN). This must be in the range 0–10%, and its exact value depends on the true effect size distribution. This value is equivalently given by

$$\text{EPN} = 1 - \mathbb{E}[\text{PIP}^{100K} \mid \text{PIP}^{100K} \geq 0.9] \tag{1}$$

To verify this form, note that under a correctly specified model, by definition of the PIP, we have

$$\text{PIP}^{100K} = \Pr(\gamma = 1 \mid \text{data}^{100K}) = 1 - \mathbb{E}[\mathbb{1}\{\gamma = 0\} \mid \text{data}^{100K}]$$

Then by the tower property of conditional expection,

$$
\begin{aligned}
\mathbb{E}[(1 - \text{PIP}^{100K})\mathbb{1}\{\text{PIP}^{100K} \geq 0.9\}] &= \mathbb{E}\big[\mathbb{E}[\mathbb{1}\{\gamma = 0\} \mid \text{data}^{100K}]\mathbb{1}\{\text{PIP}^{100K} \geq 0.9\}\big] \\
&= \Pr(\gamma = 0, \text{PIP}^{100K} \geq 0.9)
\end{aligned}
$$

So by Bayes' rule,

$$
\begin{aligned}
\text{EPN} = \Pr(\gamma = 0 \mid \text{PIP}^{100K} \geq 0.9) &= \frac{\Pr(\gamma = 0, \text{PIP}^{100K} \geq 0.9)}{\Pr(\text{PIP}^{100K} \geq 0.9)} \\
&= \frac{\mathbb{E}[(1 - \text{PIP}^{100K})\mathbb{1}\{\text{PIP}^{100K} \geq 0.9\}]}{\Pr(\text{PIP}^{100K} \geq 0.9)} = \mathbb{E}[1 - \text{PIP}^{100K} \mid \text{PIP}^{100K} \geq 0.9]
\end{aligned}
$$

yielding the form (1).

Under these assumptions, we may thus estimate EPN from the observed data by the empirical average

$$1 - \frac{1}{\#\{j \mid \text{PIP}_j^{100K} \geq 0.9\}} \sum_{j \mid \text{PIP}_j^{100K} \geq 0.9} \text{PIP}_j^{100K}, \tag{2}$$

which is 1 minus the mean PIP in the range [0.9,1] for the 100K subsample. This estimate of EPN is depicted in **[Fig. 1]**

To summarize, for a correctly specified Bayesian model and an exact posterior inference procedure, we expect the RFR to estimate a lower bound to the EPN plus non-discovery rate. In real data for SuSiE, we estimated RFR = 15% and EPN = 2% across the 10 UK Biobank phenotypes. Our ideal simulations suggest that the non-discovery rate is unlikely to exceed the observed difference between RFR and EPN of 13%. Instead, we believe that the high observed RFR is likely indicative of model misspecification and method miscalibration.

### E FINEMAP-inf and SuSiE-inf method descriptions

We describe in this section the details of the FINEMAP-inf and SuSiE-inf methods, to provide a complete and self-contained documentation of these procedures. The models and computational ideas closely follow those of FINEMAP [6] and SuSiE [13], with certain linear-algebraic computations re-expressed via an initial eigen-decomposition of the LD matrix to reduce computational cost when modeling the infinitesimal component. Related computational strategies have been previously used in mixed-model analyses in [10, 11, 14]. We choose to use the same prior specifications and parameter settings of the original FINEMAP and SuSiE methods where possible, so that differences in results may be clearly attributed to the addition of the infinitesimal components of these models.

#### E.1 FINEMAP-inf

**Model.** FINEMAP-inf is based on a Bayesian linear model

$$y = X\beta + X\alpha + \varepsilon \in \mathbb{R}^n \tag{3}$$

with independent coordinates

$$\beta_j \overset{iid}{\sim} \pi_0 \cdot \mathcal{N}(0, s^2) + (1-\pi_0)\cdot\delta_0, \qquad \alpha_j \overset{iid}{\sim} \mathcal{N}(0,\tau^2), \qquad \varepsilon_i \overset{iid}{\sim} \mathcal{N}(0,\sigma^2).$$

Here, $\alpha, \beta \in \mathbb{R}^p$ are the infinitesimal and sparse components of the genetic effects, $\pi_0 \in (0,1)$ is the prior probability that each SNP is causal in the sparse component, $s^2$ is the effect size variance of the sparse effects, $\tau^2$ is the effect size variance of the infinitesimal effects, and $\sigma^2$ is the noise variance.

Note that this is equivalent to a model in which the combined genetic effect $\beta + \alpha \in \mathbb{R}^p$ has coordinates following a two-component normal mixture distribution $\pi_0 \cdot \mathcal{N}(0, s^2+\tau^2) + (1-\pi_0)\cdot\mathcal{N}(0,\tau^2)$. However, it will be convenient for computation to parametrize the sparse and infinitesimal effects separately as in Eq. (3), and to compute Bayes factors by first marginalizing over $(\alpha, \varepsilon)$ followed by the effect sizes of $\beta$.

FINEMAP-inf fixes a given prior causal probability $\pi_0$ and a discrete uniform hyperprior for $s^2$,

$$s^2 \sim \text{Uniform}(\mathcal{S})$$

where $\mathcal{S}$ is a pre-specified set of possible effect size variances. The variance parameters $(\sigma^2, \tau^2)$ are assumed unknown, and empirical Bayes point estimates of these parameters are provided by the method.

**Method.** Let us represent the point-normal distribution for $\beta_j$ as

$$\beta_j = \mu_j \cdot \gamma_j, \qquad \mu_j \sim \mathcal{N}(0, s^2), \qquad \gamma_j \sim \text{Bernoulli}(\pi_0).$$

The binary indicator $\gamma_j \in \{0,1\}$ indicates whether SNP $j$ is "causal", and $\mu_j$ is the signed effect size. The method returns estimates of $(\sigma^2, \tau^2)$ for this locus, together with approximations of the following quantities for each SNP $j \in \{1,\ldots,p\}$:

- $\text{PIP}_j = \Pr[\gamma_j = 1 \mid X, y]$ the posterior-inclusion-probability.
- $b_j|\text{causal} = \mathbb{E}[\beta_j \mid \gamma_j = 1,\, X, y]$ the posterior mean of the sparse effect $\beta_j$ conditional on inclusion.

- $s_j|\text{causal} = \sqrt{\text{Var}[\beta_j \mid \gamma_j = 1, X, y]}$ the posterior standard deviation of $\beta_j$ conditional on inclusion.
- $a_j = \mathbb{E}[\alpha_j \mid X, y]$ the posterior mean infinitesimal effect.

The posterior mean sparse effect not conditional on inclusion is then $b_j = \mathbb{E}[\beta_j \mid X, y] = \text{PIP}_j \cdot b_j|\text{causal}$, and the total posterior mean genetic effect of SNP $j$ is $b_j + a_j$.

Denote the non-zero effects of $\beta$ as the "causal model"

$$\Gamma = \{j : \gamma_j = 1\} \subseteq \{1, \ldots, p\}.$$

FINEMAP-inf alternates between

1. Performing iterations of shotgun-stochastic-search (SSS) to explore and compute Bayes factors for candidate causal models, and
2. Updating $(\sigma^2, \tau^2)$ and the Bayes factors of causal models already explored.

The schedule of total SSS iterations may thus be divided into $K$ epochs $[T_1, \ldots, T_K]$, where $T_k$ is the number of SSS iterations performed in the $k^{\text{th}}$ epoch, and $(\sigma^2, \tau^2)$ are updated $K - 1$ times, once between every two consecutive epochs.

*Shotgun stochastic search.* The SSS procedure is similar to FINEMAP, and we review this here for completeness. Let $\emptyset$ denote the empty model where every coordinate of $\beta$ is 0. Each iteration of SSS starts with an active causal model $\Gamma$, with $\Gamma = \emptyset$ for the first iteration. The neighborhood $N(\Gamma)$ of $\Gamma$ is defined as all models $\Gamma'$ that add, remove, or swap a single SNP in $\Gamma$, and that have at most $L$ total SNPs for a user-specified number $L$. For each neighboring model $\Gamma' \in N(\Gamma)$, SSS computes the Bayes factor

$$q(\Gamma') = \frac{\text{Pr}[\text{causal model is } \Gamma' \mid X, y]}{\text{Pr}[\text{causal model is } \emptyset \mid X, y]} \tag{4}$$

and adds $\Gamma'$ to a list $\mathcal{L}$ of all explored models. The active model $\Gamma'$ for the next SSS iteration is then selected randomly from $N(\Gamma)$, where each $\Gamma' \in N(\Gamma)$ is selected with probability proportional to $q(\Gamma')$.

The Bayes factor $q(\Gamma')$ is proportional to the posterior probability that $\Gamma'$ is the causal model, up to a multiplicative constant. Normalizing by $\sum_{\Gamma \in \mathcal{L}} q(\Gamma)$, the sum across all explored models $\Gamma$, will provide an approximation to this posterior probability that is increasingly accurate as SSS explores an increasingly large subset of the model space.

*Updates for* $(\sigma^2, \tau^2)$*.* Between epochs of SSS, $(\sigma^2, \tau^2)$ are updated using a method-of-moments procedure, based on the moment identities

$$\mathbb{E}\Big[\mathbb{E}[\|y - X\beta\|_2^2 \mid X, y]\Big] = \mathbb{E}[\|y - X\beta\|_2^2] = \mathbb{E}[\|X\alpha + \varepsilon\|_2^2] = \sigma^2 n + \tau^2 \operatorname{Tr} X^\top X, \tag{5}$$

$$\mathbb{E}\Big[\mathbb{E}[\|X^\top(y - X\beta)\|_2^2 \mid X, y]\Big] = \mathbb{E}[\|X^\top(y - X\beta)\|_2^2] = \mathbb{E}[\|X^\top(X\alpha + \varepsilon)\|_2^2] = \sigma^2 \operatorname{Tr} X^\top X + \tau^2 \operatorname{Tr}(X^\top X)^2 \tag{6}$$

derived from the FINEMAP-inf model of Eq. (3). (Related method-of-moments approaches have been used in Haseman-Elston regression and its multi-component generalizations for models without the sparse genetic component $\beta$ [9, 12].) Let $(\sigma^2, \tau^2)$ denote their current values, and let $\mathcal{L}$ be the current list of all models explored by SSS. FINEMAP-inf approximates the posterior expectations on the left sides of Eqs. (5–6) by

$$\mathbb{E}[\|y - X\beta\|_2^2 \mid X, y] \approx m_1 := \frac{\sum_{\Gamma \in \mathcal{L}} \mathbb{E}[\|y - X\beta\|_2^2 \mid \Gamma, X, y] \cdot q(\Gamma)}{\sum_{\Gamma \in \mathcal{L}} q(\Gamma)} \tag{7}$$

$$\mathbb{E}[\|X^\top(y - X\beta)\|_2^2 \mid X, y] \approx m_2 := \frac{\sum_{\Gamma \in \mathcal{L}} \mathbb{E}[\|X^\top(y - X\beta)\|_2^2 \mid \Gamma, X, y] \cdot q(\Gamma)}{\sum_{\Gamma \in \mathcal{L}} q(\Gamma)}. \tag{8}$$

Here, the true posterior distribution over all causal models $\Gamma$ is approximated by the distribution over only explored models $\Gamma \in \mathcal{L}$, and $m_1$ and $m_2$ represent posterior expectations of $\|y - X\beta\|^2$ and $\|X^\top(y - X\beta)\|^2$ computed by first conditioning on $\Gamma$, and then marginalizing over this approximate posterior law for $\Gamma$.

The values of $(\sigma^2, \tau^2)$ are then updated as

$$\begin{pmatrix} \sigma^2_{\text{new}} \\ \tau^2_{\text{new}} \end{pmatrix} = \begin{pmatrix} n & \operatorname{Tr} X^\top X \\ \operatorname{Tr} X^\top X & \operatorname{Tr}(X^\top X)^2 \end{pmatrix}^{-1} \begin{pmatrix} m_1 \\ m_2 \end{pmatrix} \tag{9}$$

This corresponds to substituting $(m_1, m_2)$ for the left sides of the moment equations Eqs. (5–6) and then solving the resulting linear system for $(\sigma^2, \tau^2)$. For loci where the resulting estimate $\sigma^2_{\text{new}}$ or $\tau^2_{\text{new}}$ is negative, we constrain $\tau^2_{\text{new}} = 0$ and use instead the update $(\sigma^2_{\text{new}}, \tau^2_{\text{new}}) = (n^{-1}m_1, 0)$. After updating $(\sigma^2, \tau^2)$ to $(\sigma^2_{\text{new}}, \tau^2_{\text{new}})$, the Bayes factor $q(\Gamma)$ is re-computed for each explored model $\Gamma \in \mathcal{L}$ with these new parameters.

*Computing* $\text{PIP}_j$, $b_j|\text{causal}$, $s_j|\text{causal}$, *and* $a_j$. The PIPs and posterior moments of $\beta_j$ conditional on inclusion are approximated (under the final values for $(\sigma^2, \tau^2)$) by

$$\text{PIP}_j = \Pr[\gamma_j = 1 \mid X, y] \approx \frac{\sum_{\Gamma \in \mathcal{L}:\, j \in \Gamma} q(\Gamma)}{\sum_{\Gamma \in \mathcal{L}} q(\Gamma)} \tag{10}$$

$$b_j|\text{causal} = \mathbb{E}[\beta_j \mid \gamma_j = 1,\, X, y] \approx \frac{\sum_{\Gamma \in \mathcal{L}:\, j \in \Gamma} \mathbb{E}[\beta_j \mid \Gamma, X, y] \cdot q(\Gamma)}{\sum_{\Gamma \in \mathcal{L}:\, j \in \Gamma} q(\Gamma)} \tag{11}$$

$$\mathbb{E}[\beta_j^2 \mid \gamma_j = 1,\, X, y] \approx \frac{\sum_{\Gamma \in \mathcal{L}:\, j \in \Gamma} \mathbb{E}[\beta_j^2 \mid \Gamma, X, y] \cdot q(\Gamma)}{\sum_{\Gamma \in \mathcal{L}:\, j \in \Gamma} q(\Gamma)} \tag{12}$$

These again correspond to approximating the true posterior distribution over $\Gamma$ by the distribution over only explored models $\Gamma \in \mathcal{L}$. The standard deviation $s_j|\text{causal}$ is computed from the above approximations as

$$s_j|\text{causal} = \sqrt{\mathbb{E}[\beta_j^2 \mid \gamma_j = 1,\, X, y] - (b_j|\text{causal})^2}. \tag{13}$$

Finally, the posterior mean for $\alpha$ is computed from the above approximations as

$$a = \mathbb{E}[\hat{\alpha}(\beta) \mid X, y], \qquad \hat{\alpha}(\beta) = \mathbb{E}[\alpha \mid \beta, X, y]. \tag{14}$$

Here, $\hat{\alpha}(\beta)$ takes the form of the Best Linear Unbiased Predictor (BLUP) for $\alpha$ given the residuals $y - X\beta$, which is linear in $\beta$. Its posterior mean $a = \mathbb{E}[\hat{\alpha}(\beta) \mid X, y]$ may thus be computed from $\mathbb{E}[\beta \mid X, y] = (\text{PIP}_j \cdot b_j|\text{causal})_{j=1}^p$ and the above approximations for $b_j|\text{causal}$ and $\text{PIP}_j$.

**Parameter settings.** In our analyses, we standardize the phenotype vector $y \in \mathbb{R}^n$ and genotype matrix $X \in \mathbb{R}^{n \times p}$ to have columns of mean 0 and variance 1. We then set $\mathcal{S} = \{0.05^2\}$ corresponding to a fixed prior effect size variance $s^2 = 0.05^2$. We fix the prior causal probability $\pi_0 = 1/p$ where $p$ is the number of SNPs in the locus, so that the prior expected number of causal SNPs is 1. This is more conservative than the prior that we will use for SuSiE and SuSiE-inf, and we have chosen these parameters to match the default prior parameters used by FINEMAP.

We fix $L = 10$ as the maximum number of causal SNPs in the locus. In our reported results, since we apply both FINEMAP-inf and SuSiE-inf, we simply use $(\sigma^2, \tau^2)$ estimated by SuSiE-inf instead of re-estimating these quantities within FINEMAP-inf, to save on computational cost. For applications of FINEMAP-inf alone, we recommend initializing $(\sigma^2, \tau^2) = (1, 0)$ and using a default SSS schedule with $K = 7$ epochs and $[T_1, \ldots, T_K] = [100, 100, 100, 100, 100, 100, 10000]$, corresponding to 100 SSS iterations before each $(\sigma^2, \tau^2)$ update.

In the last epoch of SSS, we set two parameters $T_{\text{hist}} = 100$ and $\text{tol} = 10^{-3}$, and assess convergence using an approach similar to FINEMAP: Let $\mathcal{L}_{T_{\text{hist}}}$ denote the models explored in the last $T_{\text{hist}}$ iterations. Then, after iteration $T_{\text{hist}}$, the algorithm is terminated if

$$\frac{\sum_{\Gamma \in \mathcal{L}_{T_{\text{hist}}}} q(\Gamma)}{\sum_{\Gamma \in \mathcal{L}} q(\Gamma)} < \text{tol},$$

i.e. if the total posterior probability of models explored in the last $T_{\text{hist}}$ iterations constitutes a proportion of the total probability of all explored models that is less than tol.

**Efficient computation.** For computational efficiency, FINEMAP-inf pre-computes the eigen-decomposition

$$X^\top X = VDV^\top \tag{15}$$

from either an input genotype matrix $X$ or an input LD correlation matrix $\text{LD} = \frac{1}{n}X^\top X$, where columns of $V \in \mathbb{R}^{p\times p}$ are the eigenvectors and $D \in \mathbb{R}^{p\times p}$ is the diagonal matrix of eigenvalues. It then pre-computes the rotated and scaled z-scores

$$u = V^\top X^\top y \tag{16}$$

from the above eigenvector matrix $V$ and either an input phenotype vector $y$ or a z-score vector $z = \frac{1}{\sqrt{n}}X^\top y$. All remaining quantities are computed from $V, D, u$ and the mean-squared phenotype $\langle y^2\rangle = \frac{1}{n}y^\top y$.

We describe in the remainder of this section the details of computing the Bayes factors $q(\Gamma)$ in Eq. (4), the posterior moments $\mathbb{E}[\beta_j, \beta_j^2 \mid \Gamma, X, y]$ for $j \in \Gamma$ in Eqs. (11–12), the quantities $\mathbb{E}[\|y - X\beta\|_2^2, \|X^\top(y - X\beta)\|_2^2 \mid \Gamma, X, y]$ needed for method-of-moments estimation of $(\sigma^2, \tau^2)$ in Eqs. (7–8), and the BLUP $\hat{\alpha}(\beta)$ and its posterior expectation in Eq. (14). We conclude with a discussion of hash map data structures that are used by FINEMAP-inf to reduce the computational cost, and a discussion of the total computational complexity.

*Bayes factors.* Fixing $\sigma^2$ and $\tau^2$, the FINEMAP-inf model in Eq. (3) is equivalently written as

$$y = X\beta + \tilde{\varepsilon} \in \mathbb{R}^n, \qquad \tilde{\varepsilon} \sim \mathcal{N}(0, \Omega^{-1}), \qquad \Omega = (\tau^2 XX^\top + \sigma^2 I)^{-1} \tag{17}$$

where $\Omega$ is the precision matrix of the effective residual $\tilde{\varepsilon} = X\alpha + \varepsilon$ that includes the infinitesimal genetic effect. For a candidate causal model $\Gamma$, denote

$$X_\Gamma = (x_j : j \in \Gamma) \in \mathbb{R}^{n\times|\Gamma|}$$

as the genotype matrix for SNPs belonging to $\Gamma$, where $x_j$ is the $j^{\text{th}}$ column of $X$. Conditioning on the causal model being $\Gamma$, and marginalizing over both $\mu_j \sim \mathcal{N}(0, s^2)$ and $s^2 \sim \text{Uniform}(\mathcal{S})$ in the distribution of $\beta_j$, the phenotypes $y$ have the normal mixture distribution

$$y = \sum_{j\in\Gamma} \beta_j x_j + \tilde{\varepsilon} \sim \frac{1}{|\mathcal{S}|}\sum_{s^2\in\mathcal{S}} \mathcal{N}(0, s^2 X_\Gamma X_\Gamma^\top + \Omega^{-1}).$$

So the probability density of $y$ given $\Gamma$ is

$$\Pr[y \mid \Gamma, X] = \frac{1}{|\mathcal{S}|}\sum_{s^2\in\mathcal{S}} (2\pi)^{-n/2} \det\left(s^2 X_\Gamma X_\Gamma^\top + \Omega^{-1}\right)^{-1/2} \cdot \exp\left(-y^\top \left(s^2 X_\Gamma X_\Gamma^\top + \Omega^{-1}\right)^{-1} y/2\right).$$

For the empty model $\Gamma = \emptyset$, this is

$$\Pr[y \mid \emptyset, X] = (2\pi)^{-n/2} \det\left(\Omega^{-1}\right)^{-1/2} \cdot \exp\left(-y^\top \Omega y/2\right).$$

Define

$$z_\Gamma = X_\Gamma^\top \Omega y \in \mathbb{R}^{|\Gamma|}, \qquad \omega_\Gamma(s^2) = X_\Gamma^\top \Omega X_\Gamma + s^{-2} I.$$

Applying the Woodbury identity and a low-rank update rule for the matrix determinant,

$$y^\top \left(s^2 X_\Gamma X_\Gamma^\top + \Omega^{-1}\right)^{-1} y = y^\top \left(\Omega - \Omega X_\Gamma \left(X_\Gamma^\top \Omega X_\Gamma + s^{-2} I\right)^{-1} X_\Gamma^\top \Omega\right) y = y^\top \Omega y - z_\Gamma^\top \omega_\Gamma(s^2)^{-1} z_\Gamma$$
$$\det\left(s^2 X_\Gamma X_\Gamma^\top + \Omega^{-1}\right) = \det(s^2 X_\Gamma^\top \Omega X_\Gamma + I) \cdot \det(\Omega^{-1}) = \det(s^2 \omega_\Gamma(s^2)) \cdot \det(\Omega^{-1}).$$

Then, canceling factors of $(2\pi)^{-n/2}\det(\Omega^{-1})^{-1/2}$ and $\exp(-y^\top \Omega y/2)$, the Bayes factor for $\Gamma$ is

$$q(\Gamma) = \frac{\Pr[\Gamma \mid X, y]}{\Pr[\emptyset \mid X, y]} = \frac{\Pr[y \mid \Gamma, X]}{\Pr[y \mid \emptyset, X]} \cdot \frac{\Pr[\Gamma]}{\Pr[\emptyset]}$$
$$= \frac{1}{|\mathcal{S}|}\sum_{s^2\in\mathcal{S}} \det(s^2 \omega_\Gamma(s^2))^{-1/2} \cdot \exp\left(z_\Gamma^\top \omega_\Gamma(s^2)^{-1} z_\Gamma/2\right) \cdot \left(\frac{\pi_0}{1-\pi_0}\right)^{|\Gamma|}. \tag{18}$$

In this expression, $z_\Gamma$ and $\omega_\Gamma(s^2)$ may be computed from $V, D, u$ defined in Eqs. (15) and (16) as follows: Applying the Woodbury identity,

$$\begin{aligned} X^\top \Omega X &= \tau^{-2} X^\top \Big(XX^\top + (\sigma^2/\tau^2) I\Big)^{-1} X = \tau^{-2}\Big(I - (I + (\tau^2/\sigma^2) X^\top X)^{-1}\Big) \\ &= \tau^{-2} V \Big(I - (I + (\tau^2/\sigma^2) D)^{-1}\Big) V^\top = V D (\tau^2 D + \sigma^2 I)^{-1} V^\top \end{aligned} \tag{19}$$

Let $V_\Gamma \in \mathbb{R}^{|\Gamma| \times p}$ be the rows of $V$ corresponding to the SNPs of $\Gamma$. Then taking the rows of $\Gamma$ on both sides yields

$$X_\Gamma^\top \Omega X_\Gamma = V_\Gamma D (\tau^2 D + \sigma^2 I)^{-1} V_\Gamma^\top, \tag{20}$$

and adding to this $s^{-2} I$ gives $\omega_\Gamma(s^2)$. Similarly, from the Woodbury identity,

$$\begin{aligned} X^\top \Omega &= \sigma^{-2} X^\top \Big((\tau^2/\sigma^2) XX^\top + I\Big)^{-1} = \sigma^{-2} X^\top \Big(I - X\Big((\sigma^2/\tau^2) I + X^\top X\Big)^{-1} X^\top\Big) \\ &= \sigma^{-2}\Big(I - (X^\top X)\Big((\sigma^2/\tau^2) I + X^\top X\Big)^{-1}\Big) X^\top \\ &= \sigma^{-2} V \Big(I - D\Big((\sigma^2/\tau^2) I + D\Big)^{-1}\Big) V^\top X^\top = V (\tau^2 D + \sigma^2 I)^{-1} V^\top X^\top. \end{aligned} \tag{21}$$

Multiplying on the right by $y$ and taking the rows of $\Gamma$ on both sides yields

$$z_\Gamma = X_\Gamma^\top \Omega y = V_\Gamma (\tau^2 D + \sigma^2 I)^{-1} u. \tag{22}$$

*Posterior moments of $\beta$.* For a candidate causal model $\Gamma$, denote $\beta_\Gamma = (\beta_j : j \in \Gamma)$ as the coordinates of $\beta$ belonging to $\Gamma$. Conditioning on the true causal model being $\Gamma$, and absorbing proportionality factors not depending on $\beta_\Gamma$ into the notation $\propto$, the posterior density for $\beta_\Gamma$ is

$$\begin{aligned} &\Pr[\beta_\Gamma \mid \Gamma, X, y] \propto \Pr[y \mid \beta_\Gamma, \Gamma, X] \cdot \Pr[\beta_\Gamma \mid \Gamma] \\ &\qquad \propto \exp\left(-\frac{1}{2}(y - X_\Gamma \beta_\Gamma)^\top \Omega (y - X_\Gamma \beta_\Gamma)\right) \cdot \frac{1}{|\mathcal{S}|} \sum_{s^2 \in \mathcal{S}} (2\pi s^2)^{-|\Gamma|/2} \exp\left(-\frac{1}{2s^2} \|\beta_\Gamma\|_2^2\right) \\ &\qquad \propto \sum_{s^2 \in \mathcal{S}} (s^2)^{-|\Gamma|/2} \exp\left(-\frac{1}{2} \beta_\Gamma^\top \omega_\Gamma(s^2) \beta_\Gamma + \beta_\Gamma^\top z_\Gamma\right). \end{aligned}$$

Completing the square in the exponent and defining the mixture probability weights over $s^2 \in \mathcal{S}$

$$p(s^2) = \frac{\det(s^2 \omega_\Gamma(s^2))^{-1/2} \exp(z_\Gamma^\top \omega_\Gamma(s^2) z_\Gamma / 2)}{\sum_{s^2 \in \mathcal{S}} \det(s^2 \omega_\Gamma(s^2))^{-1/2} \exp(z_\Gamma^\top \omega_\Gamma(s^2) z_\Gamma / 2)}, \tag{23}$$

this posterior law for $\beta_\Gamma$ is the normal mixture distribution

$$(\beta_\Gamma \mid \Gamma, X, y) \sim \sum_{s^2 \in \mathcal{S}} p(s^2) \cdot \mathcal{N}\Big(\omega_\Gamma(s^2)^{-1} z_\Gamma,\, \omega_\Gamma(s^2)^{-1}\Big). \tag{24}$$

Thus, for each $j \in \Gamma$,

$$\begin{aligned} \mathbb{E}[\beta_j \mid \Gamma, X, y] &= \sum_{s^2 \in \mathcal{S}} p(s^2) \cdot \big(\omega_\Gamma(s^2)^{-1} z_\Gamma\big)_j \\ \mathbb{E}[\beta_j^2 \mid \Gamma, X, y] &= \sum_{s^2 \in \mathcal{S}} p(s^2) \cdot \Big[\big(\omega_\Gamma(s^2)^{-1} z_\Gamma\big)_j^2 + \big(\omega_\Gamma(s^2)^{-1}\big)_{jj}\Big]. \end{aligned}$$

In these expressions, $\omega_\Gamma(s^2)$ and $z_\Gamma$ may again be computed from $V, D, u$ using Eqs. (20) and (22).

*Method-of-moments for $(\sigma^2, \tau^2)$.* Define

$$b_\Gamma = \sum_{s^2 \in \mathcal{S}} p(s^2) \cdot \omega_\Gamma(s^2)^{-1} z_\Gamma, \qquad M_\Gamma = \sum_{s^2 \in \mathcal{S}} p(s^2) \cdot \Big(\omega_\Gamma(s^2)^{-1} z_\Gamma z_\Gamma^\top \omega_\Gamma(s^2)^{-1} + \omega_\Gamma(s^2)^{-1}\Big),$$

so that the mixture distribution of Eq. (24) gives

$$\mathbb{E}[\beta_\Gamma \mid \Gamma, X, y] = b_\Gamma, \qquad \mathbb{E}[\beta_\Gamma \beta_\Gamma^\top \mid \Gamma, X, y] = M_\Gamma.$$

Let $V_\Gamma \in \mathbb{R}^{|\Gamma| \times p}$ be the rows of $V$ corresponding to the SNPs of $\Gamma$. Then $X_\Gamma^\top y = V_\Gamma V^\top X^\top y = V_\Gamma u$, $X_\Gamma^\top X X^\top y = V_\Gamma D V^\top X^\top y = V_\Gamma D u$, $X_\Gamma^\top X_\Gamma = V_\Gamma D V_\Gamma^\top$, and $X_\Gamma^\top X X^\top X_\Gamma = V_\Gamma D V^\top V D V_\Gamma^\top = V_\Gamma D^2 V_\Gamma^\top$. Applying these identities,

$$\begin{aligned}
\mathbb{E}[\beta^\top X^\top y \mid \Gamma, X, y] &= \mathbb{E}[\beta_\Gamma \mid \Gamma, X, y]^\top X_\Gamma^\top y = b_\Gamma^\top V_\Gamma u,\\
\mathbb{E}[\beta^\top (X^\top X) X^\top y \mid \Gamma, X, y] &= \mathbb{E}[\beta_\Gamma \mid \Gamma, X, y]^\top X_\Gamma^\top X X^\top y = b_\Gamma^\top V_\Gamma D u,\\
\mathbb{E}[\beta^\top X^\top X \beta \mid \Gamma, X, y] &= \mathrm{Tr}\left(\mathbb{E}[\beta_\Gamma \beta_\Gamma^\top \mid \Gamma, X, y] \cdot X_\Gamma^\top X_\Gamma\right) = \mathrm{Tr}\, M_\Gamma V_\Gamma D V_\Gamma^\top,\\
\mathbb{E}[\beta^\top (X^\top X)^2 \beta \mid \Gamma, X, y] &= \mathrm{Tr}\left(\mathbb{E}[\beta_\Gamma \beta_\Gamma^\top \mid \Gamma, X, y] \cdot X_\Gamma^\top X X^\top X_\Gamma\right) = \mathrm{Tr}\, M_\Gamma V_\Gamma D^2 V_\Gamma^\top.
\end{aligned}$$

So the posterior expectations needed in the method-of-moments computations of Eqs. (7–8) are given by

$$\begin{aligned}
\mathbb{E}[\|y - X\beta\|_2^2 \mid \Gamma, X, y] &= \|y\|_2^2 - 2\mathbb{E}[\beta^\top X^\top y \mid \Gamma, X, y] + \mathbb{E}[\beta^\top X^\top X \beta \mid \Gamma, X, y]\\
&= n\langle y^2 \rangle - 2 b_\Gamma^\top V_\Gamma u + \mathrm{Tr}\, M_\Gamma V_\Gamma D V_\Gamma^\top, \qquad (25)\\
\mathbb{E}[\|X^\top(y - X\beta)\|_2^2 \mid \Gamma, X, y] &= \|X^\top y\|_2^2 - 2\mathbb{E}[\beta^\top (X^\top X) X^\top y \mid X, y] + \mathbb{E}[\beta (X^\top X)^2 \beta \mid X, y]\\
&= \|u\|_2^2 - 2 b_\Gamma^\top V_\Gamma D u + \mathrm{Tr}\, M_\Gamma V_\Gamma D^2 V_\Gamma^\top. \qquad (26)
\end{aligned}$$

These expressions may be used to compute $(m_1, m_2)$ in Eqs. (7–8). Then $(\sigma^2_{\mathrm{new}}, \tau^2_{\mathrm{new}})$ may be updated as in Eq. (9), using $(m_1, m_2)$ and $\mathrm{Tr}\, X^\top X = \mathrm{Tr}\, D$ and $\mathrm{Tr}(X^\top X)^2 = \mathrm{Tr}\, D^2$.

*BLUP for $\alpha$.* Conditioning on $\beta$ and absorbing proportionality factors not depending on $\alpha$ into $\propto$, the posterior density for $\alpha$ is

$$\begin{aligned}
\Pr[\alpha \mid \beta, X, y] &\propto \Pr[y \mid \alpha, \beta, X] \cdot \Pr[\alpha]\\
&\propto \exp\left(-\frac{1}{2\sigma^2}\|y - X\beta - X\alpha\|_2^2\right) \cdot \exp\left(-\frac{1}{2\tau^2}\|\alpha\|_2^2\right)\\
&\propto \exp\left(-\frac{1}{2\sigma^2\tau^2}\alpha^\top(\tau^2 X^\top X + \sigma^2 I)\alpha + \frac{1}{\sigma^2}\alpha^\top X^\top (y - X\beta)\right).
\end{aligned}$$

This posterior law for $\alpha$ given $\beta$ is a normal distribution with mean

$$\hat{\alpha}(\beta) = \mathbb{E}[\alpha \mid \beta, X, y] = \tau^2(\tau^2 X^\top X + \sigma^2 I)^{-1} X^\top (y - X\beta),$$

which coincides with the BLUP for $\alpha$ given the residuals $y - X\beta$. Applying Eqs. (15) and (16), this is expressed in terms of $V, D, u$ as

$$\hat{\alpha}(\beta) = \tau^2 V(\tau^2 D + \sigma^2 I)^{-1}[u - DV^\top \beta].$$

Then the full posterior mean of $\alpha$ from (14) is

$$a = \mathbb{E}[\hat{\alpha}(\beta) \mid X, y] = \tau^2 V(\tau^2 D + \sigma^2 I)^{-1}[u - DV^\top \mathbb{E}[\beta \mid X, y]]. \qquad (27)$$

Here, $\mathbb{E}[\beta \mid X, y] = (\mathrm{PIP}_j \cdot b_j|\mathrm{causal})_{j=1}^p$, and applying their approximations in Eqs. (10–11) gives the approximation for $a = (a_j)_{j=1}^p$.

*Data structures and computational cost.* To describe the total computational cost of the method, let $\mathcal{L}_k$ be the list of models visited by SSS (i.e. all active models and their neighbors) up to and including epoch $k$, so that $|\mathcal{L}_1| \le \ldots \le |\mathcal{L}_K|$. Let $\mathcal{P}_k$ be the set of distinct SNP pairs $i, j \in \Gamma$ across all models $\Gamma \in \mathcal{L}_k$, i.e. SNP pairs that are jointly causal in any explored model. A trivial upper bound on its size is $|\mathcal{P}_k| \le p^2$.

For each $i = 1, \ldots, p$, let $v_i$ denote the $i^{\text{th}}$ row of $V$. FINEMAP-inf maintains the following five hash maps throughout the computation:

(A) Map from each model $\Gamma$ to the Bayes factor $q(\Gamma)$.

(B) Map from each index $i \in \{1, \ldots, p\}$ to the value of $x_i^\top \Omega y = v_i^\top (\tau^2 D + \sigma^2 I)^{-1} u$.

(C) Map from each index pair $i, j \in \{1, \ldots, p\}$ to the value of $x_i^\top \Omega x_j = v_i^\top D(\tau^2 D + \sigma^2 I)^{-1} v_j$.

(D) Map from each index $i \in \{1, \ldots, p\}$ to the value of $v_i^\top D u$.

(E) Map from each index pair $i, j \in \{1, \ldots, p\}$ to the values of $v_i^\top D v_j$ and $v_i^\top D^2 v_j$.

We will assume that lookup and insertion into these hash maps may be performed in $O(1)$ time. The values in Maps A, B, C depend on $(\sigma^2, \tau^2)$ and are re-computed when $(\sigma^2, \tau^2)$ are updated between epochs. FINEMAP-inf populates all five hash maps by lazy evaluation, i.e. the method does not evaluate $x_i^\top \Omega x_j$ for an index pair $(i, j)$ unless required in the SSS computations. Up to each epoch $k$, this yields a computational savings if $|\mathcal{P}_k|$ is much smaller than $p^2$.

For each model $\Gamma' \in N(\Gamma)$ visited by SSS, FINEMAP-inf retrieves its Bayes factor $q(\Gamma')$ from Map A if available. Otherwise, $q(\Gamma')$ is computed and saved to Map A. In this computation, for every SNP pair $i, j \in \Gamma'$, the values of $x_i^\top \Omega y$ and $x_i^\top \Omega x_j$ are either retrieved from Maps B and C, or computed anew and saved to these maps. Across all models $\Gamma' \in \mathcal{L}_k$, the number of distinct such pairs $i, j \in \Gamma'$ is at most $|\mathcal{P}_k|$, and the cost of computing $x_i^\top \Omega y$ or $x_i^\top \Omega x_j$ anew is $O(p)$ using their above forms in terms of $v_i, v_j, D$. Having computed or retrieved all such values for $i, j \in \Gamma'$, the Bayes factor $q(\Gamma')$ may be computed via Eq. (18) in time $O(|\mathcal{S}|L^3)$. So the total cost to compute $q(\Gamma)$ for all $\Gamma \in \mathcal{L}_k$ is

$$O\Big(|\mathcal{L}_k||\mathcal{S}|L^3 + |\mathcal{P}_k|p\Big).$$

After epoch $k$ of SSS, for each $\Gamma \in \mathcal{L}_k$ and each SNP pair $i, j \in \Gamma$, FINEMAP-inf retrieves the values of $v_i^\top D u$, $v_i^\top D v_j$, and $v_i^\top D^2 v_j$ from Maps D and E, or computes these anew and saves them to these maps. As above, this requires time $O(|\mathcal{P}_k|p)$ across all models $\Gamma \in \mathcal{L}_k$. Given these values, the quantities $b_\Gamma, M_\Gamma$ and $\mathbb{E}[\|y - X\beta\|_2^2, \|X^\top(y - X\beta)\|_2^2 \mid \Gamma, X, y]$ in Eqs. (25–26) may be computed in time $O(|\mathcal{S}|L^3)$ for each model $\Gamma \in \mathcal{L}_k$. Thus, the update of $(\sigma^2_{\text{new}}, \tau^2_{\text{new}})$ using Eq. (9) may be computed also in time

$$O\Big(|\mathcal{L}_k||\mathcal{S}|L^3 + |\mathcal{P}_k|p\Big),$$

and the Bayes factors of all models $q(\Gamma)$ for $\Gamma \in \mathcal{L}_k$ may be re-computed using $(\sigma^2_{\text{new}}, \tau^2_{\text{new}})$ in this same time.

Thus, assuming that the initial eigen-decomposition of $X^\top X$ is computed in time $O(p^3)$, the total computation time of FINEMAP-inf is

$$O\left(p^3 + \sum_{k=1}^{K} |\mathcal{L}_k||\mathcal{S}|L^3 + \sum_{k=1}^{K} |\mathcal{P}_k|p\right).$$

Note that naively recomputing $x_i^\top \Omega x_j$ and $x_i^\top \Omega y$ after each $(\sigma^2, \tau^2)$ update by direct matrix inversion, without precomputing the eigen-decomposition of LD, would instead require time $O(p^3)$, yielding a total complexity of $O(Kp^3 + \sum_{k=1}^{K} |\mathcal{L}_k||\mathcal{S}|L^3)$. For "easy" fine-mapping instances and parameter settings where the numbers of SSS iterations $T_1, \ldots, T_{K-1}$ in all but the last epoch are kept small, we observe that $\mathcal{P}_k$ may contain few index pairs $i, j$ that both do not belong to the true causal model. Hence $|\mathcal{P}_k|$ may be on the order of $Lp$ rather than $p^2$ in such settings, so that $\sum_{k=1}^{K} |\mathcal{P}_k|p$ is smaller than its naive upper bound of $Kp^3$.

#### E.2 SuSiE-inf

**Model.** SuSiE-inf is based on the linear model

$$y = X\beta + X\alpha + \varepsilon \in \mathbb{R}^n, \qquad \beta = \sum_{l=1}^{L} \beta^{(l)} = \sum_{l=1}^{L} \mu^{(l)} \cdot \gamma^{(l)} \tag{28}$$

with independent components

$$\mu^{(l)} \sim \mathcal{N}(0, s_l^2), \qquad \gamma^{(l)} \sim \text{Uniform}(e_1, \ldots, e_p), \qquad \alpha_j \overset{iid}{\sim} \mathcal{N}(0, \tau^2), \qquad \varepsilon_i \overset{iid}{\sim} \mathcal{N}(0, \sigma^2).$$

Here $e_1, \dots, e_p \in \mathbb{R}^p$ denote the standard basis vectors, so each $\gamma^{(l)} \in \mathbb{R}^p$ is the indicator vector of a single uniformly-chosen random SNP, and $\beta^{(l)} = \mu^{(l)} \cdot \gamma^{(l)} \in \mathbb{R}^p$ represents the effect of that SNP with signed effect size $\mu^{(l)}$.

SuSiE-inf fixes a pre-specified number of causal variants $L$. The variance parameters $s_1^2, \dots, s_L^2, \sigma^2, \tau^2$ are all assumed unknown, and empirical Bayes point estimates of these parameters are provided by the method.

**Method.** The method returns estimates of $(s_1^2, \dots, s_L^2)$ and $(\sigma^2, \tau^2)$ for this locus, together with approximations of the following quantities for each single effect $l \in \{1, \dots, L\}$ and each SNP $j \in \{1, \dots, p\}$:

- $\text{PIP}_j^{(l)} = \Pr[\gamma^{(l)} = e_j \mid X, y]$ the posterior probability that $j$ is the causal SNP for effect $l$.
- $b_j^{(l)}|\text{causal} = \mathbb{E}[\mu^{(l)} \mid \gamma^{(l)} = e_j,\, X, y]$ the posterior mean effect size conditional on inclusion.
- $s_j^{(l)}|\text{causal} = \sqrt{\text{Var}[\mu^{(l)} \mid \gamma^{(l)} = e_j,\, X, y]}$ the posterior standard deviation of effect size conditional on inclusion.
- $a_j = \mathbb{E}[\alpha_j \mid X, y]$ the posterior mean infinitesimal effect.

As in SuSiE, SuSiE-inf computes a variational approximation to the posterior distribution of $\beta^{(1)}, \dots, \beta^{(L)}$ in which these single effects are independent. Under this independence approximation, the posterior-inclusion-probability for SNP $j$ aggregated across all $L$ effects is then computed as

$$\text{PIP}_j = \Pr[\beta_j \neq 0 \mid X, y] \approx 1 - \prod_{l=1}^{L} (1 - \text{PIP}_j^{(l)}).$$

The total posterior mean of the sparse effect for SNP $j$ not conditional on inclusion is $b_j = \mathbb{E}[\beta_j \mid X, y] = \sum_{l=1}^{L} \text{PIP}_j^{(l)} \cdot b_j^{(l)}|\text{causal}$, and the total posterior mean genetic effect of SNP $j$ is $b_j + a_j$.

Let us denote the variational approximation to the posterior distribution of $\beta$ marginalized over $(\alpha, \varepsilon)$ as

$$\Pr[\beta^{(1)}, \dots, \beta^{(L)} \mid X, y] \approx \prod_{l=1}^{L} q(\beta^{(l)}). \tag{29}$$

SuSiE-inf alternates between

1. Updating the prior effect size variance $s_l^2$ and the distribution $q(\beta^{(l)})$, sequentially for each effect $l = 1, \dots, L$, and
2. Updating the infinitesimal effect size and noise variances $(\sigma^2, \tau^2)$.

The first step uses a coordinate-ascent variational Bayes approach that is similar to that of SuSiE, which we review below for completeness. The second step is based on either a maximum-ELBO or method-of-moments procedure.

*Updates for $s_l^2$ and $q(\beta^{(l)})$.* Denote the prior density of $\beta^{(l)}$ as $\Pr[\beta^{(l)}]$ and the log-likelihood of $\beta^{(1)}, \dots, \beta^{(L)}$ in the SuSiE model marginalized over $(\alpha, \varepsilon)$ as

$$\log \Pr[y \mid X, \beta^{(1)}, \dots, \beta^{(L)}].$$

Define the variational Bayes evidence lower bound

$$\text{ELBO} = \mathbb{E}_{\beta^{(1)}, \dots, \beta^{(L)} \sim q} \left[ \log \Pr[y \mid X, \beta^{(1)}, \dots, \beta^{(L)}] + \sum_{l=1}^{L} \log \frac{\Pr[\beta^{(l)}]}{q(\beta^{(l)})} \right] \tag{30}$$

where the expectation is over $(\beta^{(1)}, \dots, \beta^{(L)}) \sim q$ described by the posterior approximation of Eq. (29). SuSiE-inf maximizes the ELBO with respect to each pair $(s_l^2, q(\beta^{(l)}))$, sequentially for $l = 1, \dots, L$.

Each such maximization has an explicit form: Define a residual vector

$$r^{(l)} = y - X \sum_{k:k\neq l} \mathbb{E}_{\beta^{(k)}\sim q}[\beta^{(k)}] \tag{31}$$

and consider the single-effect regression (SER) model

$$r^{(l)} = X\beta^{(l)} + X\alpha + \varepsilon \tag{32}$$

where $\beta^{(l)} = \mu^{(l)} \cdot \gamma^{(l)}$ and $(\alpha, \varepsilon)$ are as specified in the SuSiE model. Then the update for $s_l^2$ is given by

$$s_l^2 = \arg\max_{s_l^2} \log \Pr[r^{(l)} \mid X], \tag{33}$$

the maximum-likelihood estimate for $s_l^2$ in the SER model upon marginalizing over the effect size $\mu^{(l)} \sim \mathcal{N}(0, s_l^2)$ of $\beta^{(l)}$. This maximization is carried out numerically using Brent's method over a user-specified range $[s^2_{\min}, s^2_{\max}]$. The update for $q(\beta^{(l)})$ is then given by

$$q(\beta^{(l)}) = \Pr[\beta^{(l)} \mid X, r^{(l)}], \tag{34}$$

the posterior distribution for $\beta^{(l)}$ in the SER model with this estimated prior variance $s_l^2$. This distribution $q(\beta^{(l)})$ remains a distribution over a single effect $\beta^{(l)} = \mu^{(l)} \cdot \gamma^{(l)}$, and it may be parameterized by the probability $q(\gamma^{(l)} = e_j)$ and a normal conditional posterior distribution $q(\mu^{(l)} \mid \gamma^{(l)} = e_j)$ for each SNP $j = 1, \ldots, p$.

*Updates for* $(\sigma^2, \tau^2)$*.* SuSiE-inf implements two alternative approaches for updating $(\sigma^2, \tau^2)$. The first is to set $(\sigma^2, \tau^2)$ to maximize the ELBO in Eq. (30), which depends on $(\sigma^2, \tau^2)$ via the marginal log-likelihood $\log \Pr[y \mid X, \beta^{(1)}, \ldots, \beta^{(L)}]$. This maximization is performed fixing the current estimates of $(s_l^2, q(\beta^{(l)}))$ for all effects $l = 1, \ldots, L$, and is carried out numerically using L-BFGS-B over user-specified ranges $[\sigma^2_{\min}, \sigma^2_{\max}]$ and $[\tau^2_{\min}, \tau^2_{\max}]$. The second is a method-of-moments procedure based again on the moment identities of Eqs. (7–8), which are valid also under the SuSiE-inf model of Eq. (28). The posterior expectations in these moment identities are approximated by expectations under the current estimates of $q(\beta^{(1)}), \ldots, q(\beta^{(L)})$,

$$\mathbb{E}[\|y - X\beta\|_2^2 \mid X, y] \approx m_1 := \mathbb{E}_{\beta^{(1)},\ldots,\beta^{(L)}\sim q}\left[\left\| y - X\sum_{l=1}^{L} \beta^{(l)} \right\|_2^2\right] \tag{35}$$

$$\mathbb{E}[\|X^\top(y - X\beta)\|_2^2 \mid X, y] \approx m_2 := \mathbb{E}_{\beta^{(1)},\ldots,\beta^{(L)}\sim q}\left[\left\| X^\top\left(y - X\sum_{l=1}^{L} \beta^{(l)}\right) \right\|_2^2\right]. \tag{36}$$

The values of $(\sigma^2, \tau^2)$ are then updated by solving Eqs. (5–6) with $(m_1, m_2)$ in place of the left sides of these moment equations, yielding

$$\begin{pmatrix} \sigma^2_{\text{new}} \\ \tau^2_{\text{new}} \end{pmatrix} = \begin{pmatrix} n & \operatorname{Tr} X^\top X \\ \operatorname{Tr} X^\top X & \operatorname{Tr}(X^\top X)^2 \end{pmatrix}^{-1} \begin{pmatrix} m_1 \\ m_2 \end{pmatrix}. \tag{37}$$

For loci where the resulting estimate $\sigma^2_{\text{new}}$ or $\tau^2_{\text{new}}$ is negative, we constrain $\tau^2_{\text{new}} = 0$ and use instead the update $(\sigma^2_{\text{new}}, \tau^2_{\text{new}}) = (n^{-1}m_1, 0)$.

*Computing* $\text{PIP}_j^{(l)}$*,* $b_j^{(l)}|\text{causal}$*,* $s_j^{(l)}|\text{causal}$*, and* $a_j$*.* The PIPs and posterior moments of $\beta_j^{(l)}$ conditional on inclusion are approximated using the final variational approximation given by $q(\beta^{(1)}), \ldots, q(\beta^{(L)})$. Representing $\beta^{(l)} = \mu^{(l)} \cdot \gamma^{(l)}$ under both the true posterior law and its approximation by $q(\beta^{(l)})$,

$$\text{PIP}_j^{(l)} = \Pr[\gamma^{(l)} = e_j \mid X, y] \approx q(\gamma^{(l)} = e_j) \tag{38}$$

$$b_j^{(l)}|\text{causal} = \mathbb{E}[\mu^{(l)} \mid \gamma^{(l)} = e_j,\, X, y] \approx \mathbb{E}_{\beta^{(l)}\sim q}[\mu^{(l)} \mid \gamma^{(l)} = e_j] \tag{39}$$

$$\mathbb{E}[(\mu^{(l)})^2 \mid \gamma^{(l)} = e_j,\, X, y] \approx \mathbb{E}_{\beta^{(l)}\sim q}[(\mu^{(l)})^2 \mid \gamma^{(l)} = e_j]. \tag{40}$$

The standard deviation $s_j^{(l)}|\text{causal}$ is computed from the above approximations as

$$s_j^{(l)}|\text{causal} = \sqrt{\mathbb{E}[(\mu^{(l)})^2 \mid \gamma^{(l)} = e_j,\, X, y] - (b_j^{(l)}|\text{causal})^2}.$$

The posterior mean $a_j = \mathbb{E}[\alpha_j \mid X, y]$ is computed as in Eq. (14), where $\hat{\alpha}(\beta)$ is the BLUP for $\alpha$ given residuals $y - X\beta$, and its posterior mean $a = \mathbb{E}[\hat{\alpha}(\beta) \mid X, y]$ is approximated by $\mathbb{E}_{\beta^{(1)},\ldots,\beta^{(L)}\sim q}[\hat{\alpha}(\beta)]$.

**Parameter settings.** In our analyses, we standardize $y \in \mathbb{R}^n$ and $X \in \mathbb{R}^{n\times p}$ to have columns of mean 0 and variance 1. We initialize $(\sigma^2, \tau^2) = (1, 0)$, $s_l^2 = 0.2$, and $q(\beta^{(l)}) = \Pr[\beta^{(l)}]$ (the prior distribution for $\beta^{(l)} = \mu^{(l)} \cdot \gamma^{(l)}$) for each effect $l = 1, \ldots, L$. We make these choices so as to match the default prior parameters and initializations of SuSiE.

We fix $L = 10$ as the number of causal variants. We have found the method-of-moments approach for updating $(\sigma^2, \tau^2)$ to be more stable and robust than the maximum-ELBO method, and use this in our reported results. (If using the alternative maximum-ELBO method, we recommend setting a non-zero lower bound for $\sigma^2_{\min}$ to that the maximization does not degenerate to a solution where $\sigma^2 = 0$.) We restrict the optimization for $s_l^2$ to the range $[s^2_{\min}, s^2_{\max}] = [0, 1]$. Convergence is assessed by the absolute change in $\text{PIP}_j^{(l)}$ between iterations, and we terminate the algorithm when the maximum such change across all $L$ effects and all $p$ SNPs falls below $\text{tol} = 10^{-3}$.

**Efficient computation.** As in FINEMAP-inf, for computational efficiency, SuSiE-inf pre-computes the eigen-decomposition $X^\top X = VDV^\top$ and rotated and scaled z-scores $u = V^\top X^\top y$. All remaining quantities are computed from $V, D, u$ and the mean-squared phenotype $\langle y^2\rangle = \frac{1}{n} y^\top y$.

We denote $\mathbb{E}_q$ as shorthand for the expectation over $\beta^{(1)}, \ldots, \beta^{(L)} \sim q$. We describe in the remainder of this section the quantitative form of the ELBO in Eq. (30), the details of its maximization over $(s_l^2, q(\beta^{(l)}))$, its maximization over $(\sigma^2, \tau^2)$ in the maximum-ELBO approach, computation of $\mathbb{E}_q[\|y - X\sum_{l=1}^L \beta^{(l)}\|_2^2, \|X^\top(y - X\sum_{l=1}^L \beta^{(l)})\|_2^2]$ in the method-of-moments approach, and computation of $\mathbb{E}_q[\hat{\alpha}(\beta)]$ to obtain the posterior mean of $\alpha$. We conclude with a discussion of the total computational complexity.

*Form of the ELBO.* Analogously to Eq. (17), the SuSiE-inf model is equivalently written as

$$y = X\sum_{l=1}^L \beta^{(l)} + \tilde{\varepsilon} \in \mathbb{R}^n, \qquad \tilde{\varepsilon} \sim \mathcal{N}(0, \Omega^{-1}), \qquad \Omega = (\tau^2 XX^\top + \sigma^2 I)^{-1}$$

where $\Omega$ is the precision matrix of the effective residual $\tilde{\varepsilon} = X\alpha + \varepsilon$. Thus $y$ has the normal log-likelihood

$$\log \Pr[y \mid X, \beta^{(1)}, \ldots, \beta^{(L)}] = \log\left(\frac{\det \Omega^{1/2}}{(2\pi)^{n/2}} \exp\left[-\frac{1}{2}\left(y - X\sum_{l=1}^L \beta^{(l)}\right)^\top \Omega \left(y - X\sum_{l=1}^L \beta^{(l)}\right)\right]\right),$$

and the ELBO in Eq. (30) takes the form

$$\text{ELBO} = -\frac{n}{2}\log 2\pi + \frac{1}{2}\log\det\Omega + \mathbb{E}_q\left[-\frac{1}{2}\left(y - X\sum_{l=1}^L \beta^{(l)}\right)^\top \Omega\left(y - X\sum_{l=1}^L \beta^{(l)}\right) + \sum_{l=1}^L \log\frac{\Pr[\beta^{(l)}]}{q(\beta^{(l)})}\right]. \quad (41)$$

*Maximization over $s_l^2$ and $q(\beta^{(l)})$.* Isolating the terms depending on $\beta^{(l)}$, computing first the expectation over $\beta^{(k)} \sim q(\beta^{(k)})$ for all $k \neq l$, and introducing the residual $r^{(l)}$ defined by Eq. (31), the ELBO is equal to

$$\text{ELBO} = \mathbb{E}_{\beta^{(l)}\sim q}\left[-\frac{1}{2}(r^{(l)} - X\beta^{(l)})^\top \Omega (r^{(l)} - X\beta^{(l)}) + \log\frac{\Pr[\beta^{(l)}]}{q(\beta^{(l)})}\right] + \text{const}$$

where const does not depend on $(s_l^2, q(\beta^{(l)}))$. Up to an additive constant independent of $(s_l^2, q(\beta^{(l)}))$, the first term inside $\mathbb{E}_{\beta^{(l)}\sim q}$ is the log-likelihood of $\beta^{(l)}$ in the single-effect regression (SER) model

$$r^{(l)} = X\beta^{(l)} + \tilde{\varepsilon}, \qquad \tilde{\varepsilon} \sim \mathcal{N}(0, \Omega^{-1}),$$

which is equivalent to the previously stated model of Eq. (32). Then, up to an additive constant, the above expression is the evidence lower bound in this SER model with prior $\Pr[\beta^{(l)}]$. Its maximum over distributions $q(\beta^{(l)})$ is attained at the posterior distribution of Eq. (34), and its maximum value is given by the log-marginal density of $r^{(l)}$,

$$\max_{q(\beta^{(l)})} \text{ELBO} = \log \Pr[r^{(l)} \mid X] + \text{const.}$$

Hence the maximum over $s_l^2$ is attained at the maximum-likelihood estimate of Eq. (33). This verifies the previously stated forms of Eqs. (33) and (34) for these updates.

To derive explicit forms for these updates, let $x_j$ be the $j^{\text{th}}$ column of $X$, and denote

$$z_j^{(l)} = x_j^\top \Omega r^{(l)}, \qquad \omega_j(s_l^2) = x_j^\top \Omega x_j + s_l^{-2}. \tag{42}$$

Writing $\beta^{(l)} = \mu^{(l)} \cdot \gamma^{(l)}$, the SuSiE-inf prior specifies $\gamma^{(l)} \sim \text{Uniform}(e_1, \ldots, e_p)$. Conditioning on $\gamma^{(l)} = e_j$ and marginalizing over $\mu^{(l)} \sim \mathcal{N}(0, s_l^2)$, we have $r^{(l)} = \mu^{(l)} x_j + \tilde{\varepsilon} \sim \mathcal{N}(0, s_l^2 x_j x_j^\top + \Omega^{-1})$. Thus the marginal density of $r^{(l)}$ in the SER model is

$$\Pr[r^{(l)} \mid X] = \frac{1}{p} \sum_{j=1}^{p} (2\pi)^{-n/2} \det\left(s_l^2 x_j x_j^\top + \Omega^{-1}\right)^{-1/2} \cdot \exp\left(-\frac{1}{2}(r^{(l)})^\top \left(s_l^2 x_j x_j^\top + \Omega^{-1}\right)^{-1} r^{(l)}\right)$$

By the Sherman-Morrison identity and rank-one update rule for matrix determinant,

$$\left(s_l^2 x_j x_j^\top + \Omega^{-1}\right)^{-1} = \Omega - \frac{s_l^2 \Omega x_j x_j^\top \Omega}{1 + s_l^2 x_j^\top \Omega x_j} = \Omega - \frac{\Omega x_j x_j^\top \Omega}{\omega_j(s_l^2)}$$
$$\det(s_l^2 x_j x_j^\top + \Omega^{-1}) = (1 + s_l^2 x_j^\top \Omega x_j) \cdot \det(\Omega^{-1}) = s_l^2 \omega_j(s_l^2) \cdot \det(\Omega^{-1})$$

Substituting these forms, we obtain

$$\log \Pr[r^{(l)} \mid X] = \log \sum_{j=1}^{p} (s_l^2 \omega_j(s_l^2))^{-1/2} \cdot \exp\left((z_j^{(l)})^2 / 2\omega_j(s_l^2)\right) + \text{const} \tag{43}$$

for a constant independent of $s_l^2$. This is maximized over $s_l^2 \in [s_{\min}^2, s_{\max}^2]$ to obtain the update of $s_l^2$.

For the posterior distribution of $\beta^{(l)}$, absorbing constants not depending on $j$ into the notation $\propto$,

$$\begin{aligned}
q(\gamma^{(l)} = e_j) &= \Pr[\gamma^{(l)} = e_j \mid X, r^{(l)}] \\
&\propto \Pr[r^{(l)} \mid X, \gamma^{(l)} = e_j] \cdot \Pr[\gamma^{(l)} = e_j] \\
&\propto \det\left(s_l^2 x_j x_j^\top + \Omega^{-1}\right)^{-1/2} \cdot \exp\left(-\frac{1}{2}(r^{(l)})^\top \left(s_l^2 x_j x_j^\top + \Omega^{-1}\right)^{-1} r^{(l)}\right) \\
&\propto \omega_j(s_l^2)^{-1/2} \cdot \exp\left((z_j^{(l)})^2 / 2\omega_j(s_l^2)\right).
\end{aligned}$$

Thus

$$q(\gamma^{(l)} = e_j) = \frac{\omega_j(s_l^2)^{-1/2} \cdot \exp\left((z_j^{(l)})^2 / 2\omega_j(s_l^2)\right)}{\sum_{i=1}^{p} \omega_i(s_l^2)^{-1/2} \cdot \exp\left((z_i^{(l)})^2 / 2\omega_i(s_l^2)\right)}. \tag{44}$$

Conditional on $\gamma^{(l)} = e_j$, the posterior distribution of $\mu^{(l)}$ is

$$\begin{aligned}
q(\mu^{(l)} \mid \gamma^{(l)} = e_j) &= \Pr[\mu^{(l)} \mid \gamma^{(l)} = e_j, X, r^{(l)}] \\
&\propto \exp\left(-\frac{1}{2}(r^{(l)} - \mu^{(l)} x_j)^\top \Omega (r^{(l)} - \mu^{(l)} x_j)\right) \cdot \exp\left(-\frac{1}{2s_l^2}(\mu^{(l)})^2\right) \\
&\propto \exp\left(-(\mu^{(l)})^2 \cdot \omega_j(s_l^2)/2 + \mu^{(l)} \cdot z_j^{(l)}\right).
\end{aligned}$$

This is the normal distribution

$$q(\mu^{(l)} \mid \gamma^{(l)} = e_j) = \mathcal{N}\Big(\omega_j(s_l^2)^{-1} z_j^{(l)},\, \omega_j(s_l^2)^{-1}\Big). \tag{45}$$

Eqs. (44) and (45) together define the update for $q(\beta^{(l)})$.

To express Eqs. (43), (44), and (45) in terms of $V, D, u$ defined in Eqs. (15) and (16), observe that as a special case of Eq. (20) with $\Gamma = \{j\}$, we have

$$x_j^\top \Omega x_j = v_j^\top D(\tau^2 D + \sigma^2 I)^{-1} v_j \tag{46}$$

where $v_j$ is the $j^{\text{th}}$ row of $V$. Adding to this $s_l^{-2}$ gives $\omega_j(s_l^2)$. For $z_j^{(l)}$, observe that multiplying Eq. (21) on the right by $y - X\sum_{k:k\neq l} \mathbb{E}_q[\beta^{(k)}]$ and taking row $j$ gives

$$z_j^{(l)} = x_j^\top \Omega r^{(l)} = v_j^\top (\tau^2 D + \sigma^2 I)^{-1} \left( u - DV^\top \sum_{k:k\neq l} \mathbb{E}_q[\beta^{(k)}] \right). \tag{47}$$

These expressions may then be used to compute the quantities of Eqs. (43), (44), and (45).

*Maximization over* $(\sigma^2, \tau^2)$. Denote

$$b = \sum_{l=1}^{L} \mathbb{E}_q[\beta^{(l)}], \qquad M = bb^\top + \sum_{l=1}^{L} \mathbb{E}_q[\beta^{(l)}\beta^{(l)\top}] - \mathbb{E}_q[\beta^{(l)}]\mathbb{E}_q[\beta^{(l)}]^\top. \tag{48}$$

This matrix $M$ is defined such that, for any symmetric matrix $K \in \mathbb{R}^{p\times p}$,

$$\mathbb{E}_q\left[ \left( \sum_{l=1}^{L} \beta^{(l)} \right)^\top K \left( \sum_{l=1}^{L} \beta^{(l)} \right) \right] = \operatorname{Tr}\left( K \cdot \sum_{l=1}^{L}\sum_{m=1}^{L} \mathbb{E}_q[\beta^{(l)}\beta^{(m)\top}] \right) = \operatorname{Tr}(K \cdot M) \tag{49}$$

where the second equality uses independence of $\beta^{(1)}, \ldots, \beta^{(L)}$ under $q$.

Let us express the terms of the ELBO in Eq. (41) that depend on $\Omega = (\tau^2 XX^\top + \sigma^2 I)^{-1}$ in terms of $V, D, u$: The eigenvalues of $XX^\top$ coincide with those of $X^\top X$ up to the addition or removal of $|n-p|$ zeros. Thus, the eigenvalues of $\tau^2 XX^\top + \sigma^2 I$ coincide with those of $\tau^2 X^\top X + \sigma^2 I = V(\tau^2 D + \sigma^2 I)V^\top$ up to the addition or removal of $|n-p|$ eigenvalues equal to $\sigma^2$. This implies

$$\log\det \Omega^{-1} = \log\det(\tau^2 XX^\top + \sigma^2 I) = (n-p)\log\sigma^2 + \log\det(\tau^2 D + \sigma^2 I).$$

Applying this, the definitions of $b$ and $M$ in Eq. (48), and the identity of Eq. (49), the ELBO takes the form

$$\text{ELBO} = -\frac{n-p}{2}\log\sigma^2 - \frac{1}{2}\log\det(\tau^2 D + \sigma^2 I) - \frac{1}{2} y^\top \Omega y + b^\top X^\top \Omega y - \frac{1}{2}\operatorname{Tr}\left(X^\top \Omega X \cdot M\right) + \text{const}$$

for a constant independent of $(\sigma^2, \tau^2)$. From the identities of Eq. (19) and (21), we have

$$\operatorname{Tr}\left(X^\top \Omega X \cdot M\right) = \operatorname{Tr}\left(D(\tau^2 D + \sigma^2 I)^{-1} \cdot V^\top M V\right),$$
$$b^\top X^\top \Omega y = (V^\top b)^\top (\tau^2 D + \sigma^2 I)^{-1} u.$$

We may similarly derive from the Woodbury identity

$$\begin{aligned} y^\top \Omega y &= \sigma^{-2} y^\top \Big((\tau^2/\sigma^2) XX^\top + I\Big)^{-1} y \\ &= \sigma^{-2} y^\top \left( I - X\Big((\sigma^2/\tau^2) I + X^\top X\Big)^{-1} X^\top \right) y \\ &= \sigma^{-2} y^\top \left( I - XV\Big((\sigma^2/\tau^2) I + D\Big)^{-1} V^\top X^\top \right) y = \frac{n\langle y^2\rangle}{\sigma^2} - \frac{\tau^2}{\sigma^2} u^\top (\tau^2 D + \sigma^2 I)^{-1} u. \end{aligned}$$

Then

$$\text{ELBO} = -\frac{n-p}{2}\log\sigma^2 - \frac{1}{2}\log\det(\tau^2 D + \sigma^2 I) - \frac{n\langle y^2\rangle}{2\sigma^2} + \frac{\tau^2}{2\sigma^2}u^\top(\tau^2 D + \sigma^2 I)^{-1}u$$
$$+ (V^\top b)^\top(\tau^2 D + \sigma^2 I)^{-1}u - \frac{1}{2}\operatorname{Tr}\left(D(\tau^2 D + \sigma^2 I)^{-1}\cdot V^\top M V\right) + \text{const.} \tag{50}$$

This is maximized over $\sigma^2 \in [\sigma^2_{\min}, \sigma^2_{\max}]$ and $\tau^2 \in [\tau^2_{\min}, \tau^2_{\max}]$ to obtain the update for $(\sigma^2, \tau^2)$ in the maximum ELBO approach.

*Method-of-moments for* $(\sigma^2, \tau^2)$. We expand the squares on the right sides of Eqs. (35–36) and apply again the definitions of $b$ and $M$ in Eq. (48) and the identity of Eq. (49) to obtain

$$\begin{aligned}\mathbb{E}_q\left[\left\|y - X\sum_{l=1}^L \beta^{(l)}\right\|_2^2\right] &= \|y\|^2 - 2b^\top(X^\top y) + \operatorname{Tr}(X^\top X\cdot M)\\ &= n\langle y^2\rangle - 2(V^\top b)^\top u + \operatorname{Tr}(D\cdot V^\top M V) \qquad (51)\\ \mathbb{E}_q\left[\left\|X^\top\left(y - X\sum_{l=1}^L \beta^{(l)}\right)\right\|_2^2\right] &= \|X^\top y\|^2 - 2b^\top(X^\top X)(X^\top y) + \operatorname{Tr}((X^\top X)^2\cdot M)\\ &= \|u\|^2 - 2(V^\top b)^\top D u + \operatorname{Tr}(D^2\cdot V^\top M V). \qquad (52)\end{aligned}$$

These give the forms of $(m_1, m_2)$ in the method-of-moments update in Eq. (37), and $(\sigma^2_{\text{new}}, \tau^2_{\text{new}})$ may be computed from these forms and $\operatorname{Tr} X^\top X = \operatorname{Tr} D$ and $\operatorname{Tr}(X^\top X)^2 = \operatorname{Tr} D^2$ in the alternative method-of-moments approach.

*BLUP for* $\alpha$. The form of $\hat{\alpha}(\beta) = \mathbb{E}[\alpha \mid \beta, X, y]$ in the SuSiE-inf model of Eq. (28) is the same as in the FINEMAP-inf model. Then, marginalizing over $\beta$, we have analogously to Eq. (27)

$$a = \mathbb{E}[\hat{\alpha}(\beta) \mid X, y] = \tau^2 V(\tau^2 D + \sigma^2 I)^{-1}[u - DV^\top \mathbb{E}[\beta \mid X, y]].$$

Here, $\mathbb{E}[\beta \mid X, y] = \sum_{l=1}^L (\text{PIP}_j^{(l)} \cdot b_j^{(l)}|\text{causal})_{j=1}^p$, and this may be approximated using the approximations of Eqs. (38–39) under $q$.

*Computational cost.* SuSiE-inf maintains across iterations the values of

$$q(\gamma^{(l)} = e_j), \qquad \mathbb{E}_q[\mu^{(l)} \mid \gamma^{(l)} = e_j], \qquad \mathbb{E}_q[(\mu^{(l)})^2 \mid \gamma^{(l)} = e_j] \tag{53}$$

for each effect $l = 1, \ldots, L$ and SNP $j = 1, \ldots, p$, which represent the distributions $q(\beta^{(l)})$. These values may be used to compute the moments of $\beta_j^{(l)}$ not conditional on inclusion,

$$\mathbb{E}_q[\beta_j^{(l)}] = q(\gamma^{(l)} = e_j)\cdot \mathbb{E}_q[\mu^{(l)} \mid \gamma^{(l)} = e_j], \tag{54}$$
$$\mathbb{E}_q[(\beta_j^{(l)})^2] = q(\gamma^{(l)} = e_j)\cdot \mathbb{E}_q[(\mu^{(l)})^2 \mid \gamma^{(l)} = e_j]. \tag{55}$$

In addition, for $b$ and $M$ as defined in Eq. (48), SuSiE-inf maintains the value of $V^\top b$ and the diagonal values only of the matrix

$$V^\top M V = V^\top\left(bb^\top + \sum_{l=1}^L \mathbb{E}_q[\beta^{(l)}\beta^{(l)\top}] - \mathbb{E}_q[\beta^{(l)}]\mathbb{E}_q[\beta^{(l)}]^\top\right)V. \tag{56}$$

In each iteration of SuSiE-inf, $(s_l^2, q(\beta^{(l)}))$ for a single effect $l$ may be updated in time $O(p^2)$: The vector

$$u - DV^\top \sum_{k:k\neq l}\mathbb{E}_q[\beta^{(k)}]$$

appearing in Eq. (47) may be computed from $V^\top b$ and Eq. (54) in time $O(p^2)$. Then $x_j^\top \Omega x_j$ in Eq. (46) and $z_j^{(l)}$ in Eq. (47) may be computed for every $j = 1, \ldots, p$ also in time $O(p^2)$. Given these values, each evaluation of Eq. (43) for a single $s_l^2$ requires time $O(p)$. Assuming that the number of evaluations needed to optimize Eq. (43) is $O(1)$, the update of $s_l^2$ hence requires time $O(p)$. Computing Eqs. (44) and (45) across all $j = 1, \ldots, p$ then also requires time $O(p)$, and these update the quantities of Eq. (53) for effect $l$. Then $V^\top b$ and the diagonal entries of $V^\top bb^\top V$ and $V^\top \mathbb{E}_q[\beta^{(l)}]\mathbb{E}_q[\beta^{(l)}]^\top V$ appearing in $V^\top MV$ may be updated in time $O(p^2)$. For the remaining term $V^\top \mathbb{E}_q[\beta^{(l)}\beta^{(l)\top}]V$ of $V^\top MV$, observe that $\mathbb{E}_q[\beta^{(l)}\beta^{(l)\top}]$ is in fact a diagonal matrix, with diagonal entries given by Eq. (55), because $\beta^{(l)}$ has only a single non-zero coordinate under its distribution $q(\beta^{(l)})$. Then $\mathbb{E}_q[\beta^{(l)}\beta^{(l)\top}]V$ may be updated in time $O(p^2)$, and the diagonal entries of $V^\top \mathbb{E}_q[\beta^{(l)}\beta^{(l)\top}]V$ may then also be updated in time $O(p^2)$. This verifies that $(s_l^2, q(\beta^{(l)}))$ is updated in time $O(p^2)$.

Given $V^\top b$ and the diagonal values of $V^\top MV$, each evaluation of Eq. (50) and its derivatives in $(\sigma^2, \tau^2)$ requires time $O(p)$. Assuming that the number of such evaluations needed to optimize Eq. (50) is $O(1)$, the maximum-ELBO update of $(\sigma^2, \tau^2)$ requires time $O(p)$. Alternatively, given $V^\top b$ and the diagonal values of $V^\top MV$, the quantities of Eqs. (51–52) may also be evaluated in time $O(p)$, and hence the method-of-moments update of $(\sigma^2, \tau^2)$ via Eq. (37) also requires time $O(p)$.

Thus, each iteration of SuSiE-inf may be performed in time $O(Lp^2)$. Letting $T$ be the total number of iterations, and assuming that the initial eigen-decomposition of LD requires time $O(p^3)$, the total computational cost of SuSiE-inf is

$$O(p^3 + TLp^2).$$

Note that naively recomputing $x_i^\top \Omega x_j$ and $x_i^\top \Omega y$ after each $(\sigma^2, \tau^2)$ update by direct matrix inversion, without precomputing the eigen-decomposition of LD, would instead require time $O(p^3)$ per iteration, yielding a total computational cost of $O(Tp^3)$.

### F Variants in Low Complexity Regions (LCR)

We recommend filtering out variants in LCR prior to fine-mapping. Ideally, variants in low-complexity regions (LCR) would be filtered out before imputation due to higher chance of genotyping error and therefore worse imputation performance. However, we found that these variants are in the imputed genotypes provided by UK Biobank, therefore they are included in our fine-mapping pipeline. Around 5% of total variants included in our GWAS are in LCR, and around 6% of total fine-mapped variants are in LCR. We provide a list of variants that are in LCR and obtained nontrivial PIP ($\geq$0.1) from any of the six fine-mapping methods (SuSiE, FINEMAP, COJO-ABF, ABF, SuSiE-inf, and FINEMAP-inf) in **Supplementary Table 28**. Since the function of variants in LCR is mostly unknown and the accuracy of genotyping in these regions is also unknown, we recommend caution when interpreting results at or near these variants. **Supplementary Fig. 6-7** show different fine-mapping results with and without LCR variants at the APOE locus for LDLC. This does not affect the overall message of our manuscript since all results involving computing RFR and PRS are based on comparison between methods where the same sets of variants are included.

### G Figures

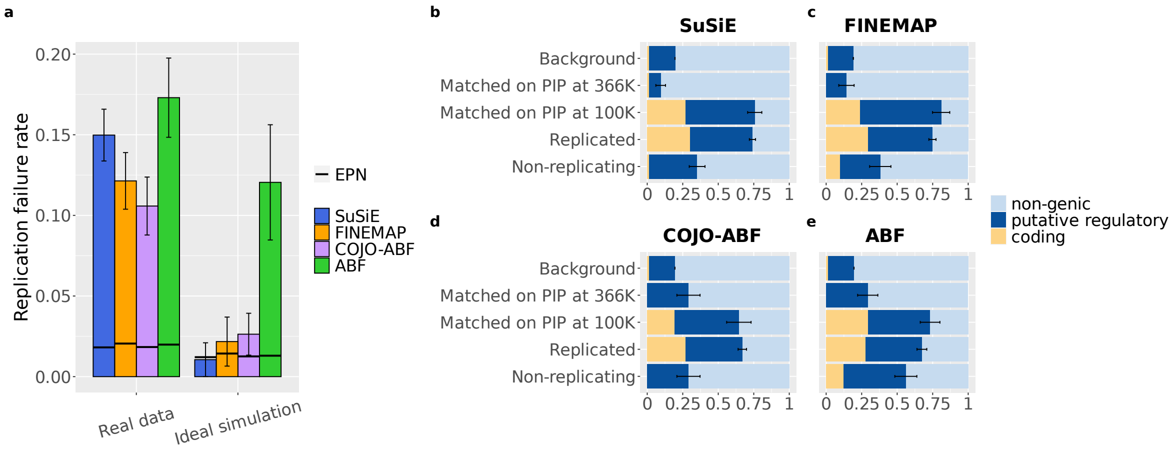

**Supplementary Fig. 1 | RFR and functional enrichment of SuSiE, FINEMAP, ABF and COJO-ABF. a.** RFR in real data (aggregated across 10 UKBB phenotypes) and in ideal simulations. **b.** Functional enrichment for 5 groups of variants. See **Supplementary Methods** for the definitions of these groups. Numerical results available in **Supplementary Table 4-5**.

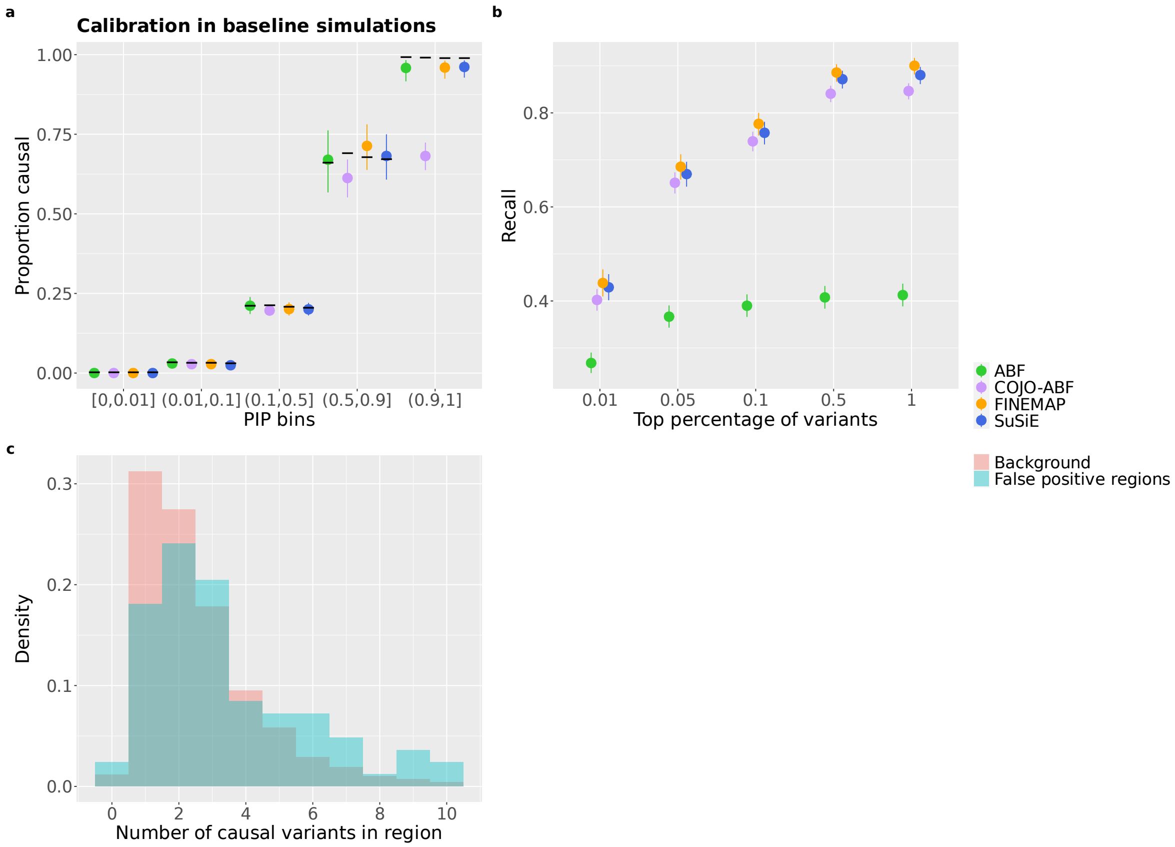

**Supplementary Fig. 2 | ABF and COJO-ABF in ideal simulations. a.** Calibration of ABF and COJO-ABF in ideal simulations. **b.** Recall of ABF and COJO-ABF for the top 0.01%, 0.05%, 0.1%, 0.5% and 1% SNPs, ordered by PIP. **c.** Distribution of the number of causal variants per region in regions containing COJO-ABF false positive SNPs compared to all regions. Numerical results available in **Supplementary Table 29-31**.

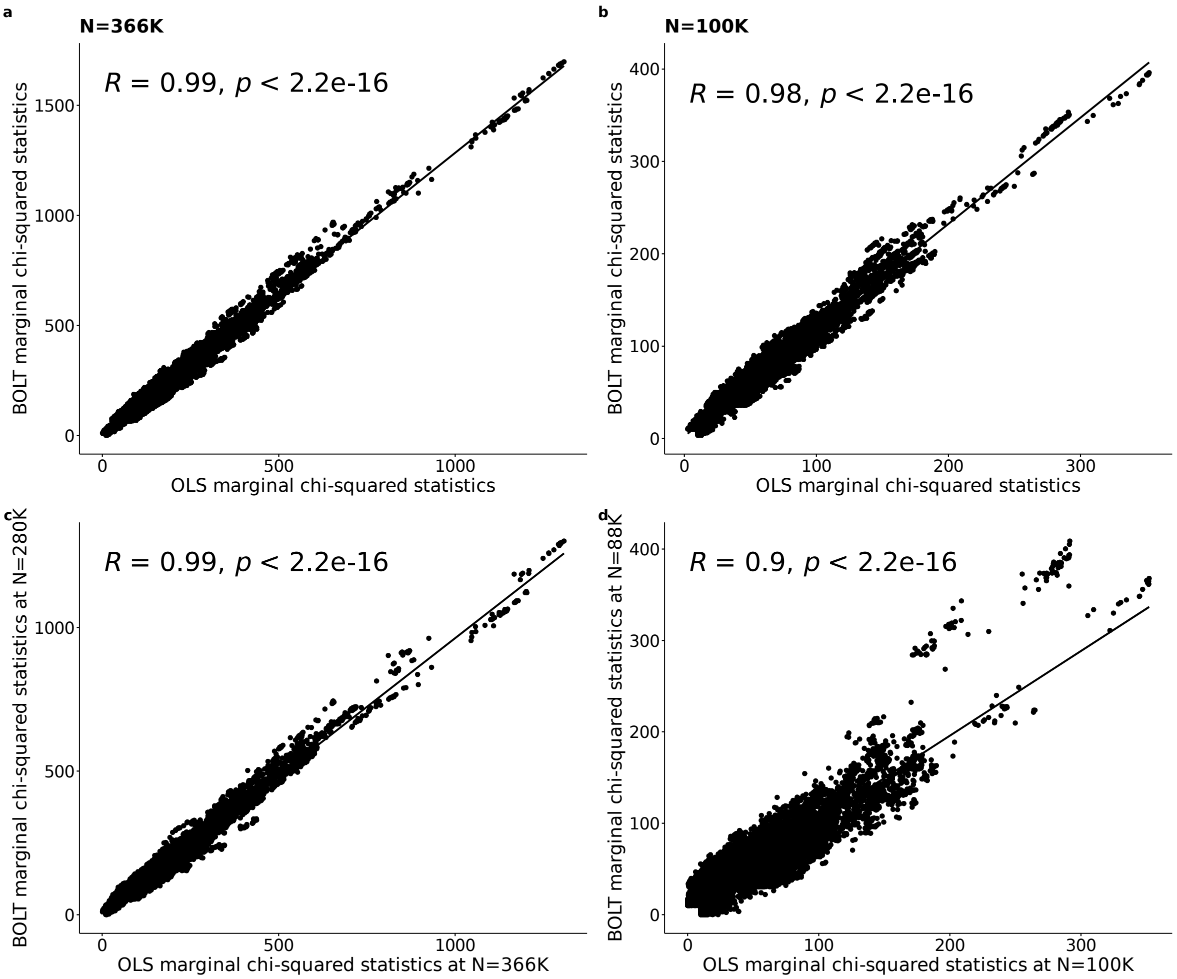

**Supplementary Fig. 3 | Marginal chi-squared statistics comparison between BOLT-LMM and OLS at two different sample sizes. a-b.** Marginal chi-squared statistics at N=366K and N=100K using OLS and BOLT-LMM. **c.** Marginal chi-squared statistics using OLS at N=366K and using BOLT at N=280K. **d.** Marginal chi-squared statistics using OLS at N=100K and using BOLT at N=88K. Not all data points are represented. We plotted Chi-squared statistics greater than 10 for either method, all variants with chi-squared in the top 10 percentile for either method, and sampled a random 3000 variants from the rest.

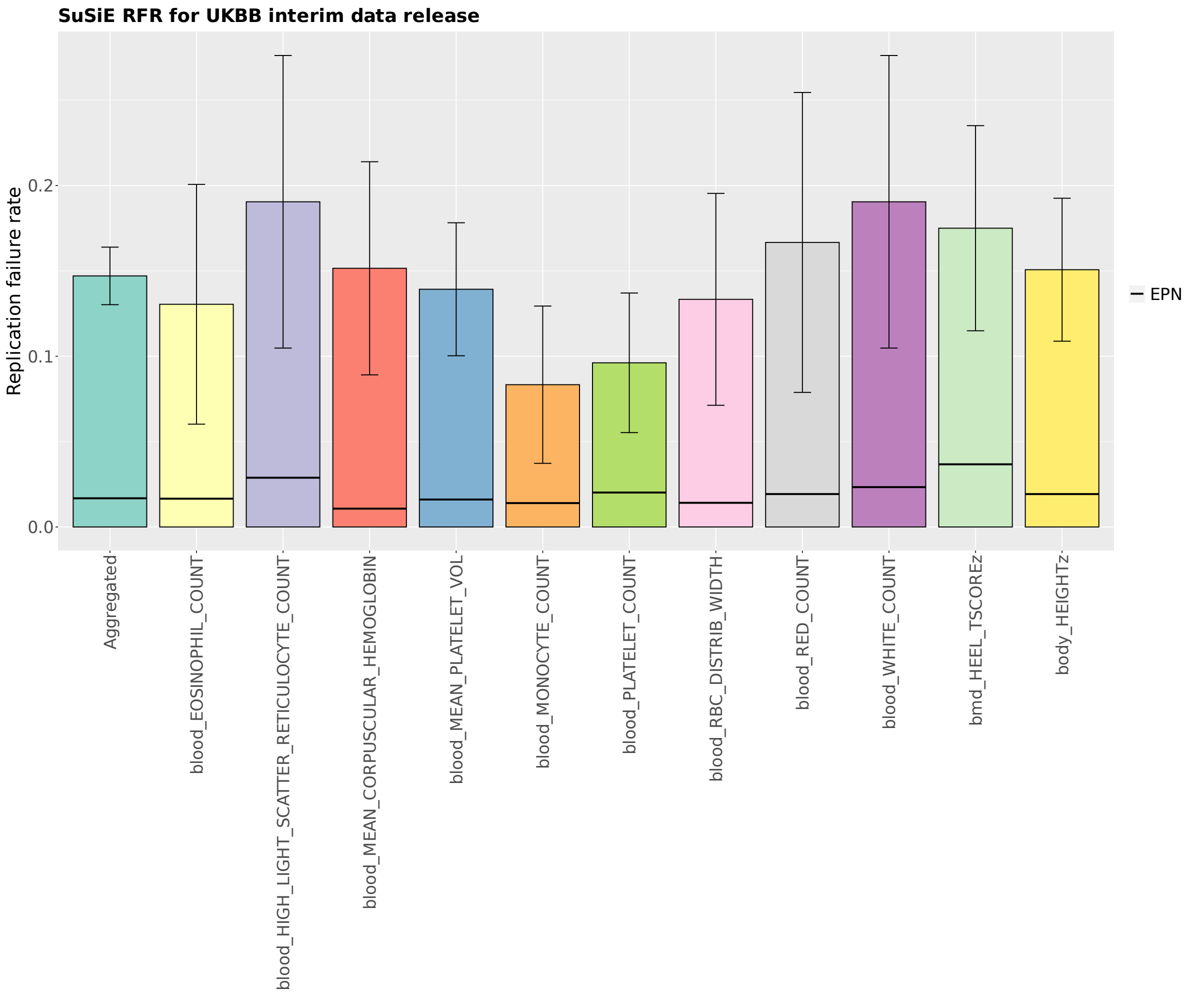

**Supplementary Fig. 4 | SuSiE RFR for UK Biobank interim data release.** Replication failure rates for 11 UK Biobank traits and their aggregated RFR are shown on the plot. The data used to compute these RFRs were obtained from previously published work[7], where Weissbrod et al. performed fine-mapping using SuSiE at sample size N=337K (see [7] for QC and phenotype definitions) and N=107K (interim release). Numerical results available in **Supplementary Table 32**.

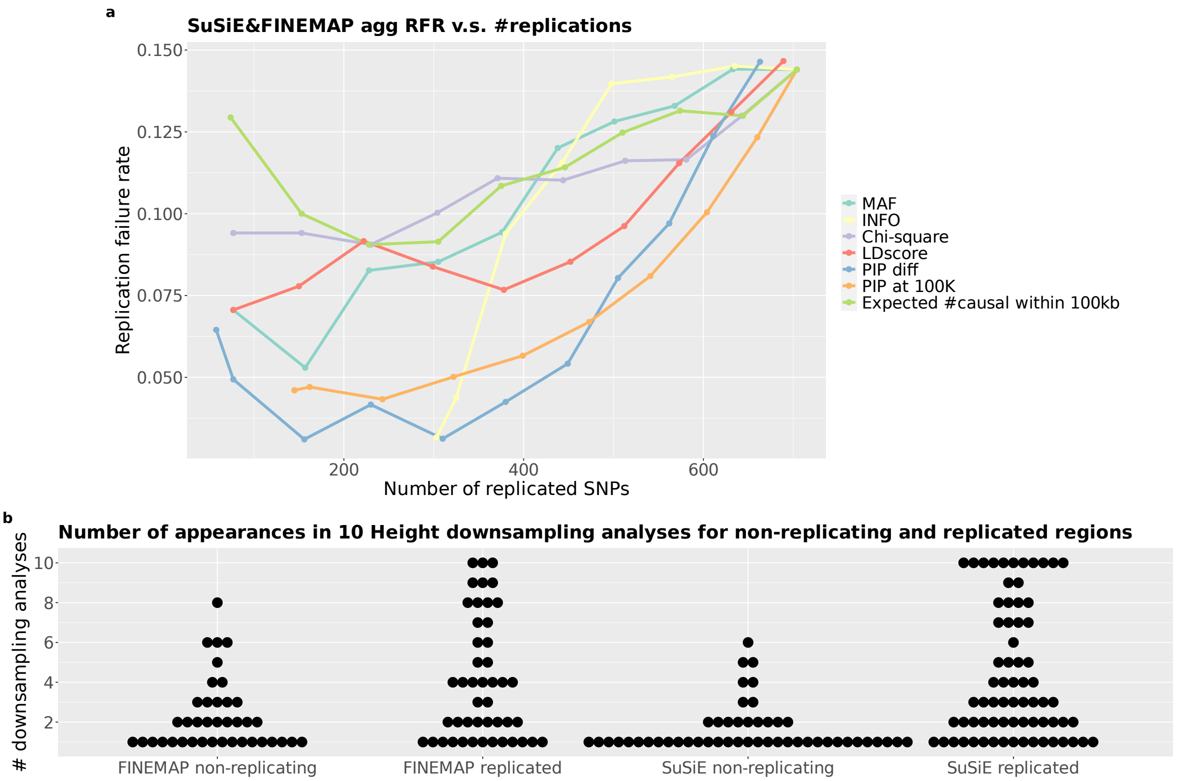

**Supplementary Fig. 5 | Further investigations into non-replication. a.** Number of replicated SNPs and RFR are computed for 10 thresholding values of 7 SNP properties (properties whose values were used as the lower threshold: INFO score, Marginal association chi-squared statistic, PIP at 100K, and expected number of causal variants within 100Kb; upper threshold: MAF, LD score, SuSiE FINEMAP PIP difference. Whether to use value as lower or upper threshold was determined by which setting gives better performance). **b.** Dot plot for the number of occurrences in 10 Height downsampling analyses for replicated/non-replicating regions (region definitions taken at N=366K), each dot represents one region. Numerical results available in **Supplementary Table 33-34**.

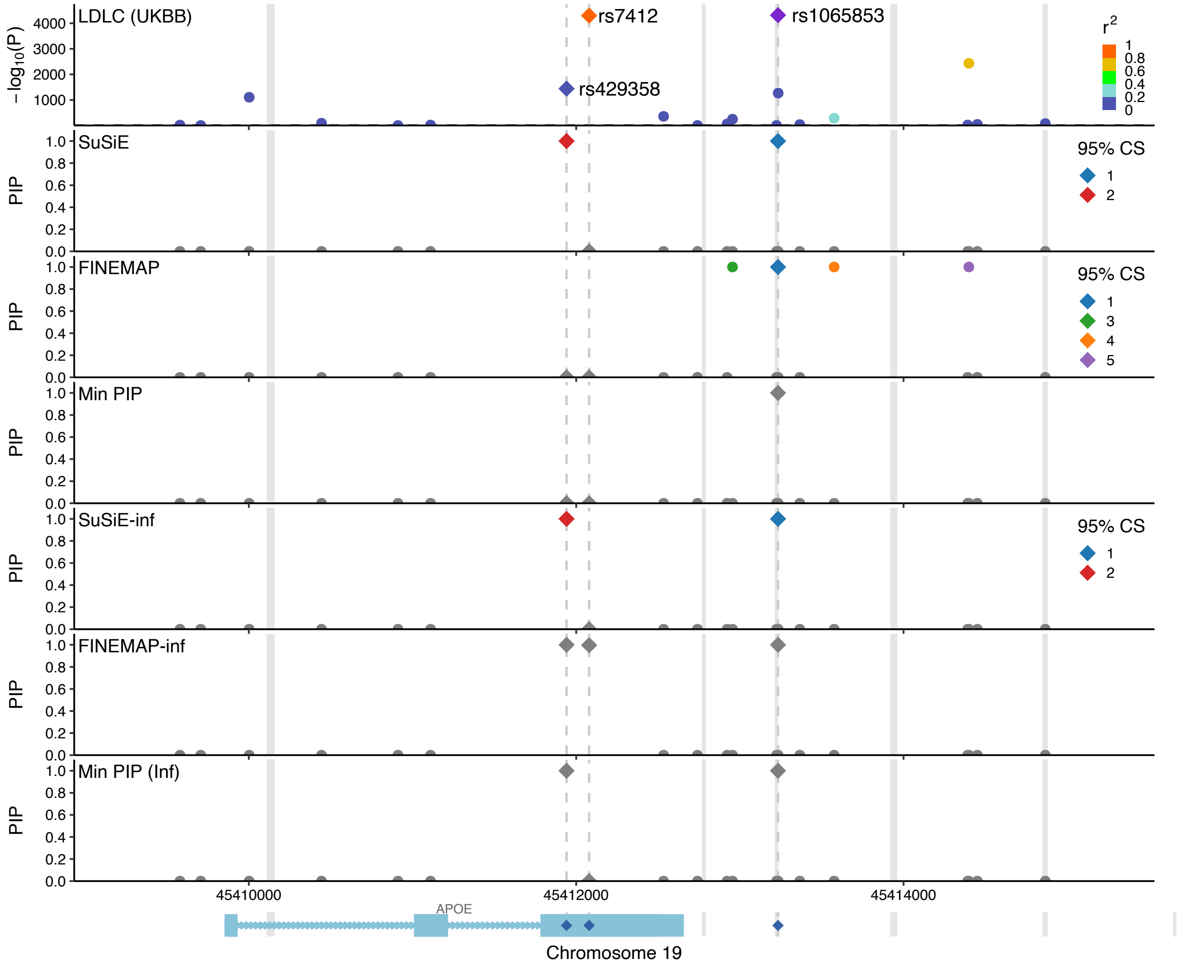

**Supplementary Fig. 12 | APOE locus including variants in LCR**. 6kbp window at the APOE gene location is shown on the plot. GWAS -log10 P-values for trait LDLC are plotted on the top panel, PIPs from 4 fine-mapping methods and 2 aggregating methods are plotted on the subsequent panels. Gray areas denote low-complexity regions (LCR). Variant rs1065853 is in LCR and with high LD to the known causal missense variant rs7412. With LCR included, only FINEMAP-inf was able to identify rs7412 as a high confidence variant. The second known causal variant rs429358 is identified by SuSiE, SuSiE-inf and FINEMAP-inf but not FINEMAP. Taking min PIP between SuSiE and FINEMAP did not capture any of the two known causal variants, whereas minPIP-inf captured one of the two.

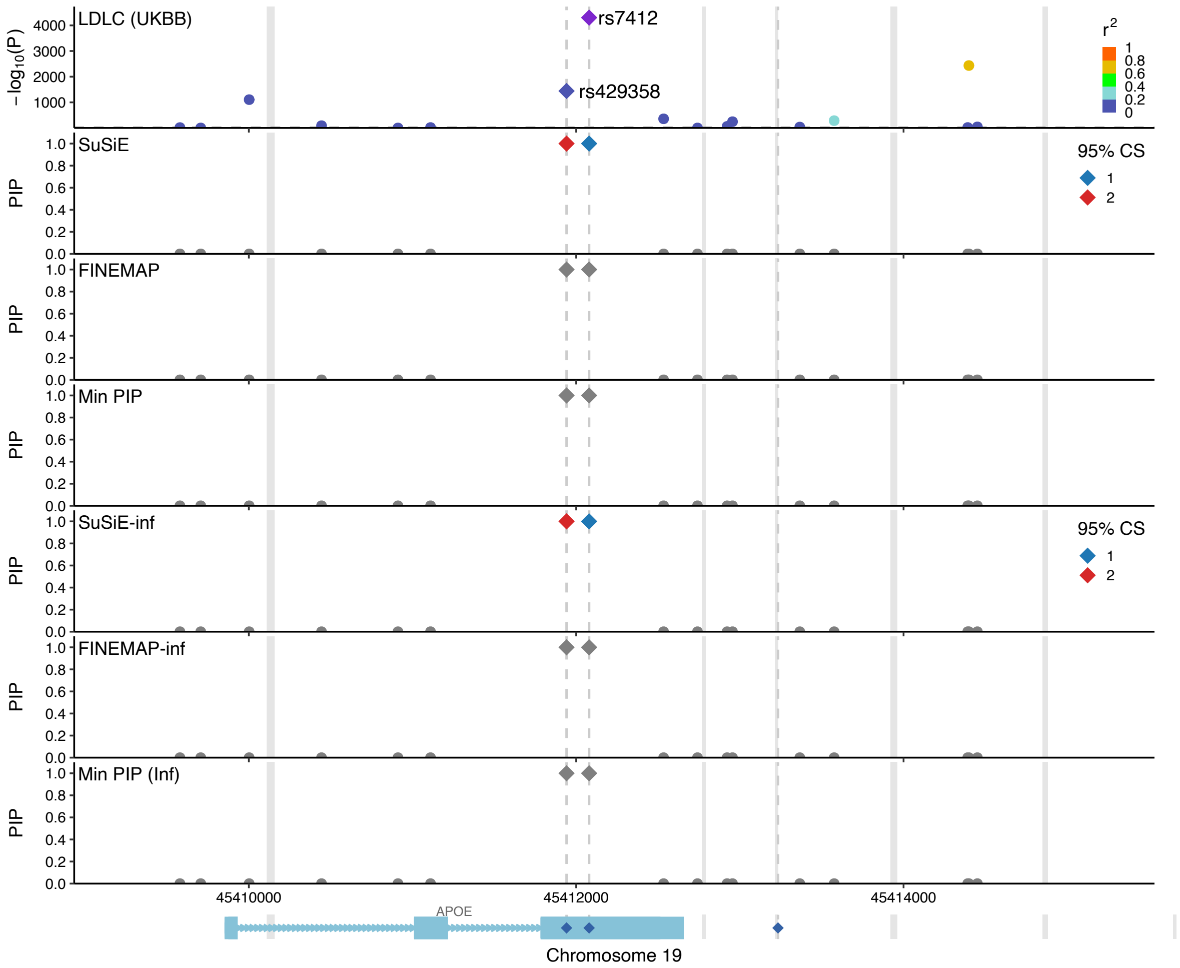

**Supplementary Fig. 13 | APOE locus excluding variants in LCR**. 6kbp window at the APOE gene location is shown on the plot. GWAS -log10 P-values for trait LDLC are plotted on the top panel, PIPs from 4 fine-mapping methods and 2 aggregating methods are plotted on the subsequent panels. Gray areas denote low-complexity regions (LCR). When fine-mapping without variants in LCR, all four methods correctly identified rs7412 and rs429358 as causal variants.

### References

[1] Wakefield, J. (2009). Bayes factors for genome-wide association studies: comparison with P-values. *Genetic Epidemiology: The Official Publication of the International Genetic Epidemiology Society*, 33(1), 79-86.

[2] Yang, J., Ferreira, T., Morris, A. P., Medland, S. E., Genetic Investigation of ANthropometric Traits (GIANT) Consortium, DIAbetes Genetics Replication And Meta-analysis (DIAGRAM) Consortium, ... & Visscher, P. M. (2012). Conditional and joint multiple-SNP analysis of GWAS summary statistics identifies additional variants influencing complex traits. *Nature genetics*, 44(4), 369-375.

[3] Ulirsch JC, Kanai M. An annotated atlas of causal variants for complex human traits. *In preparation.*

[4] Kanai, M., Ulirsch, J. C., Karjalainen, J., Kurki, M., Karczewski, K. J., Fauman, E., ... & Finucane, H. K. (2021). Insights from complex trait fine-mapping across diverse populations. *medRxiv*, 2021-09.

[5] Newcombe, P. J., Conti, D. V., & Richardson, S. (2016). JAM: a scalable Bayesian framework for joint analysis of marginal SNP effects. *Genetic epidemiology*, 40(3), 188-201.

[6] Benner, C., Spencer, C. C., Havulinna, A. S., Salomaa, V., Ripatti, S., & Pirinen, M. (2016). FINEMAP: efficient variable selection using summary data from genome-wide association studies. *Bioinformatics*, 32(10), 1493-1501.

[7] Weissbrod, O., Hormozdiari, F., Benner, C., Cui, R., Ulirsch, J., Gazal, S., ... & Price, A. L. (2020). Functionally informed fine-mapping and polygenic localization of complex trait heritability. *Nature genetics*, 52(12), 1355-1363.

[8] Hans C, Dobra A, West M. Shotgun Stochastic Search for "Large p" Regression, *Journal of the American Statistical Association*, 102(478):507-516, 2007.

[9] Haseman JK, Elston RC. The investigation of linkage between a quantitative trait and a marker locus. *Behav Genet* 2:3–19, 1972.

[10] Kang, H. M., Zaitlen, N. A., Wade, C. M., Kirby, A., Heckerman, D., Daly, M. J., Eskin, E. (2008). Efficient control of population structure in model organism association mapping. Genetics, 178(3), 1709–1723.

[11] Lippert, C., Listgarten, J., Liu, Y., Kadie, C.M., Davidson, R.I., Heckerman, D. (2011) FaST linear mixed models for genome-wide association studies. Nature methods, 8, 833–835.

[12] Pazokitoroudi A, Wu Y, Burch KS, Hou K, Zhou A, Pasaniuc B, Sankararaman S. Efficient variance components analysis across millions of genomes. *Nat Commun* 11:4020, 2020.

[13] Wang G, Sarkar A, Carbonetto P, Stephens M. A simple new approach to variable selection in regression, with application to genetic fine mapping. *J. R. Stat. Soc. B*, 82:1273-1300, 2020.

[14] Zhou, X., Stephens, M. (2012) Genome-wide efficient mixed-model analysis for association studies. Nature genetics, 44, 821–824.
